# Mechanistic inferences from analysis of measurements of protein phase transitions in live cells

**DOI:** 10.1101/2020.11.04.369017

**Authors:** Ammon E. Posey, Kiersten M. Ruff, Jared M. Lalmansingh, Tejbir S. Kandola, Jeffrey J. Lange, Randal Halfmann, Rohit V. Pappu

## Abstract

The combination of phase separation and disorder-to-order transitions can give rise to ordered, semi-crystalline fibrillar assemblies that underlie prion phenomena namely, the non-Mendelian transfer of information across cells. Recently, a method known as Distributed Amphifluoric Förster Resonance Energy Transfer (DAmFRET) was developed to study the convolution of phase separation and disorder-to-order transitions in live cells. In this assay, a protein of interest is expressed to a broad range of concentrations and the acquisition of local density and order, measured by changes in FRET, is used to map phase transitions for different proteins. The high-throughput nature of this assay affords the promise of uncovering sequence-to-phase behavior relationships in live cells. Here, we report the development of a supervised method to obtain automated and accurate classifications of phase transitions quantified using the DAmFRET assay. Systems that we classify as undergoing two-state discontinuous transitions are consistent with prion-like behaviors, although the converse is not always true. We uncover well-established and surprising new sequence features that contribute to two-state phase behavior of prion-like domains. Additionally, our method enables quantitative, comparative assessments of sequence-specific driving forces for phase transitions in live cells. Finally, we demonstrate that a modest augmentation of DAmFRET measurements, specifically time-dependent protein expression profiles, can allow one to apply classical nucleation theory to extract sequence-specific lower bounds on the probability of nucleating ordered assemblies. Taken together, our approaches lead to a useful analysis pipeline that enables the extraction of mechanistic inferences regarding phase transitions in live cells.

## Introduction

Phase transitions can lead to the formation of various types of macromolecular assemblies in cells [1–7]. These include liquid or gel-like biomolecular condensates that concentrate protein and nucleic acid molecules [8–10], liquid crystalline assemblies [11], and semi-crystalline assemblies [12] such as prions [13, 14], which are protein-based elements that enable non-Mendelian inheritance [15–17]. Phase transitions are characterized by cooperative changes to order parameters and different types of transitions are associated with changes to different types of order parameters [18]. Phase separation is a form of phase transition where macromolecular concentration is the relevant order parameter [6, 18, 19]. In a macromolecular solution, if and only if homotypic interactions are the main drivers of phase separation [20, 21], then the system separates into two or more coexisting phases when the concentration of macromolecules crosses a system-specific threshold concentration, with each pair of phases being delineated by a distinct phase boundary.

On a microscopic level, phases are defined and distinguished on the basis of structural symmetry and the type of order found in a phase [18]. Distinguishing order versus disorder in different phases requires quantification of how a phase responds to a set of symmetry operations such as the Euclidean group [22], which represents a set of translational, rotational, and reflection operations. Isotropic liquids and gases are statistically invariant under all symmetry operations. Accordingly, the entire Euclidean group is the symmetry group for isotropic fluids and such systems have the highest possible symmetry. As a result, a vapor-liquid transition of isotropic fluids does not involve a change in symmetry, but it does involve a change in density. The working hypothesis that has emerged from *in vitro* characterizations is that liquid-liquid phase separation that gives rise to some biomolecular condensates [1] is akin to a vapor-liquid transition that involves a change in density [23], without a change in symmetry [24]. There can be significant differences in compositions [25–27], including the exclusion of certain components such as macromolecular crowders from dense phases [28] points to a change in compositional symmetries.

The onset of order in a phase implies that the phase in question is statistically invariant to a subgroup of operations that define the Euclidean group. The extent and type of order versus disorder is governed by the size of the symmetry group for a given phase and the type of symmetry operations to which the phase in question remains statistically invariant. At equilibrium, higher entropy disordered phases have high symmetries whereas lower entropy ordered phases have lower symmetries. Accordingly, disorder-to-order transitions are also described as symmetry-breaking transitions [18]. Symmetries are quantified in terms of non-conserved order parameters, typically taking values between 0 (maximal disorder) and 1 (maximal order). This is relevant because phase separation *in vitro* and in live cells can also be accompanied by the breaking of symmetries that drive collective disorder-to-order transitions [19, 29, 30]. Examples include protein crystallization *in vitro* [31] and the formation of fibrillar solids both *in vitro* and in cells [18, 19, 29, 30].

The order parameter for phase transitions that combine phase separation and disorder-to-order transitions has two components namely, macromolecular concentration and a measure of order / disorder [32, 33]. The free energy barrier for nucleating an ordered phase is determined in part by the supersaturation [34–36], defined as the natural logarithm of the ratio of the bulk concentration to the saturation concentration [37]. Above the saturation concentration, the supersaturation increases, thereby decreasing the nucleation barrier and increasing the probability of spontaneously nucleating the ordered assembly [38] (**Figure 1a**). The assembly of prion-like states arises through templated growth of the nucleus, which is the embryo of the ordered phase that forms within the disordered phase [14]. The latter is either a dilute or dense liquid where the distinction between dilute and dense points to the difference in macromolecular concentration within the liquid phase, and the extent of order versus disorder refers to the fraction of molecules that are incorporated into the ordered phase.

**Figure 1:**
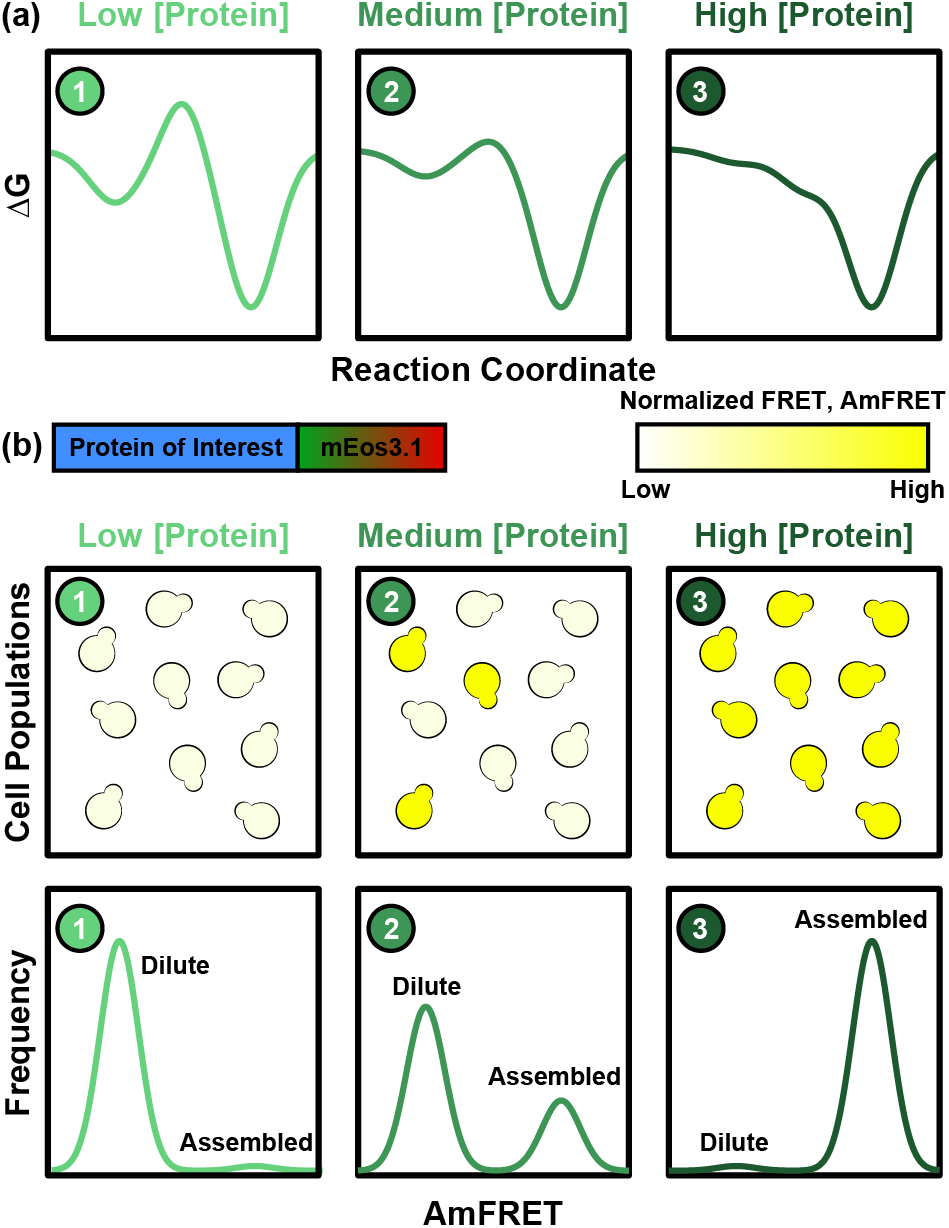
Schematic of disorder-order transitions and details of the DAmFRET assay. (a) At low (1) and medium (2) protein concentrations there is a free energy barrier for nucleation of ordered assemblies, whereas at high (3) protein concentrations this barrier is eliminated. Here, the protein concentration in (1) corresponds to a system that is subsaturated and protein concentrations in (2) and (3), respectively correspond to systems that are supersaturated. (b) DAmFRET is performed using a chimera of an aggregation-prone or prion-like protein of interest and a photoconvertible fluorescent protein mEoS3.1. This allows for the examination of FRET in live cells and AmFRET can be measured using flow cytometry for thousands of cells with different expression levels. We assume that steady-state assembly is reached instantaneously upon nucleation, at least given the temporal resolution of measurements and the rapid timescales one is likely to associate with actual barrier crossings, i.e., *transition path dynamics* in typical physico-chemical reactions [46]. At low concentrations, the system is subsaturated, and nucleation is highly disfavored so all cells should have low AmFRET values indicating the protein remains in the dilute phase (off-white color). At intermediate concentrations, the system is supersaturated and the barrier for nucleation is reduced and now some cells maintain a dilute population and low AmFRET, whereas other cells undergo nucleation and thus the formation of ordered assembles as indicated by high AmFRET (bright yellow). At high concentrations the system is significantly supersaturated, and there is no longer a barrier for nucleation and most cells show high AmFRET.

Results of measurements made under conditions where the effects of active processes are minimized are helpful for understanding how the milieu of a living cell impacts the intrinsic driving forces for phase transitions [39]. Live cell investigations of macromolecular phase transitions have been driven by adaptations of optogenetics technologies [40, 41], advances in super-resolution microscopy [34, 42], and single particle tracking [43, 44]. Recently, a new method known as Distributed Amphifluoric Förster Resonance Energy Transfer (DAmFRET) was introduced to investigate phase transitions that lead to prion-like assemblies in yeast [39] and mammalian cells [45]. In DAmFRET measurements, live cells are used as femto-liter sized test tubes in which protein self-assembly is measured. FRET is used as the reporter for protein assembly within each cell. Because changes to FRET intensities result mainly from changes to intermolecular distance, DAmFRET measures changes to both density and the extent of order / disorder. Accordingly, phase transitions such as prion formation that combine phase separation and disorder-order transitions can be measured using DAmFRET.

In DAmFRET experiments a photo-switchable fluorescent protein mEos3.1 is expressed as a chimera with the protein of interest (**Figure 1**) [39]. mEos3.1 is a green fluorescent protein that can be converted to a red fluorescent protein upon illumination with violet light. This conversion can be achieved in a controlled, time- and intensity-dependent manner. Photo conversion allows the generation of FRET pairs from a single genetic construct in a consistent and controllable ratio that is independent of expression level. The resulting green and red forms constitute the FRET donor and acceptor, respectively. In yeast, variability in expression under the *GAL1* promoter and the 2 microns replication system is achieved by harnessing the wide range (more than 100-fold) of plasmid copy numbers across thousands of cells. This allows wide coverage of protein expression levels across a population of cells analyzed by flow cytometry. Intracellular phase behavior is interrogated after cell division and protein degradation have been turned off upon galactose induction. This ensures that each cell mimics a closed system and that phase transitions are being interrogated in a cell autonomous manner since propagation via cell division is eliminated. In effect, the DAmFRET approach affords two advantages: from the perspective of physical chemistry, it allows one to treat each live cell as being a femtoliter test tube. Secondly, live cells are not passive reservoirs and therefore the insights gleaned from the DAmFRET assay are likely to have a direct bearing on understanding how spontaneous and driven phase transitions are controlled by protein expression and influenced by a dynamic cellular milieu.

DAmFRET assays enable the interrogation of protein assembly behavior across a large population of live cells by monitoring AmFRET, defined as the ratio of sensitized emission FRET to acceptor fluorescence intensity [39]. AmFRET measured by flow cytometry is not an order parameter in the strictest sense of the term. For example, two different proteins that have similar AmFRET values do not necessarily have the same degree of order / disorder. Likewise, the densities of assemblies corresponding to similar AmFRET values do not have to be similar. However, it can be used as a proxy of a non-conserved parameter for understanding the nature of the phase transition that a specific system undergoes because, for a specific system, interrogated across a population of cells, it provides a quantitative assessment of relative order versus disorder. Further, as we show here, parameters extracted from analysis of DAmFRET histograms can be compared across different proteins. Dispersed monomers should have low AmFRET values around zero, whereas the AmFRET value will increase upon assembly formation. For proteins that do not become trapped in long-lived partially assembled intermediates, we can distinguish the concentration dependence of assemblies into three categories. At low overall expression levels, the effective concentration is below the saturation concentration, and a dilute, disordered phase will form for the protein of interest across the population of cells (**Figure 1b**). At intermediate levels of expression, the protein of interest is supersaturated, and two distinct populations of cells can coexist with one another: In one population, the protein of interest will be entirely in the dilute, disordered phase, whereas in the coexisting population, we will observe proteins concentrated into dense, ordered assemblies. The likelihood of observing a coexisting population of cells that feature dilute, disordered phases will be tied to the supersaturation-dependent nucleation probability. At high expression levels, dense ordered assemblies will form for the protein of interest in a majority of the cells. In each of these three categories cells only fall into one of the two distinct states, dilute or ordered assembled; therefore, these proteins can be classified as undergoing a two-state discontinuous phase transition.

Given that the DAmFRET assay collects data across expression levels that span 2-3 orders of magnitude [39], one can analyze two-dimensional histograms of expression levels and AmFRET values across a large (~10^4^) population of cells and categorize proteins based on the types of assembly behaviors observed. Information extracted from analysis of two-dimensional histograms allows us to quantify the expression level at which proteins in 50% of the cells are in the assembled phase. Because the driving forces for phase transitions are dependent on protein concentration through the degree of supersaturation, the measure of the expression level at which proteins in 50% of the cells are in the assembled phase allows for a comparative assessment of sequence-encoded driving forces for phase transitions that show two-state discontinuous behavior. Further, one can also analyze the measured phase behavior to identify proteins into groups that do not undergo a measurable transition or undergo continuous or discontinuous transitions into different phases that are amenable to interrogation by the DAmFRET assay.

Here, we assess and analyze the information gleaned from DAmFRET measurements to answer three questions: (1) Given the large amount of data that can be generated through the DAmFRET assay, approximately 10^3^ histograms in a day, we ask if we can automate the classification of phase transitions measured by DAmFRET into distinct categories? (2) Further, do the phase transitions of proteins that form *bona fide* prions belong to a specific category? And (3), for proteins that show discontinuous two-state phase behavior, consistent with nucleated transitions, can we use data from DAmFRET experiments to extract information regarding the barriers to nucleation at different degrees of supersaturation? To answer the first question, we develop a supervised method [47] that classifies different types of DAmFRET histograms based on analyzing synthetic AmFRET data as a function of expression level. We apply these methods to analyze DAmFRET data obtained for a large number of candidate prion-like domains (cPrDs) [16] and uncover sequence features that correlate with the degree of discontinuity of a two-state transition. We also show that expression levels at which 50% of the cells are in the assembled state can be accurately extracted by analysis of DAmFRET measurements using our automated method. This value, designated as *c*_50_, can be used to rank order proteins based on their driving forces for assembly. As for the second question, we find that proteins classified as undergoing two-state discontinuous transitions tend to show prion-like properties, although *bona fide* prions can be classified into other categories of transition classes as well. Finally, we deploy a numerical method motivated by classical nucleation theory to show that information from DAmFRET measurements can be adapted to estimate the sizes of free energy barriers to nucleation of ordered assemblies. Usage of this analysis does require augmentation to the data collected from DAmFRET measurements. Overall, the analysis pipeline we develop here paves the way for gleaning quantitative and mechanistic inferences from analysis of large-scale, proteome-level investigations of prion formation and related phase transitions in live cells.

## Results

### Classifiers of phase transitions measured using DAmFRET

Depending on how AmFRET changes as a function of protein expression levels, we classify DAmFRET datasets into five categories.

### One-state – no transition

Across the concentration range interrogated by DAmFRET experiments, AmFRET does not change its value. If AmFRET is low across the concentration range, then this is a manifestation of *one-state behavior without assembly*. Alternatively, if AmFRET is uniformly high across the concentration range, then the data are concordant with the designation of *one-state behavior with assembly*.

### Two-state discontinuous transitions

These are characterized by the existence of a threshold expression value, such that above this value AmFRET changes in a stepwise fashion from low to high values. Accordingly, cells fall into one of two categories defined by the presence or absence of dense, ordered assemblies.

### Two-state continuous transitions

Here, AmFRET increases continuously once the expression level crosses a threshold value. Unlike two-state discontinuous transitions, the two-state continuous transitions are best modeled using a smooth function that interpolates between low and high AmFRET values. Continuous transitions likely reflect the formation of a distribution of low-affinity oligomers or mesophases whereby the sizes and / or numbers of assemblies increase as concentration increases. This would result in a continuous increase in FRET intensity, which is also proportional to the numbers of complexes of specific sizes.

### Three- or multistate transitions

Here, the transitions proceed through multiple states characterized by intermediate values of AmFRET. It is worth noting that intermediate values of AmFRET may also be a reflection of non-equilibrium steady states or long-lived transients.

### Enabling supervised learning and classification based on synthetic data sets for each of the different categories of phase transitions

Synthetic two-dimensional histograms of expression level and AmFRET were generated for each of the five categories of phase transitions (**Figure 2a**). We generate synthetic AmFRET datasets using a simple step function or a sigmoid function. The minimum and maximum values of the step and sigmoid functions in the synthetic data for the supervised learning are set as A_min_ = 0 and A_max_ = 1.5. Each histogram was generated using ~10^4^ data points, where the points are proxies for measurements in individual cells. Points were generated using a log10(expression) value in the range of 2.0 to 8.0 and an AmFRET value between 0.0 and 1.5. Noise was added in the log10(expression) value using a Gaussian with a mean of 0.5 and standard deviation of 0.75, which we term *ce*. The expression at which 50% of the cells show high AmFRET assemblies is designated as *c*_50_ and log_10_(*c*_50_) was set to be 1.0, 2.0, 4.0, 6.0, 8.0, 10.0, or 12.0; to increase the width of the range of concentrations where high and low AmFRET values overlap, we introduced random noise to *c*_50_ with a variance of ± 0.8, which we term *c*_50e_. For three-state transitions, the variance value was increased to ± 4.0 in order to represent observations in some instances of experimental data where a wider range of concentrations have both low and high AmFRET populations, and to test the ability of our algorithm to classify data with larger overlaps. For our analysis, we only consider data points with log_10_(expression) values between 3.0 and 10.0. Note that datasets with log_10_(*c*_50_) less than 3.0 or greater than 10.0 were intentionally chosen to be outside this concentration range. Accordingly, datasets generated using log_10_(*c*_50_) = 1.0 or 2.0 were designated as “*One state: with assembly*”, and datasets with log_10_(*c*_50_) = 12.0 were designated as “*One state: No assembly*”. Data for log_10_(*c*_50_) = 10 fall at the edge of the range of generated points. However, because of noise, data generated using this model show evidence for transitions from low AmFRET to high AmFRET states, although this transition is incomplete within the simulated concentration range. These synthetic data were used to represent experimentally observed cases where the complete transition is not fully captured.

**Figure 2:**
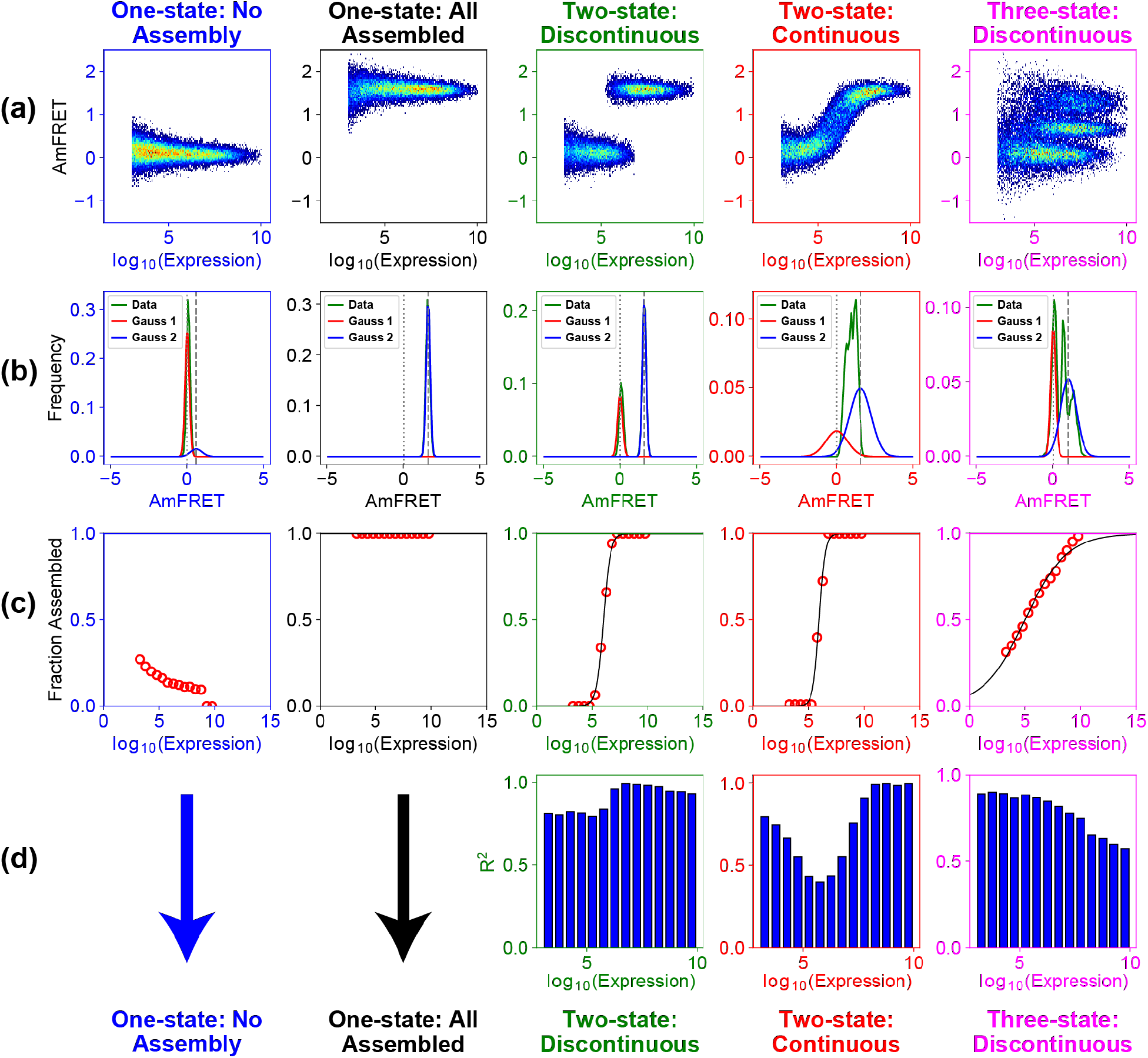
Schematic of the method used to classify synthetic DAmFRET data. (a) Row one shows representative synthetic DAmFRET histograms for the five different classes we aim to classify. (b) Each synthetic DAmFRET histogram is sliced into expression level windows and the corresponding 1-dimensional AmFRET histograms are fit to a sum of two Gaussians where the position of the first peak is set to AmFRET=0 and the position of the second peak is set to the maximum mean AmFRET across all expression slices after removal of AmFRET values around zero. Here, example fits are shown for the expression level slice of 6-6.5. For the other expression slices see **Figure S1**. (c) The fits are used to extract the fraction of cells that undergo assembly in each expression slice by taking the area under the curve of the second peak and dividing it by the total area under both curves. If the change in fraction assembled from the last expression slice minus the minimum fraction assembled is less than 0.1, then the profile is classified as one state and the mean AmFRET across all expression slices determines whether the protein is in the no assembly versus all assembled class. (d) For all profiles that are not classified as one-state, we use the *R*^2^ values of the fit to the sum of two Gaussians across the expression slices to rule out histograms that do not show two-state discontinuous behavior. Profiles that show a minimum in *R*^2^ values around log_10_(*c*_50_) are classified as two-state continuous and profiles that show a continuous decrease in *R*^2^ values across expression slices are classified as three-state discontinuous. All other profiles pass the test of the null hypothesis and are thus classified as two-state discontinuous.

To mimic data from measurements that correspond to two-state transitions, the synthetic data for AmFRET values were generated using a piecewise step function with log_10_(*c*_50_) as the transition point to mimic two-state discontinuous transitions, or using a Boltzmann sigmoid logistic equation [48] to mimic continuous two-state transitions. For the latter case, data were generated by sampling points from the following equation for AmFRET, which will be a function of the bulk concentration *c* and the values will depend on the baselines for A_max_ and A_min_, our choices for *c*_50_ and the choice of *m*, which is the slope of the transition between low and high AmFRET values. Accordingly, for synthetic continuous two-state data, AmFRET as a function of *c* is computed using:

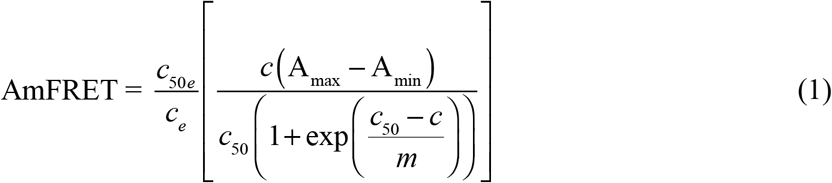

Synthetic data to mimic three-state transitions were generated with two overlapping piecewise step functions, one stepping from zero to 0.4A_max_ and the other to 0.8A_max_. For a given set of parameters namely, (i) the values of *c*_50_, (ii) stepwise versus sigmoid function, and (iii) whether the transitions are one-, two- or three-state, synthetic data were generated in three independent repetitions using different random number seeds with three different AmFRET noise levels (low, medium, high).

### Heuristics to classify different categories of phase transitions

Since synthetic datasets were generated from known priors, we can assess the accuracy of a classifier with certainty. This supervised approach, whereby we know how data were generated and ask if a classifier is accurate in its assignment of the category of phase transitions, yields heuristics that can be deployed in the analysis of real data while also knowing the level of confidence one can ascribe to a specific classification.

The null hypothesis is that all synthetic DAmFRET histograms belong to the two-state discontinuous transition category. We use the synthetic DAmFRET histograms to rule in or rule out this hypothesis. Specifically, each histogram is divided along the expression-axis into expression level slices. The one-dimensional histogram of AmFRET values within each expression level slice is fit to a sum of two Gaussians (**Figure 2b** and **Figure S1**). Here, the position of the first peak is set to be at zero AmFRET and the position of the second peak is set to the maximum mean AmFRET across expression level slices. For calculating the latter, we discard all AmFRET values within the noise around AmFRET=0. We use a sum of two Gaussians since the null model is that of a discontinuous two-state transition wherein all cells either have no assembly or have reached steady-state assembly. Next, the fraction assembled in each expression slice is calculated by computing the area under the curve of the second Gaussian peak and dividing it by the total area under both curves (**Figure 2c**).

If the fraction assembled in the last expression slice minus the minimum value for the fraction assembled is less than 0.1, then the DAmFRET histogram is classified as being one-state. The mean AmFRET value is then used to determine if the one-state behavior corresponds to no assembly or all assembled in the given expression range. Specifically, if the mean AmFRET value is less than the noise cutoff around AmFRET=0 it is classified as being one-state without assembly.

For DAmFRET histograms that are inconsistent with one-state assembly we examine the *R*^2^ values as a function of expression slice for the fit of the AmFRET histogram to the sum of the two Gaussians (**Figure 2d**). For discontinuous two-state transitions, the *R*^2^ values should be high (near 1.0) across all expression level slices. Additionally, Gaussian fits of AmFRET histograms for two-state continuous and three-state discontinuous data show distinct *R*^2^ dependencies on expression level. Specifically, for two-state continuous transitions, the *R*^2^ values are lowest at expression level slices around log_10_(*c*_50_). This is because, within the transition region, most cells have AmFRET values that lie between zero and high AmFRET for a two-state continuous transition (**Figure 2b** and **Figure S1**). The percent of cells that lie between the limits will depend on the slope of the transition. At equivalent noise levels, smaller slopes imply that a larger percentage of cells lie between the limits, thereby leading to a smaller *R*^2^ value. In contrast, the three-state discontinuous transitions show a linear decrease in *R*^2^ values across expression level slices due to the existence of three overlapping states. We used these two trends to determine if a DAmFRET histogram is consistent with either a two-state continuous or multi-state transition (see Materials and Methods for details). If *R*^2^ as a function of expression slice does not show either of these two trends, then it passes our null hypothesis and the transition measured by DAmFRET is classified as being two-state discontinuous.

**Figure 3** shows how well the classifier performs in categorizing the synthetic data. The corresponding synthetic DAmFRET histograms are shown in **Figure S2**. Each dataset is assigned a color that corresponds to its classification and the shade of the color indicates the confidence level in the classification, the darker the shade the more confidence in the classification. Synthetic data were created in three replicates and therefore each row represents a different replica. Of the 162 histograms that we generated, our classification scheme yields a 90% accuracy, classifying 146 of the histograms correctly. In general, we succeed in classifying synthetic datasets into the correct categories, although the confidence in these classifications decreases as log_10_(*c*_50_) approaches the limits of the expression range that can be assessed, or the noise becomes too high. This is evident for synthetic data corresponding to two-state continuous transitions (**Figure 3d**).

**Figure 3:**
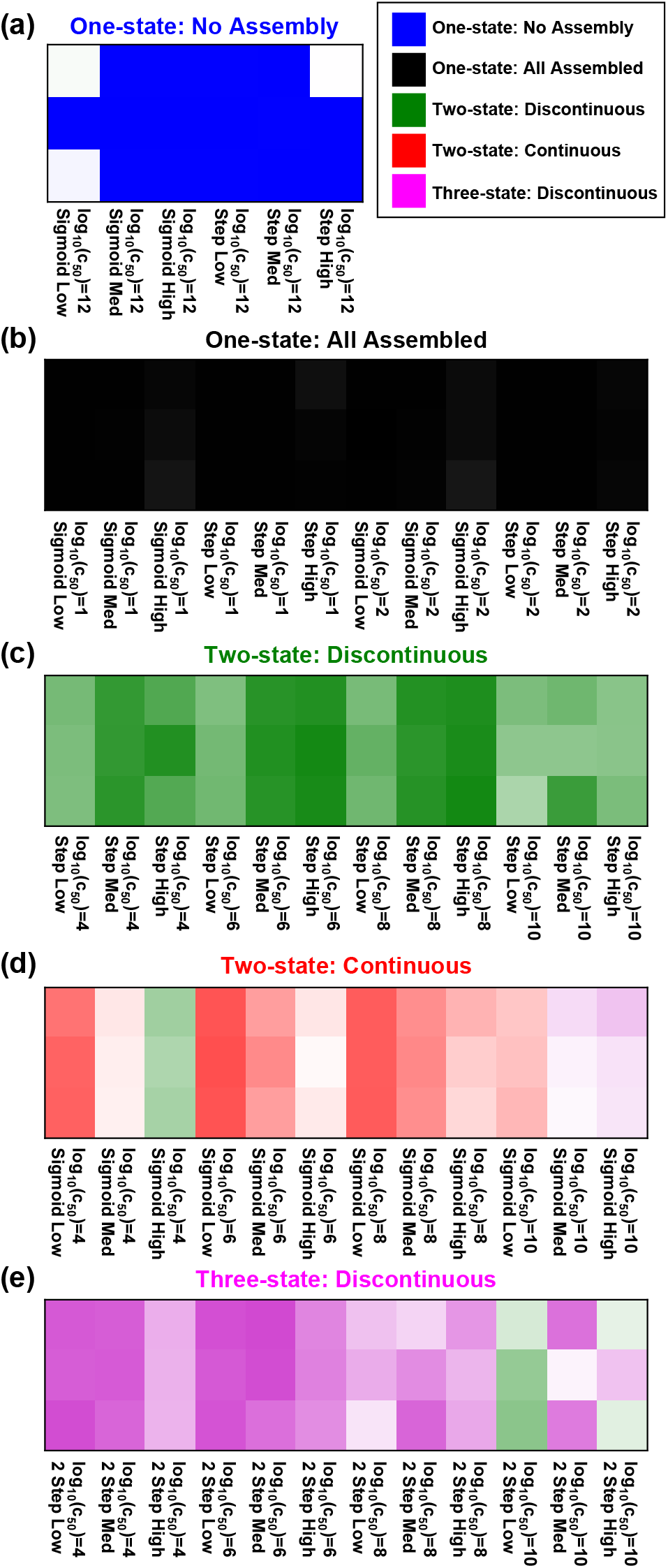
Classification of synthetic DAmFRET histograms. Each histogram is colored by its classification and the shade of the color indicates the confidence of that classification. The darker the shade the more confident the method is in its classification (**Table S1**). For a non-shaded version of **Figure 3** see **Figure S3**. Each row denotes a different replica. Each column is denoted by the log_10_(*c*_50_) (1.0, 2.0, 4.0, 6.0, 8.0, 10.0, or 12.0), the function (Sigmoid, Step, or 2 Step), and the noise level (Low, Medium, or High) used to generate the synthetic DAmFRET histograms.

Beyond classifying datasets into distinct categories, for data classified as two-state systems we can fit the fraction assembled as a function of expression to a logistic function and extract *c*_50_. This is a strategy we adopt for analyzing real DAmFRET data, and we prototype it here for synthetic data. **Figure 4** shows the comparison between the actual value of *c*_50_ used to generate the synthetic datasets and the estimate for *c*_50_ extracted from the logistic fits to the synthetic datasets. We observe good agreement between the actual versus extracted values. However, deviations from the actual value of *c*_50_ occurs when this value approaches the limits of the expression range that can be accessed. Overall, the results in **Figures 3** and **4** show that the method introduced here allows for accurate classifications of phase transitions, while also enabling the extraction of accurate quantitative estimates for *c*_50_.

**Figure 4:**
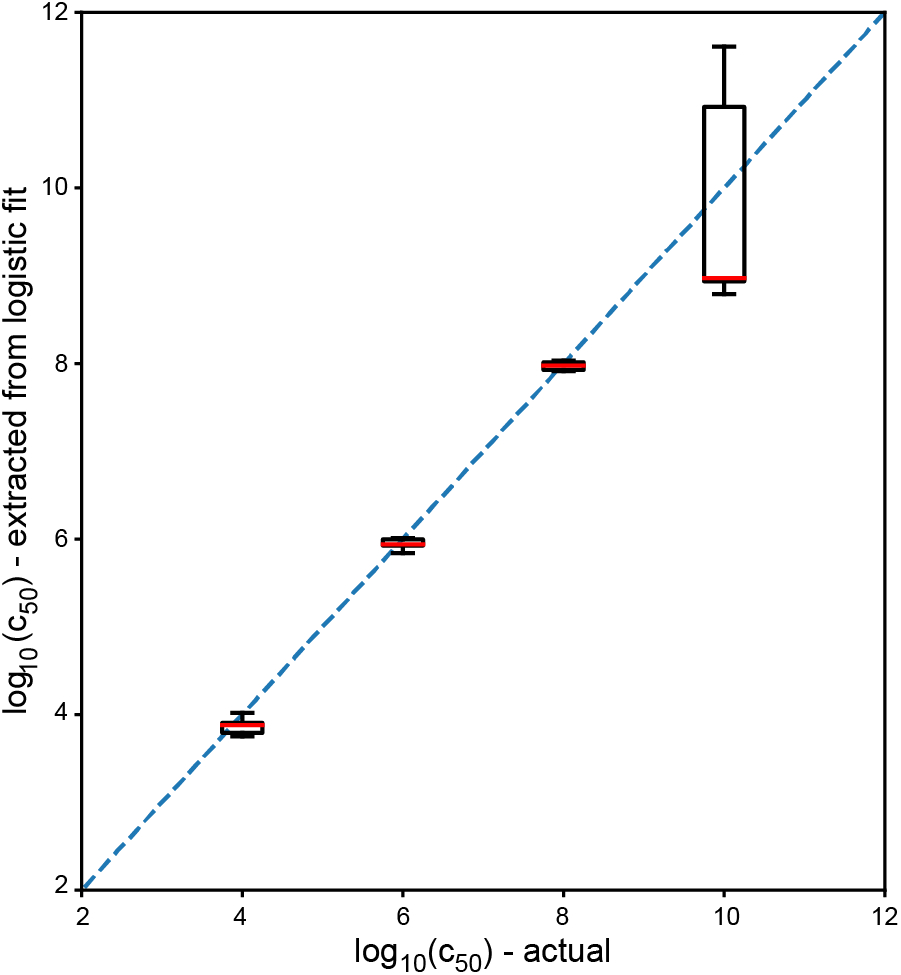
Actual log_10_(*c*_50_) values versus the log_10_*c*_50_) values extracted from the logistic fit of the fraction assembled profiles for all two-state DAmFRET histograms (**Figure 2c**). Boxplots show distributions of log_10_(*c*_50_) extracted from 54 histograms for log_10_(*c*_50_)=4.0, 6.0, and 8.0 and 38 histograms for log_10_(*c*_50_)=10.0.

### Application of the numerical classifier to DAmFRET data for a set of cPrDs

We collected and analyzed DAmFRET data for 84 of the 100 candidate prion domains (cPrDs) identified and analyzed by Alberti et al., [16] in their screen for proteins with prion behavior (see Materials and Methods). Alberti et al. used a combination of four different assays to test if the cPrDs are *bona fide* prion formers. In their assessments, a *bona fide* prion former should have the following characteristics: (a) it should form foci in cells; (b) the assemblies extracted from cells should be stable as assessed by sensitivity to detergent; (c) the assembly state should be transferable from mother to daughter cells during cell division and (d) the assemblies should stain positively with amyloid sensitive dyes such as thioflavin T (ThT). In all, Alberti et al., identified 18 cPrDs that showed prion-like behavior in all four assays [16]. Here, we make comparisons against the assessments made by Alberti et al. in order to test our classification method and to uncover information about mechanisms for prion formation as assessed by DAmFRET.

**Figure 5** shows our classification of the cPrDs for which DAmFRET measurements were performed. The corresponding DAmFRET histograms are shown in **Figure S4**. For these data, we included an additional class to allow for proteins that are not classified as one-state, but only show assembly in fewer than 10% percent of the cells that were interrogated. Given the low percent of cells in the assembled state there is not enough information to classify the type of transition observed and therefore we denote this class as “*Infrequent Transition*”. Additionally, unlike the synthetic data set, in which the transition was binary in terms of being discontinuous or continuous, the classification of real data for two-state histograms falls along a continuum from discontinuous to continuous (**Figure 5e**). Thus, we group all cPrDs that show two-state behavior together and then further annotate their position on the spectrum.

**Figure 5:**
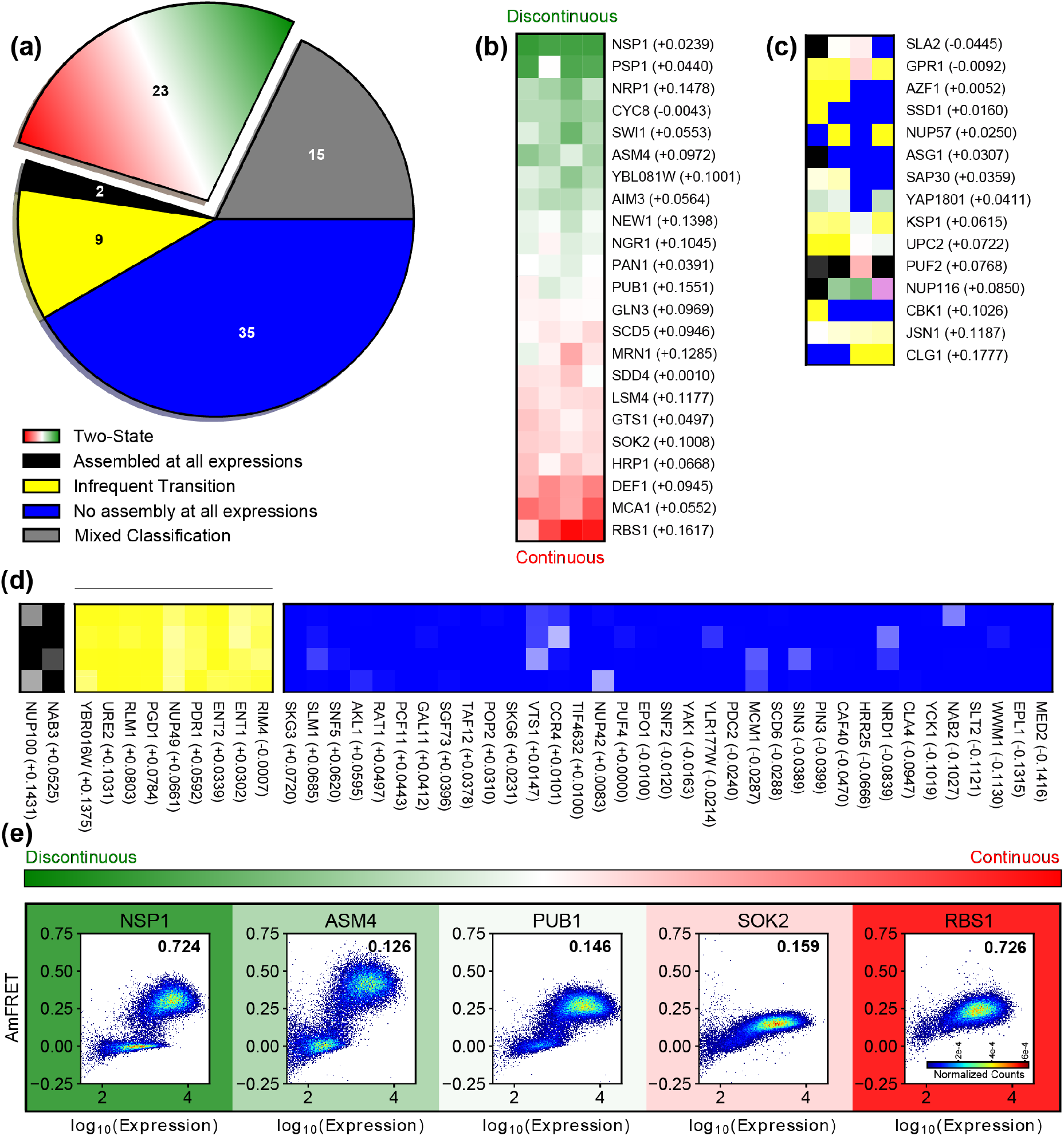
Classification of DAmFRET histograms of 84 cPrDs previously examined by Alberti et al. (a) Number of cPrDs in each type of phase transition class. (B-D) List of cPrDs that were classified as two-state (b), mixed classification-having replicates belonging to multiple classes (c), assembled at all expression values (d-black), undergoing an infrequent transition (d-yellow), and showing no assembly at all expression values (d-blue). For the checkboard plots, each square represents an experimental replicate, and the color of the square denotes the classification. The darker the shade the more confident the method is in the classification (**Table S2**). The classifications not shaded by confidence score are showed in **Figure S5**. The PAPA prion score for each cPrD is listed in parentheses. The cPrDs classified as undergoing a two-state transition are sorted by the degree of discontinuity of the transition. The cPrDs for all other classes are sorted by their PAPA prion score [50]. (e) Representative DAmFRET histograms of cPrDs that were classified as undergoing a two-state transition. The color corresponding to each histogram denotes the degree of discontinuity in the transition (see Materials and Methods).

**Figure 5a** shows that of the 84 cPrDs that were examined using DAmFRET, 35 showed no assembly across all expression levels, 9 showed infrequent transitions, 23 showed two-state behavior, and 2 showed assemblies at all expression levels. Additionally, 15 cPrDs showed mixed classification in which not all of the individual cPrD replicates were sorted into the same class (**Figure 5c**). **Figures 5b-d** show the classification of each cPrD for each of its four replicates. The pixel color denotes the classification, and the darkness indicates the confidence level in that classification, with darker being more confident (Materials and Methods). The two-state cPrDs are sorted by where they fall on the spectrum of discontinuous to continuous transitions (**Figure 5b**, Materials and Methods).

**Figure 5e** shows representative DAmFRET histograms for five cPrDs that show two-state behavior in the DAmFRET assays. Here, the color represents where the cPrD of interest falls on the two-state spectrum. Nsp1 shows discontinuous two-state behavior with low AmFRET up until a threshold expression level and then a transition to a high AmFRET state with few cells showing intermediate AmFRET values. In contrast, Rbs1 features continuously increasing AmFRET values as the expression level increases. The other two-state cPrDs fall between these two extremes. Along the progression from discontinuous to increasingly continuous transitions we observe a positive slope in AmFRET at low expression values and a larger population of cells with intermediate AmFRET values in the transition region. In this context, it is worth noting that Pub1, which falls in the middle of the spectrum between discontinuous and continuous transitions, has been shown to form both disordered liquid-like condensates and ordered amyloid-like assemblies [49]. This may suggest that the positive slope in AmFRET at low expression values corresponds to the formation of liquid-like condensates, which, with increasing expression, transform into ordered assemblies.

cPrDs that fall outside the classification as two-state are grouped by classification and ordered by their predicted prion aggregation propensity as quantified by PAPA (**Figure 5c-d**). Toombs et al.,[50] previously showed that a cutoff score of +0.05 yielded greater than 90% accuracy in delineating the 18 *bona fide* prion domains from the 18 non-prion domains in the Alberti et al. study. We find that 31 of the 35 cPrDs that were classified as showing no assembly across all expression levels had PAPA scores below this cutoff. In comparison, 3 of 9 cPrDs with infrequent transition (33%), 6 of 23 two-state cPrDs (26%), and 0 of 2 cPrDs that assembled at all expression levels (0%) had PAPA scores below this cutoff. Thus, our classification method is consistent with the trends expected based on PAPA scores.

### cPrDs that are classified as two-state discontinuous from DAmFRET show prion-like behavior

By cross-referencing our classifications with the results of Alberti et al., [16] we tested our hypothesis that cPrDs that are classified as undergoing two-state discontinuous transitions in the DAmFRET assay are in fact *bona fide* prion forming domains. Eight of the cPrDs were classified as undergoing a two-state discontinuous transition for all four experimental replicates. Therefore, we examined how many of these eight cPrDs engendered prion-like behavior in each of the four assays conducted by Alberti et al. (**Figure 6a-d**). All of the cPrDs that were classified as undergoing a two-state discontinuous transition formed foci in cells (**Figure 6a**), had assemblies that were SDS resistant (**Figure 6b**), and stained positively with ThT (**Figure 6d**). Two of the eight cPrDs did not pass the most stringent test for prion-like behavior, namely a heritable switch in the context of a Sup35 chimera (**Figure 6c**). One of these, Cyc8 was subsequently shown to be a *bona fide* prion [51]. These results suggest that, while a two-state discontinuous transition likely implies amyloid assembly consistent with prion formation, it does not necessitate heritability. This is expected given that DAmFRET interrogates the mechanism of assembly formation, whereas the biological context of prions dictates whether the amyloids propagate or not. For example, propagation in rapidly dividing budding yeast cells requires specific interactions with the yeast-specific prion replication factor Hsp104 [52].

**Figure 6:**
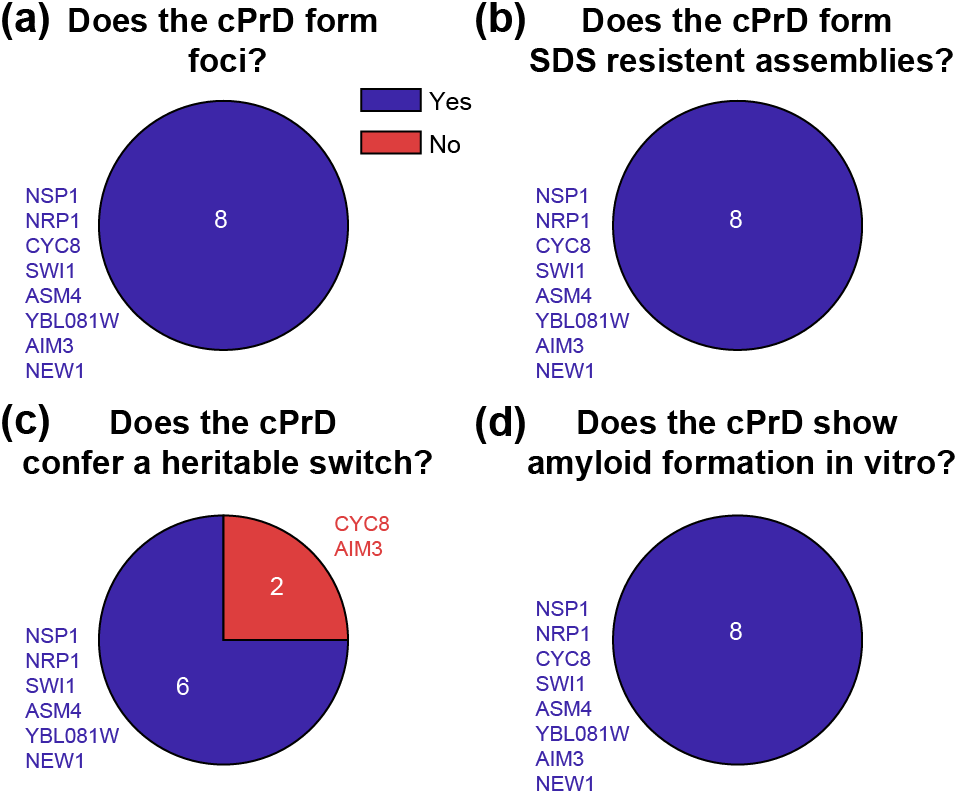
Assessment of cPrDs classified as undergoing a discontinuous two-state transition. For the eight DAmFRET profiles that were classified as two-state discontinuous we examined whether or not the associated cPrDs formed (a) foci, (b) SDS resistant assemblies, (c) heritable assemblies, and (d) ThT positive assemblies in the assays conducted by Alberti et al. [16].

**Figure 7:**
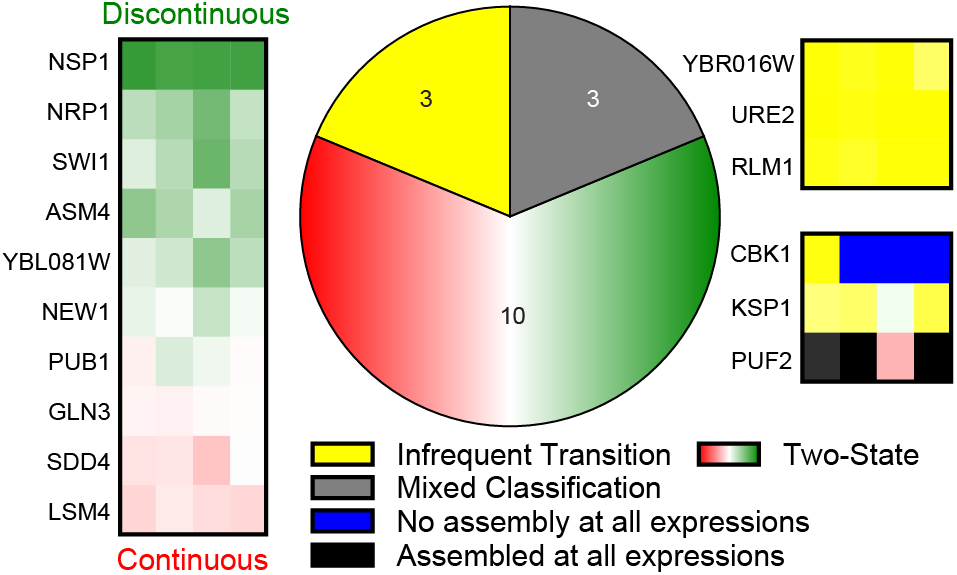
Classification of DAmFRET histograms of the 16 *bona fide* prion-forming domains identified by Alberti et al. and analyzed with DAmFRET. For the checkboard plots, each rectangle represents an experimental replicate of one of the prion domains and the color of the rectangle denotes the classification. The darkness of the shade denotes the degree of confidence in that classification, with darker shades corresponding to more confidence. Those prion domains classified as two-state are sorted by the degree of discontinuity at the transition.

### DAmFRET histograms of cPrDs that are bona fide prions are not always classified as undergoing a two-state discontinuous transition

Next, we asked whether being a *bona fide* prion implies that the cPrD is classified as undergoing a two-state discontinuous transition from the DAmFRET analysis. Of the 18 *bona fide* prion-forming domains, 16 were examined using DAmFRET. Of these, ten were classified as two-state, with six of these *bona fide* prion-forming domains being classified as two-state discontinuous in all four of their replicates (**Figure 6**). Additionally, three of the *bona fide* prion-forming domains were classified as undergoing an infrequent transition and three showed mixed classification. Given that ten of the 16 *bona fide* prion-forming domains were not classified as undergoing a two-state discontinuous transition, this suggests prions can exhibit other types of DAmFRET histograms. We find that 15 of the 16 did show at least the emergence of a transition to a higher AmFRET state. Only Cbk1 had a negligible increase in AmFRET across the expression levels analyzed. Together with the fact that Cbk1 was only weakly amyloidogenic in Alberti et al., [16] this suggests that its self-propagating ability and / or nucleation mechanism may be atypical of prions.

In general, our analysis indicates that not all proteins capable of forming *bona fide* prions exhibit two-state discontinuous behavior under the specific experimental conditions used to generate these DAmFRET data. Shorter induction times or the use of a more sensitive cytometer may allow detection of missing low-FRET states for some proteins, while longer induction times or allowing the protein states to evolve through multiple rounds of cell division may allow detection of missing or infrequent high-FRET states for other proteins.

### Can DAmFRET data be used to quantify nucleation probabilities for proteins that undergo two-state discontinuous transitions?

As shown for the dataset of Alberti et al., two-state discontinuous transitions are likely to be nucleated phase transitions. The simplest mechanism for such transitions is that of homogeneous nucleation described in terms of classical nucleation theory [35, 38, 53, 54]. We define the probability of homogeneous nucleation as the probability of observing non-zero or high AmFRET states as a function of time and concentration. The probability of nucleation and assembly are correlated, and this is governed by the degree of supersaturation and the free energy barrier. The supersaturation ratio is quantified as 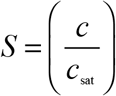 where *c* is the bulk concentration and *c*_sat_ is the saturation concentration above which the phase transition is thermodynamically favored. In DAmFRET, measurements are typically performed at a fixed time *t* across a population of cells containing a fixed concentration *c* of the protein of interest. Since each cell is akin to a femtoliter-scale test tube, each cell acts as a separate, independent experiment.

The fraction of cells in which assembly has occurred at time *t* and concentration *c* is used to define the probability 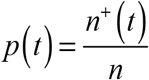. Here, *n*^+^ is the number of cells in which assembly has occurred at time *t*, and *n* is the total number of cells in which measurements are being made. Each cell is an independent observation volume in which two outcomes are possible namely, high AmFRET, implying the observation of assembly or no AmFRET, which means the lack of assembly in the cell. Accordingly, observations within each cell are Bernoulli trials, which are akin to a random experiment with exactly two outcomes. Accordingly, the overall outcomes across a population of cells can be treated as a binomial distribution. Since the outcome is also dependent on time and we are interested in the number of high AmFRET outcomes in a specific interval of time, the binomial distribution becomes a Poisson distribution, which is essentially a binomial distribution in the limit of infinite sub-divisions of the time interval [55].

We adapt the approach of Jiang and ter Horst [56], which uses the Poisson distribution to extract information regarding nucleation rates and barriers from distributions of induction time measurements. If the average number of nuclei *N* that form in a time interval is known, the probability of finding *k* nuclei within that time interval is given by the Poisson distribution as: 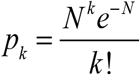. Note that the average number of nuclei that form in time interval *t* in volume *V* is directly related to the nucleation rate *J* because *N* = *JVt*. From the Poisson distribution, the probability that nuclei *do not form* in the time interval is given by *p*_0_ = *e*^−*N*^. Therefore, (1 – *p*_0_) = 1 – *e*^−*N*^ is the probability that at least one nucleus has formed in the interval of interest. Accordingly, the probability of observing a high AmFRET state at time *t* is *p*(*t*) = 1 – exp(–*JVt*) and this can be equated to the fraction of cells in which assembly has occurred at a given time and concentration. At a fixed time-interval, the probability of observing a fully assembled state is governed by the supersaturation *S* because 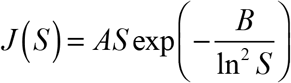. Here, *A* is a kinetic parameter that is governed by the rate of crossing the free energy barrier for nucleation, while *B* is a thermodynamic parameter governed by the size of the free energy barrier.

The Poisson distribution applies to a scenario in which supersaturation is achieved instantaneously and then fixed. While protein translation can be inhibited in cells by the addition of cycloheximide, this does not address the issue at hand, i.e., up to the point of cycloheximide addition, the concentration of protein, and therefore supersaturation, is continually changing. Although this may seem like a limitation, the reality is that, embedded within a single DAmFRET experiment is rich information about the concentration *and time* dependence of nucleation for tens to hundreds of thousands of cells, and this can be leveraged to provide an extraordinary advantage. By quantifying *p*(*t*) as a function of both time and concentration one can obtain a complete assembly probability landscape. This is shown in **Figure 8** for a specific choice of values for *A* and *B*. Here, we set *A* = 10^14^ m^-3^ s^-1^, *B* = 3.6, *c*_sat_ = 1.0 a.u., and *V* = 10^-18^ m^3^. We use a time range of 0 to 20 h, and a concentration range of 1.0 – 5.0 a.u. Note that we use acceptor intensity as a proxy for concentration and hence the choice of arbitrary units (a.u.) for concentrations. This is convenient since concentration shows up in terms of the supersaturation, which is dimensionless, and therefore the specific concentration scale is not relevant here.

**Figure 8.**
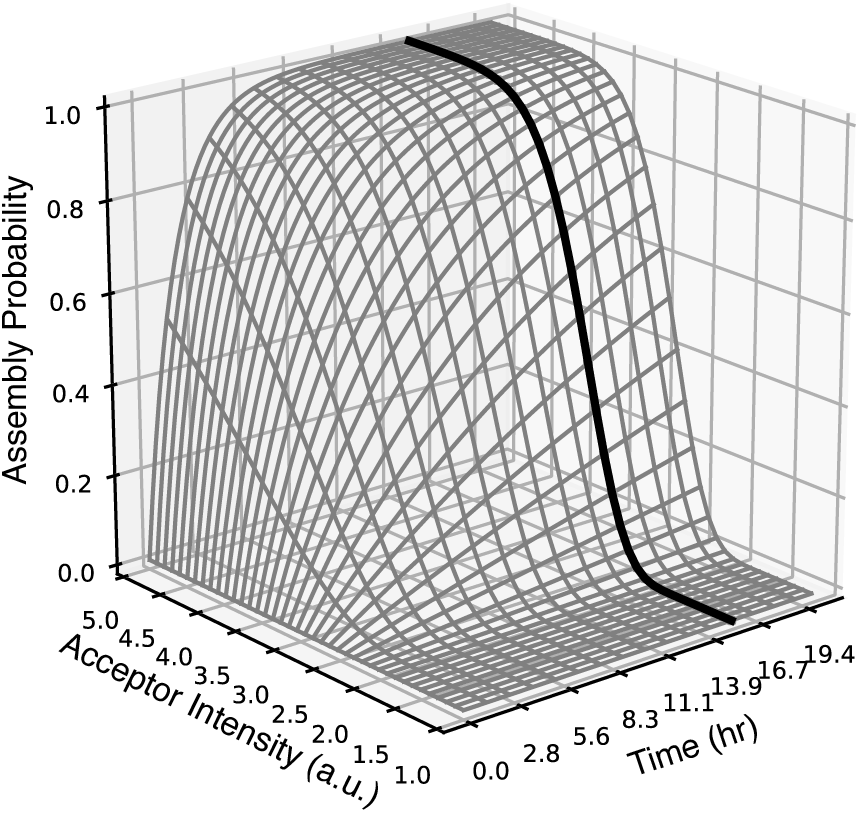
Modeling assembly probability as a function of time and concentration. Assembly probability versus time and assembly probability versus concentration are orthogonal planes forming a surface when plotted in three dimensions (time, concentration, assembly probability). As an approximation, DAmFRET data are in the assembly probability vs. concentration plane, at a single fixed time (black circles and line).

To map the assembly probability across time and concentration for a population of cells and fully capitalize on the time- and concentration-dependent information contained in DAmFRET data, we require knowledge of acceptor intensity as a function of time or at least upper and lower bounds on rates of protein expression. It is possible to observe individual cells via microscopy, while ensuring that they are incubated under conditions that are identical to the DAmFRET assay. Protein expression data for eight cells expressing the mammalian prion ASC (PYCARD) [57] fused to mEos3.1 were collected in this manner **(Figure 9a)**. Variability in expression levels across the cell population is apparent and this is the result of variation in plasmid copy number that was designed into the system. Although this approach reduces the throughput of the DAmFRET assay and does not yield the numbers desirable for large-scale statistical analyses, it shows that expression levels at distinct time points can be quantified.

**Figure 9.**
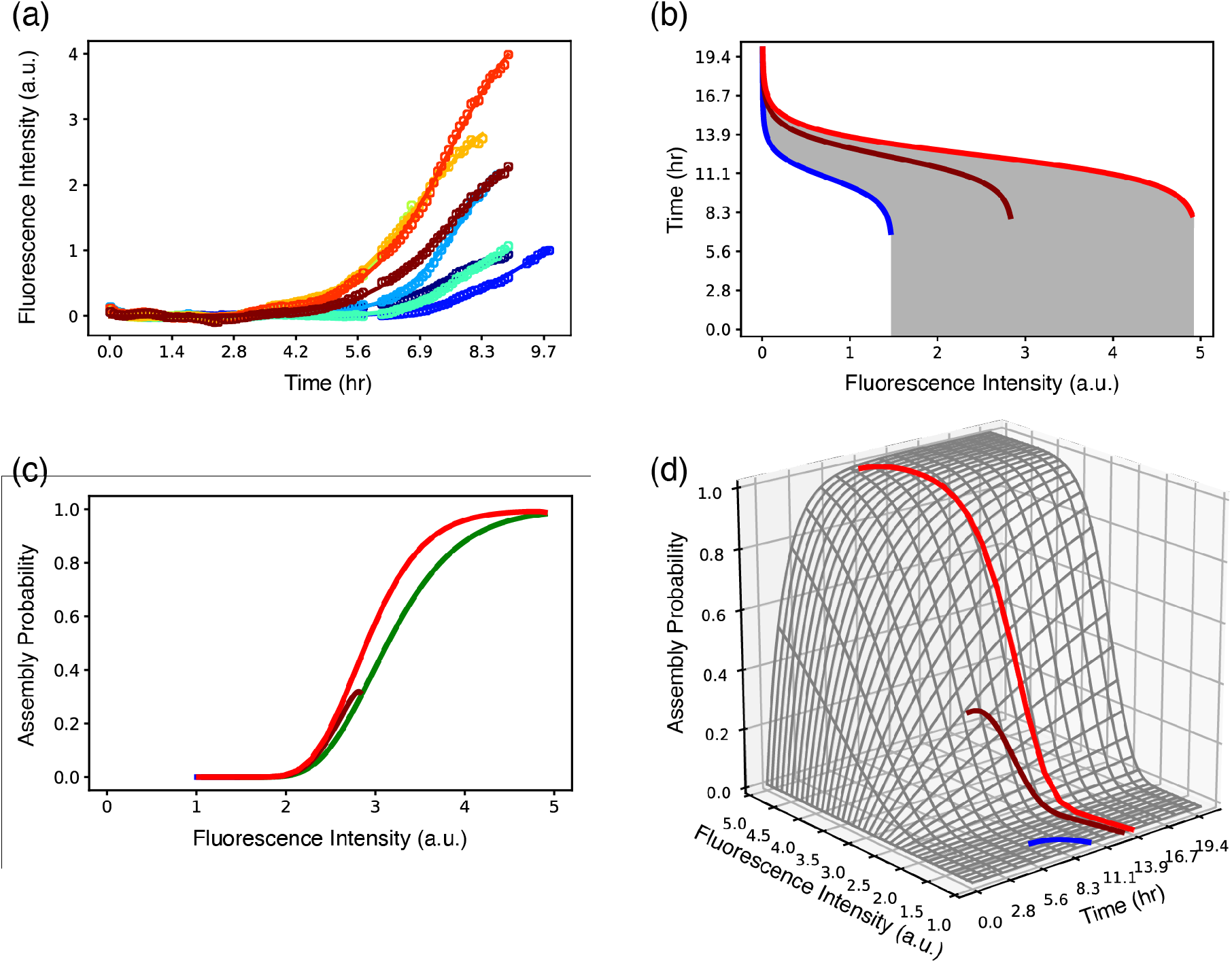
Modeling assembly probability using knowledge of protein expression trajectories. **(a)** Monitoring of protein expression in DAmFRET cells shows variability in accumulation of protein over time due to differences in plasmid copy number, as designed. Fluorescence intensity is used as a proxy for protein concentration. The data (open circles) are fit using a logistic function (lines). The raw data were from video S1 in the work of Khan et al. [39]. **(b)** To relate the expression trajectory to the three-dimensional plot more accurately, the data are converted to time elapsed at each concentration (i.e., time elapsed from when concentration C_i_ is reached (t_Ci_) to the time of measurement (t_m_), see text). Characteristic fast (red), intermediate (maroon) and slow (blue) trajectories were selected from the expression data for analysis in three dimensions. With the fit parameters from the red and blue traces as upper and lower limits, all combinations of parameters were used to calculate the range of possible logistic expression trajectories within these limits (gray fill). **(c)** Two-dimensional projection of the three-dimensional model with characteristic fast (red), intermediate (maroon) and slow (blue) trajectories selected from the expression data. The blue line is not visible because low expression did not result in appreciable assembly probability. The endpoints of the trajectories, defined as 99% of max concentration, provides a lower bound for assembly probability (green line). The upper bound for assembly probability at any concentration corresponds to the fastest expression rate (red line). **(d)** Protein expression versus elapsed time trajectories are plotted in three dimensions illustrating how the change in concentration over time during an experiment maps to the assembly probability landscape.

In typical chemical kinetics assays used to study phase transitions *in vitro*, purified and fully disaggregated proteins are dissolved or diluted into buffer prior to performing measurements as a function of time [58–60]. This ensures that measurements begin from a fully monomerized and dispersed phase. In contrast, the DAmFRET assay is performed in live cells that have been induced to express protein over a period of 10-24 hours. Unlike *in vitro* assays, the starting protein concentration is a moving target, since the protein of interest has gradually accumulated over a period of 10-24 hours prior to measurement. Further, the rate of protein accumulation varies from cell to cell. Because concentration is a moving target, the starting time *t*_0_ for the initiation of the phase transition is less well defined. We propose that the most accurate way to quantify incubation time at a particular concentration in the DAmFRET assay is to recognize that each concentration level *ci* attained during protein expression must be marked by a separate *t*_0_, which we designate as *t*_0*i*_, and hence there exists a distinct starting time *t*_0*i*_ for every *ci* reached. Therefore, time is counted as the time elapsed from time *t*_0*i*_, when a given concentration *ci* is reached, to the time of measurement designated as *t_m_* **(Figure 9b)**. This method of tracking time elapsed at each concentration only holds while the concentration is increasing. Therefore, we identify the time point when 99% of the maximum concentration is reached. The expression data are fit to a logistic function in order to facilitate extrapolation and interpolation of the expression trajectory in subsequent analysis. Based on our data and other published values of protein expression in yeast this seems to be a reasonable model for approximating expression under the *GAL1* promoter in yeast [61].

Using parameters from fits to the fastest and the slowest protein expression trajectories as limits, we used nearly exhaustive combinations of parameters between these limits to calculate possible expression trajectories that fall between the fastest and slowest trajectories **(Figure 9b, gray area)**. Where the lines approach verticality, the data can be ignored since the modified method of counting time no longer applies. In order to explore how measured expression trajectories within individual cells affect the DAmFRET readouts, we select representative fast, intermediate, and slow trajectories from the expression data and plot them as a function of both time and concentration **(Figure 9d)** using the assembly probability landscape shown in **Figure 8**. This analysis identifies the relationship between the DAmFRET data and the three-dimensional model, by demarcating the bounded range of assembly probabilities that can be fit to DAmFRET data.

Using the modeled range of expression trajectories, we estimated lower bounds for the assembly probability as a function of concentration in a two-dimensional projection of the three-dimensional data. The red, maroon and blue traces in **Figure 9c** indicate lower limits of assembly probability for the fastest, intermediate and slowest expression trajectories, respectively. While these plots show the full history of various cells in assembly probability versus concentration space based on protein expression trajectory within a cell, the actual measurement of FRET only happens when the cell is at the end of the plotted trace. As can be seen for the intermediate expression rate trace, the final concentration at the time of measurement can yield a lower probability of assembly than the lower concentration values that were passed through earlier in the trace. This is because the final concentration was only present briefly before the measurement was made, and lower concentrations that were reached earlier in that cell had a longer incubation time.

From our observations, we draw the following conclusions: 1) the endpoint represents a lower bound for assembly probability because it represents the shortest time spent at that concentration (**Figure 9c, green trace**); 2) the fastest expression trajectory provides an upper limit on the lower bound of the assembly probability because it represents the case in which the highest concentrations were achieved most rapidly, and 3) the maximum lower bounds on the assembly probabilities for all other expression trajectories will fall in between the values inferred for the slowest and fastest expression trajectories. We note that even though the measured assembly probability is attributed to the final concentration within a cell, that same cell also existed at lower concentrations for longer periods of time and therefore has increased probability of assembly as indicated by the maximum of the trace. This is evident in the red and green traces in **Figure 9c**, which demarcate the bounded range of assembly probabilities that can be fit to DAmFRET data.

Overall, we can conclude that measurements of a small number of protein expression trajectories should make it feasible to extract the nucleation probability for various proteins that undergo two-state discontinuous transitions. Essentially, the protocol to follow would be to categorize the nature of the transitions using the supervised approach we have introduced here. This, supplemented by measurements of a modest number ~10 expression trajectories for systems classified as undergoing two-state transitions, can be used to extract bounds on the in-cell saturation concentration and classical nucleation theory parameters *A* and *B*, which are directly related to the kinetics and thermodynamics of nucleation, respectively. The combination of these parameters provides a unique quantitative description of nucleation mechanism and paves the way for dissecting sequence-to-mechanism relationships. This is noteworthy because it represents acquisition of key biophysical quantities from measurements made directly in a cellular environment, rather than extrapolating from measurements made in the simplified context of a test tube.

### Do inferred values of c_50_ and the slope m provide useful information regarding the mechanism of nucleation?

As discussed above, classical nucleation theory allows for the prospect of fixing the observation time, varying the supersaturation and quantifying the fraction of proteins *g*(*S*) that have been incorporated into an assembled phase as a function of *S*. Previously, Khan et al., built on the conjecture of Sear [62] who proposed that *g*(*S*) can be described empirically using a Weibull distribution. These distributions were defined by shape and scale parameters designated as δ and *EC*_50_, respectively, which relate to the slope *m* and midpoint *c*_50_ that we calculate here by fitting logistical models to DAmFRET histograms [39]. The shape parameter δ was found to correlate with the amount of structural order in the Sup35 prion domain. With our usage of classical nucleation theory, we can directly test if and how the slope and midpoint parameters extracted from DAmFRET experiments are related to the mechanistically relevant parameters *c*sat, *A*, and *B*. Specifically, we wish to know if useful inferences can be forthcoming regarding the nucleation parameters in the absence of any additional information. We use classical nucleation theory, specifically the model based on the underlying Poisson distribution, to explore how the values of *m* and *c*_50_ of nucleation probability curves relate to the values of *A*, *B*, and *c*sat that were used to generate the data. It is necessary to use an expression trajectory, and for the sake of simplicity, we choose the highest rate of expression that we modeled previously. However, once the nucleation probability data are generated, we proceed with the subsequent analysis as if we have no knowledge of expression rates.

By assuming a fixed value for two of the three variables *A*, *B*, *c*_sat_, and varying the third within a reasonable range, we can explore how these variables affect *m* and *c*_50_. The parameter *A* was varied from 10^13^ to 10^15^ (spanning this range in log-scale increments) with *B* fixed at 3.0 and *c*sat fixed at 1.0. The resulting nucleation probability curves were collapsed onto distinct planes, and we assume these curves to be representative of the points acquired from fits to slices of DAmFRET data as previously described. We fit a logistic function to these points in order to obtain a slope, *m*, and midpoint *c*_50_ (**Figure 10a**). We then plotted *m* and *c*_50_ as a function of the values used for *A* (**Figure 10b-c**). We find that *m* has a strong dependence on *A*, as shown by the dramatic changes in slope of the curves in **Figure 10a**. Although the values of *c*_50_ also change with *A*, this effect appears to be determined by the change in slope rather than a shifting of the entire curve. This can be seen clearly in **Figure 10a**, where all of the curves begin to depart from zero nucleation probability at the same location (near an acceptor intensity of 2.0 a.u.) but with different slopes. Thus, *A*, the parameter that measures the effective shape of the nucleation barrier, appears to affect *m* directly and *c*_50_ indirectly.

**Figure 10.**
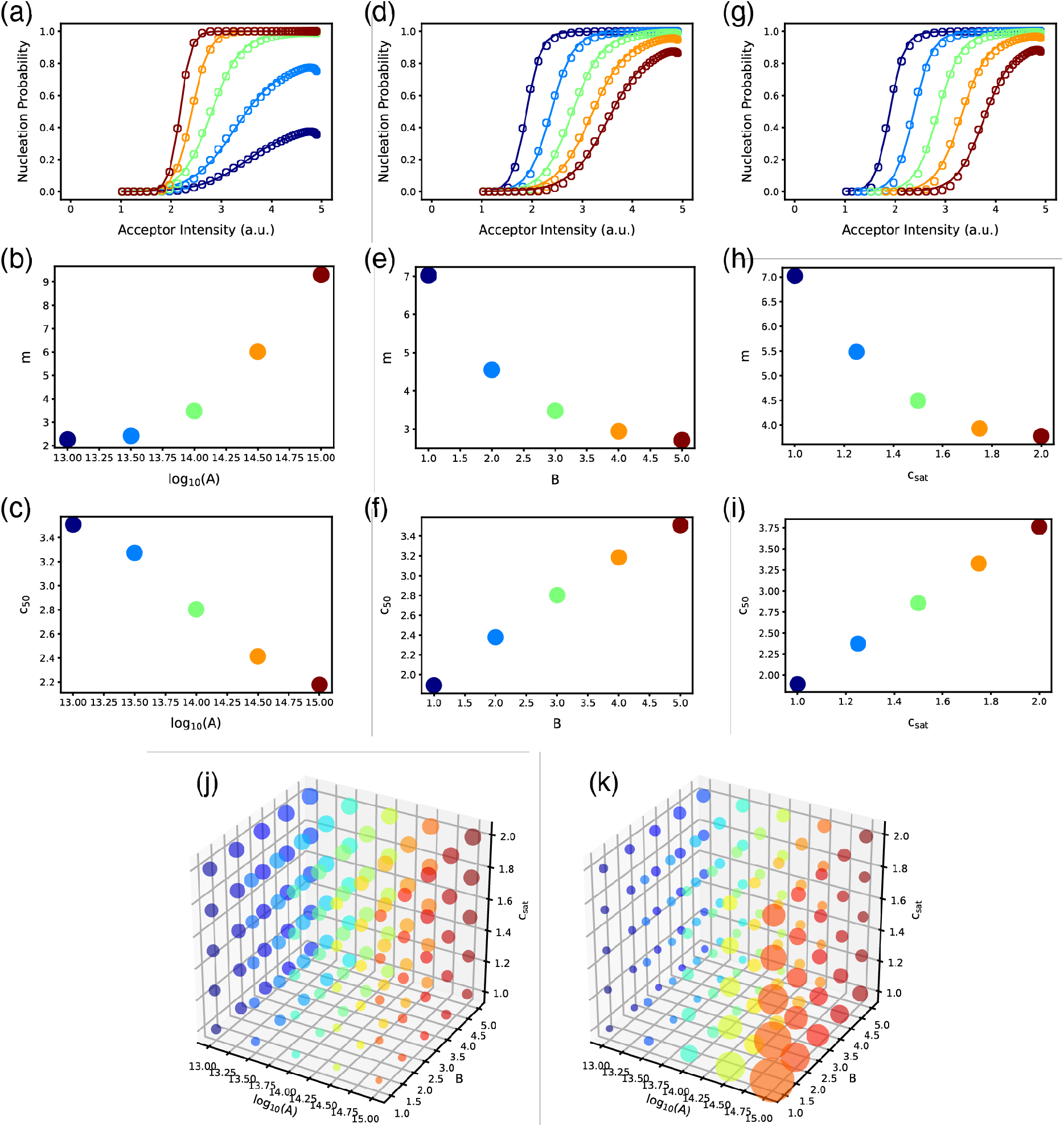
Relationship between the parameters *m* and *c*_50_, and the parameters from classical nucleation theory. Overall, we find that *c*_50_ is linearly correlated with *c*_sat_, whereas *m* is a complex convolution of contributions from *A*, *B*, and *c*_sat_. Assuming the highest expression rate (see text), and by varying one of the three parameters *A*, *B*, and *c*_sat_, while keeping the other two fixed, we generated curves for nucleation probabilities using the Poisson distribution model. The resulting data in three dimensions were projected onto distinct planes and plotted (panels a, d, and g, open circles) in order to simulate points acquired from fits to slices of DAmFRET data. A logistic function (solid lines) was fit to these points in order to obtain a slope, *m*, and midpoint *c*_50_. (a) The parameter *A* was varied from 10^13^ (dark blue) to 10^15^ (dark red) with *B* fixed at 3.0 and *c*_sat_ fixed at 1.0. (b) The relationship between slope, *m* and *A*. Colors correspond to those used in panel (a). Panel (c) The midpoints, *c*_50_, of the fitted curves in panel (a) are plotted against *A*. Panels (d), (e) and (f) are equivalent to panels (a), (b) and (c), except that the parameter *B* was varied between 1.0 and 5.0, while *A* was fixed at 10^14^ and *c*_sat_ was fixed at 1.0. Panels (g), (h), and (i) correspond to panels (a), (b), and (c), except that the parameter *c*_sat_ was varied between 1.0 and 2.0, while the parameter *A* was fixed at 10^14^ and the parameter *B* was fixed at 1.0. Panel (j) shows the fitted values of *c*_50_ represented by marker size for all combinations of parameters *A*, *B* and *c*_sat_. The positive linear correlation between *c*_50_ and *c*_sat_ holds for all combinations of *A*, *B* and *c*_sat_. In panel (k) the fitted values of *m* are represented by marker size as a function of all combinations of parameters *A*, *B* and *c*_sat_. Many combinations of these parameters result in similar intermediate slopes, while the largest slopes arise for high values of *A* and low values for *B* and *c*_sat_.

Next, we repeated the above analysis by varying *B* between 1.0 and 5.0, fixing *A* at 10^14^ and *c*sat at 1.0 (**Figure 10d**). Unlike *A*, we find that *B*, which quantifies the barrier height, has an inverse relationship with *m* (**Figure 10e**). Importantly, the value of *B* directly affects both *m* and *c*_50_ (**Figure 10f**). This is evident in the fact that in addition to changing slope, the onset of the transition i.e., the initial departure from zero nucleation probability, shifts further to the right with increasing *B* (**Figure 10d**).

Repeating the analysis with the *c*_sat_ varied between 1.0 and 2.0, fixing *A* at 10^14^ and *B* at 1.0 revealed that *c*_sat_ and *c*_50_ are positively correlated with one another (**Figure 10g, i**). This is true for all combinations of *A*, *B* and *c*_sat_ in the ranges that we tested (**Figure 10j**). Taken together, we observe that while there are many combinations of *A*, *B* and *c*_sat_ that result in similar intermediate slopes, the largest slopes corresponding with the steepest transitions are the result of high *A*, low *B*, and low *c*_sat_ (**Figure 10k**). Accordingly, the steepest slopes correspond with the highest rates, lowest barriers, and lowest saturation concentrations. These analyses indicate that even in the absence of information about expression rates, *c*_50_ can provide useful information for comparing different proteins. Even though *c*_50_ is influenced by *A*, *B*, and *c*_sat_, it has a strong positive correlation with *c*_sat_ across all values of *A* and *B* and this is a direct consequence of the two-state behavior suggesting it can be used as a reasonable proxy for *c*_sat_, which measures the driving forces for phase separation. On the other hand, *m*, (or the δ parameter from the original work [39]) is a convolution of contributions from *A*, *B* and *c*_sat_, and therefore *m* on its own is not a readily interpretable parameter. However, it still carries information regarding the relative drive for nucleation, whereby steeper slopes are often due to a combination of high values for *A*, and low values for *B* as well as *c*_sat_.

### Analysis of DAmFRET data yields information regarding the driving forces for assembly

DAmFRET data are information-rich, and additional insights can be extracted from the histograms. For instance, we can extract accurate quantitative estimates for *c*_50_ values for all proteins that show two-state behavior. **Figure 11** shows the *c*_50_ for all 23 cPrDs which were classified as undergoing a two-state transition. By extracting *c*_50_ values we can rank order these proteins according to their *c*_50_ values. Given the relationship shown in **Figure 10**, for proteins that show two-state discontinuous behavior, the lower the *c*_50_, the lower the concentration needed for assembly, and thus the greater the driving force for assembly. The analysis in **Figure 11** shows a two-order of magnitude variation in inferred *c*_50_ values. It is worth emphasizing that all cells for all measurements were prepared in identical fashion. All proteins are probed across overlapping concentration ranges in similar cellular environments. Therefore, to zeroth order, the only variable distinguishing different experiments is the sequence of the protein whose phase behavior is being probed. Accordingly, to zeroth order, assuming the cellular factors do not have cryptic sequence-specific responses as modulators of phase behavior, the data for *c*_50_ help quantify the impact of sequence-encoded interactions as drivers of phase transitions. If there are sequence-specific effects of cellular factors, it still follows that the modulation of the driving forces is governed by the sequences of the proteins whose phase behavior is being probed.

**Figure 11:**
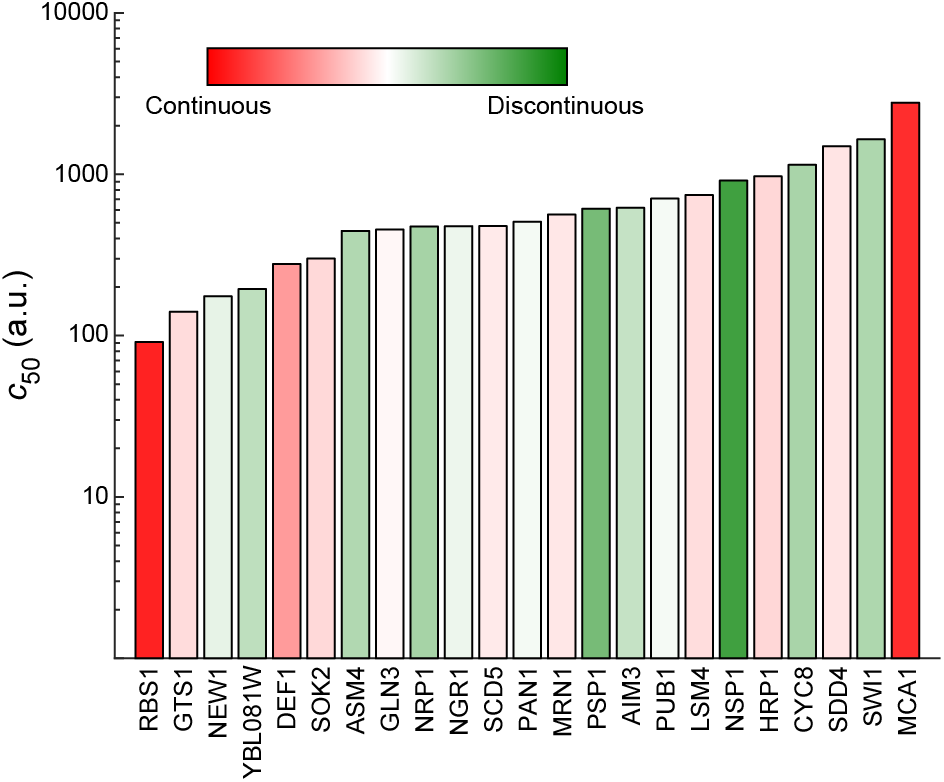
Extracted *c*_50_ values for all 23 cPrDs which were classified as undergoing a two-state transition from the DAmFRET data. The color and shade of each bar corresponds to the degree of discontinuity in the transition.

### Extracting sequence-to-assembly relationships from DAmFRET data

The minimum *R*^2^ value in the transition expression region yields information on the degree of discontinuity in the two-state transition. Larger *R*^2^ values imply that the DAmFRET data fits well to a sum of two Gaussians in the transition region and thus most cells are either at zero AmFRET or at high AmFRET, thus implying the transition is discontinuous. In contrast, small *R*^2^ values imply that AmFRET values for most cells falls between zero and high values for AmFRET. The fraction of cells that fall between the two limits should increase as the slope of the transition becomes smaller, and thus the *R*^2^ should be lowest for two-state continuous cases with shallow transitions. We can use this relationship to examine whether there are certain amino acids that correlate with one type of two-state transition over another. **Figure 12** plots the fraction of a given amino acid against the minimum *R*^2^ value in the transition region. Each point represents one in-cell experimental replicate per cPrD. These data are shown for the four amino acids with the highest positive versus negative linear correlations. The correlations for all amino acids are showed in **Figure S6**. We find that an increase in the fraction of Phe or Thr correlates positively with increased discontinuity. In contrast, an increase in the fraction of Pro or Tyr correlates negatively with increased discontinuity. The impact of Pro on the discontinuity of the two-state transition is not surprising given its tendency toward disrupting secondary structures other than beta turns. However, the non-equivalence of Phe and Tyr as promoters of two-state discontinuous transitions is surprising. This suggests that titrations of Phe versus Tyr contents in low complexity domains might be a way to tune the discontinuity of a phase transition and the tendency for forming liquid-like condensates versus ordered assemblies [63, 64]. A recent study has uncovered clear differences between Phe and Tyr as drivers of condensate formation via phase separation aided percolation transitions in prion-like low complexity domains (PLCDs). Further, analysis across homologous sequences highlights a negative correlation between Phe and Tyr contents [65]. Taken together with findings in **Figure 11**, a prediction that emerges is that weakening of the driving forces for condensate formation by lowering the Tyr content and increasing the Phe content also enables a facile transition to ordered assemblies. This would suggest that there is likely to be discernible code for distinguishing sequences that drive condensate formation that also turnover into ordered assemblies versus those that form stable, reversible condensates that are unlikely to undergo disorder-to-order transitions.

**Figure 12:**
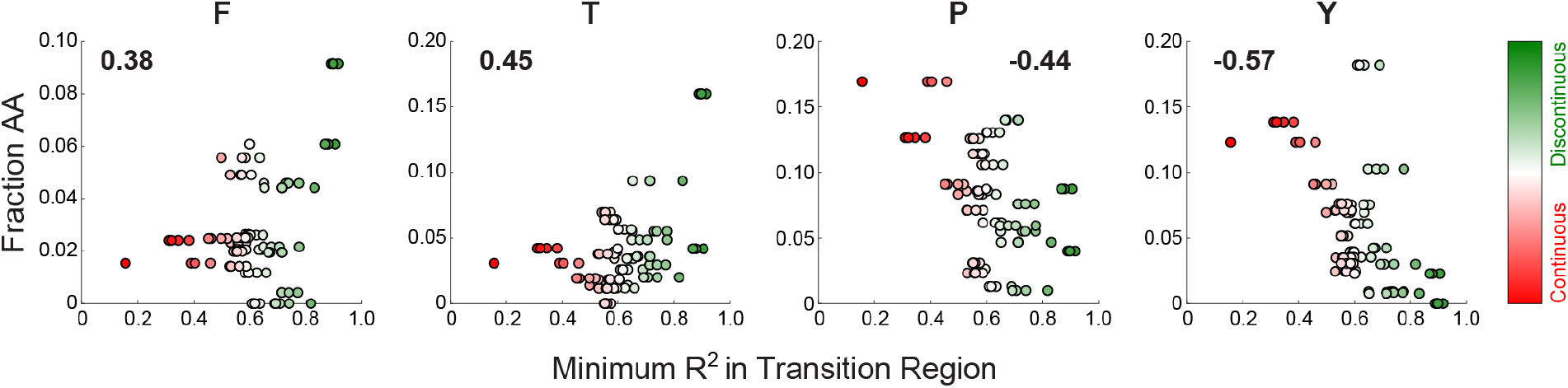
Correlation between amino acid frequency and degree of discontinuity for all 23 cPrDs classified as undergoing a two-state transition. Each point corresponds to an experimental replicate. The color of each point denotes the degree of discontinuity of the transition. Numbers indicate the Pearson *r-*values used to quantify positive or negative correlations.

## Discussion

There is growing interest in measuring phase transitions in live cells [34, 39, 40, 42]. Of particular interest are results of measurements made under conditions where the effects of active processes are minimized. These measurements are helpful for understanding how the milieu of a living cell impacts the intrinsic driving forces for phase transitions [39]. These experiments, performed as a function of controlling the expression levels of the protein of interest, can help in mapping the sequence-specific free energy landscape that underlies the driving forces for and mechanisms of phase transitions that are under thermodynamic control.

Here, we analyze in-cell phase transitions by allowing for a range of transition categories and use a supervised approach to develop a method that enables the automated analysis of DAmFRET data. This approach affords classification of the type of phase transition and comparative assessments of the driving forces for phase transitions. We applied our method derived from supervised learning to analyze DAmFRET data for 84 different candidate prion domains. Our analysis helps categorize the phase transitions for each of these domains and identify sequences that clearly show two-state behavior. Among the trends that emerge, we find a noticeable negative correlation between the Tyr / Pro content and systems that undergo discontinuous two-state transitions. Conversely, we observe a weak positive correlation between the Phe / Thr content and the propensity for showing discontinuous two-state behavior. We envisage the possibility of using information gleaned across large libraries of sequences to design novel domains that undergo specific categories of phase transitions. The ability to quantify the sequence contributions to *c*_50_ values also affords the prospect of manipulating the driving forces for forming prion-like assemblies through sequence design.

In addition to categorizing sequence-specific phase transitions and quantifying the driving forces for these transitions, we show how classical nucleation theory can be brought to bear for estimating the lower bounds on nucleation probabilities of systems that undergo discontinuous two-state transitions. This analysis requires independent measurements of expression trajectories, although this information is not available across the spectrum of proteins that have been interrogated using DAmFRET. If the parameters *S*, and *J* can be extracted using analysis of the DAmFRET data as a function of *S*, then the parameters *A* and *B* can be determined by plotting *J*/*S* versus ln^-2^*S*. This would allow mechanistic inferences such as estimates of free energy barriers to be extracted from a single DAmFRET experiment. Although expression trajectories are not currently obtained with the same level of throughput and speed as the generation of DAmFRET histograms, the data regarding expression trajectories are essential for estimating nucleation probabilities. It suffices to have these data for the fastest and slowest expression trajectories. High-throughput methods for obtaining upper and lower bounds on expression levels versus time should be feasible, and promising options are being explored. Supplementing datasets by defining bounds on probable expression rates will go a long way toward facilitating a near complete mechanistic understanding of nucleated phase transitions for systems that undergo two-state discontinuous transitions. The physical parameters extracted from the application of classical nucleation theory to the analysis of DAmFRET histograms augmented by expression trajectories could be useful in enabling proteome-wide comparisons of the driving forces for forming ordered assemblies.

Finally, the packaged code for supervised learning and for automated analysis of DAmFRET data are available via Github (https://github.com/pappulab/damfret_classifier). This package is distributed as open source, available for free download and usage, and users are invited to contribute code and insights to further the development of the package that is intended to enable automated classification of phase transitions and mechanistic inferencing based on DAmFRET data.

## Materials and Methods

### Biological reagents and Yeast transformation

The yeast strain used was rhy1713 as described in previous work [39]. The strain is a knockout of CLN3 combined with a galactose-inducible overexpression of WHI5, thereby breaking the G1 cell cycle checkpoint and inducing cell arrest [66]. This allowed us to detect only de novo nucleation events by preventing mother-daughter cell propagation of the prions. **Table S4**, attached as an Excel spreadsheet, lists all plasmids used in this study. Plasmid number, gene name, cell count and encoded polypeptide sequences for each gene region are listed for each construct and replicate. Cells were transformed using a standard lithium acid transformation protocol [67].

### Preparation of Cells for Cytometry

Protein expression is induced in a 2% synthetic galactose (SGal)medium for 14 hours before being resuspended in fresh SGal for 4 hours to minimize autofluorescence. After 18 hours of total induction, the cells are uniformly illuminated with 405nm violet light for 25 minutes to convert a highly reproducible ratio of the mEos3.1 from the green donor form to the red acceptor form.

### DAmFRET Cytometric Assay

Following photoconversion, acceptor fluorescence intensity and FRET are measured using a flow cytometer. The ratio of indirect and direct acceptor fluorescence (595 ± 10 nm when excited with 488 nm or 561 nm light, respectively) is referred to as AmFRET, and this is used to measure the extent of ordered assembly within each cell. In the original implementation of the DAmFRET assay, the acceptor intensity, excited directly, was converted into units of concentration by dividing it by the measured cytosolic volume of the cell. In this work, cell imaging during flow cytometry measurements was bypassed to increase throughput by greater than 150-fold. The acceptor fluorescence intensity is still measured and used to monitor expression level, and this serves as a useful proxy for protein concentration in the cell.

### Additional details for the generation of the synthetic dataset

The number of points generated, which mimics the number of cells being interrogated in a DAmFRET measurement, was randomly selected to be between 10^4^ and 1.5×10^4^. In order to represent different distributions of points across the concentration range, an additional 10^4^ to 1.5×10^4^ points were added to each dataset in three ways: points were chosen from a uniform distribution spanning the full range from 2.0 to 8.0, only at concentrations above the *c*_50_, or only at concentrations below *c*_50_. The values of AmFRET for these additional points were determined as described above. Data were also generated without these additional points to simulate datasets obtained using fewer measurements. These smaller datasets had between 10^4^ and1.5×10^4^ points in total, rather than 2×10^4^ or 3×10^4^ points.

### Method for classification of DAmFRET histograms

For each replicate, non-overlapping slices are made along log_10_(Expression). These slices are made in intervals of 0.5 (synthetic data), and 0.2 (real data). To reduce noise contributions at the extrema, our method uses a low cutoff of 3.0 and a high cutoff of 10.0 in log_10_(Expression) for synthetic data, and slices are only collected between these limits. A low cutoff of 1.5 and a high cutoff of 5.0 is employed for real data. For each slice, a normalized 1D histogram of the AmFRET counts is determined by binning the synthetic data in intervals of 0.1, while the real data is binned in intervals of 0.02. That histogram is fit to the sum of two Gaussians. The first Gaussian is centered at AmFRET=0, while the second Gaussian is centered at the position corresponding to 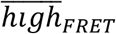. Here, 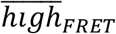 is calculated by taking the mean of AmFRET>0.5 (synthetic data) or AmFRET>0.05 (real data) in each expression level slice and taking the maximum of this value over all expression slices. The *R*^2^ value of the fitted function is saved to be utilized later on.

Using the fitted parameters, each Gaussian is numerically integrated to extract the area under the curve, yielding the quantities *g*_1_ and *g*_2_, for the first and second Gaussians, respectively. The fraction assembled in a given expression level, *i*, is then given by *f*_*A,i*_ = *g*_2_⁄(*g*_1_ + *g*_2_). To determine if no transition is observed in the DAmFRET histogram the change in the fraction assembled, Δ*f*_*A*_, is calculated as:

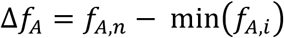

where *f*_*A,n*_ is the fraction assembled in the last expression level slice. We define an assembly threshold, Δ*f*_*A,thresh*_, for one-state assembly as 0.10 (synthetic data) and 0.15 (real data). If Δ*f*_*A*_ < Δ*f*_*A,thresh*_, then the DAmFRET histogram does not show a transition and can be classified as one-state. If the mean AmFRET<0.5 (synthetic data) or AmFRET<0.05 (real data), then the DAmFRET histogram is classified as one-state: no assembly at all expressions (blue). Else, the DAmFRET histogram is classified as one-state: assembled at all expressions (black). To determine the confidence in either assignment, we calculate a confidence score of the system using the deviation of Δ*f*_*A*_ from Δ*f*_*A,thresh*_:

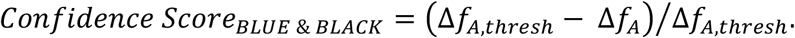

Since this score is normalized, it ranges from 0 to 1.

If Δ*f*_*A*_ ≥ Δ*f*_*A,thresh*_, the system is likely to be undergoing some kind of transition. Therefore, we fit a logistic function to fraction assembled profile using the equation: 1/(1+exp(-(*c*-*c*_50_)/*m*)). Here, *c* is the log_10_(Expression) of a given slice, *c*_50_ is the log_10_(concentration at which 50% of cells are in the assembled state), and *m* is the inverse of the slope. Thus, we can extract *c*_50_ from the fit of the fraction assembled to the logistic function.

To account for the fact that real data can limited and / or noisy at the extrema of the expression level range, we add a couple additional checks for one-state behavior when analyzing real data. If *c*_50_ ≤ 1.5, i.e., below the lower limit of log_10_(Expression) used in our analysis of the real data, then this implies the system transitioned at low log_10_(Expression) values at which we have limited and / or noisy information. Thus, we also classify these histograms as one-state: assembled at all expressions (black). For these cases, the confidence score is just set to one.

At this point we perform our final check on whether a system has undergone no assembly at all expression levels. If *c*_50_ ≤ 4.0, then we check the number of measurements that are above *c*_50_. If this number is less than or equal to 20, then we ascribe these points to noise and the corresponding DAmFRET histogram is classified as one-state: no assembly at all expressions (blue). If *C*_50_ > 4.0, then the transition is at the edge of the expression level range, which implies *c*_50_ is not likely well defined. Thus, we just examine the number of points with AmFRET>0.05 corresponding to an expression level above *c*_50_. This checks the number of points that are genuinely in the assembled state above *c*_50_. If this number is less than or equal to 20, then we also ascribe these points to noise and the corresponding DAmFRET histogram is classified as one-state: no assembly at all expressions (blue). In both cases, the confidence score of these assignments is set to one.

Next, we check whether there is enough data to classify the transition. This check was not part of the method to analyze the synthetic data, but this checkwas added for the real data to account for cases in which there was not enough data to classify the transition. If the fraction of cells above the *c*_50_ (*f*_*c*50_) is less than 10% of the total number of cells for that replicate, the system is classified as undergoing an infrequent transition (yellow). Unlike the one-state classes, our check for this class involves first eliminating the DAmFRET histogram as being classified as one-state. Thus, our confidence score in the assignment of an infrequent transition must reflect the multiple checks that are performed to lead to this assignment. We calculate *Score*1 = min (1, (Δ*f*_*A*_ − Δ*f*_*A,thresh*_)/ (0.5 − Δ*f*_*A,thresh*_)) and *Score*2 = (0.1 − *f*_c50_)/0.1. Here, *Score*1 checks the deviation from the one-state criterion and *Score*2 checks the deviation from the infrequent transition criterion. The final confidence score is set to be the minimum of *Score*1 and *Score*2.

The remaining DAmFRET histograms undergo some sort of transition. Thus, we utilize the *R*^2^ values of the fit to the sum of two Gaussians to determine features of the transition and further classify these histograms. Histograms that show a two-state continuous transitions should have low *R*^2^ values around *c*_50_ given that a majority of the points will be between AmFRET=0 and 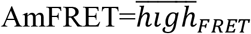. Thus, we identify an expression level window of *c*_50_ −/+1. If *c*_50_+1 exceeds the expression level range, then the last four expression slices are used for the window. Then, within this identified window both the maximum absolute change in consecutive *R*^2^ values, 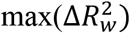, and the minimum *R*^2^ value, 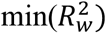, is recorded. Here, the subscript *w* denotes that we are only examining the *R*^2^ values that correspond with expression level slices around *c*_50_ as described above. If 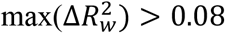 and 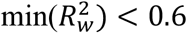, then the DAmFRET histogram is classified as undergoing a two-state continuous transition (red). The confidence score in this classification is then the minimum confidence of the preceding three checks. Specifically, we calculate *Score*1 = min (1, (Δ*f*_*A*_ −Δ*f*_*A,thresh*_)/(0.5 − Δ*f*_*A,thresh*_)) and *Score*2 = min(1, (*f*_*c*50_ − 0.1)/(0.3 − 0.1)) and 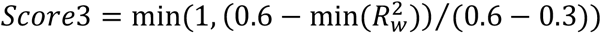. Here, *Score*1 checks the deviation from the one-state criterion, *Score*2 checks the deviation from the infrequent transition criterion, and *Score*3 checks the deviation from the two-state continuous transition criterion. The final confidence score is then the minimum of these three values.

Next, we sought to determine if any of the remaining DAmFRET histograms showed multi-state transition behavior. From the synthetic 2-step data, we noticed that the *R*^2^ values tended to linearly decrease with increasing expression level slice. This is due to the fact that the extracted 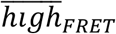 is a convolution of several high AmFRET states that are overlapping in their expression level range and thus neither high AmFRET state ever fits well to the sum of two Gaussians. To classify DAmFRET histograms that show this behavior, we fit the full *R*^2^ profile to a linear fit and restricted the maximum slope of that fit to be zero. Then the *R*^2^ value of this fit was extracted, 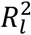. If 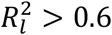, then the DAmFRET histogram was classified as undergoing a higher order state transition (magenta). As before, the confidence score in this classification was calculated using the previous checks. Here, we have *Score*1 = min (1, (Δ*f*_*A*_ − Δ*f*_*A,thresh*_)/(0.5 − Δ*f*_*A,thresh*_)) and *Score*2 = min(1, (*f*_*c*50_ − 0.1)/(0.3 − 0.1)) and 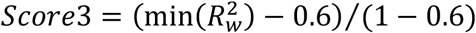 and 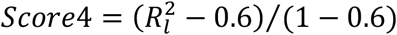. The final confidence score is the minimum of all four values.

Finally, the remaining DAmFRET histograms were classified as undergoing a two-state discontinuous transition (green) as the data did not show any features that led to the data failing the null hypothesis. The confidence score for this classification was set to the minimum of the following four scores: *Score*1 = min (1, (Δ*f*_*A*_ − ΔΔ*f*_*A,thresh*_)/(0.5 − ΔΔ*f*_*A,thresh*_)) and *Score*2 = min(1, (*f*_*c*50_ − 0.1)/(0.3 − 0.1)) and 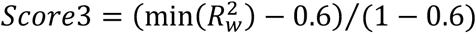 and 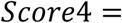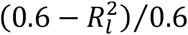

### Alberti et al. Dataset and Creation of DAmFRET histograms

DAmFRET histograms were collected for all 94 Alberti et al. cPrDs, 93 were performed in quadruplicate and one construct was performed with eight repeats. Each replicate is composed of 0 – 170,000 individual cell measurements of AmFRET at a given expression level. **Figure S7** shows the distribution of the number of cell measurements for all replicates. Given the wide range in the number of individual cell measurements performed across the dataset, we used an information theoretic approach to identify the ideal common grid size for our two-dimensional DAmFRET histograms of expression level and AmFRET. To determine an acceptable grid size which could be applied to all replicates for subsequent analysis, we examined the information density quantified by the Shannon Entropy, 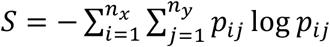, and its change as a function of increasing grid size, on a subset of replicates. Our analysis, shown in **Figure S8** and described in further detail in the supplementary material, led to the choice of a 300×300 common grid size. This grid size was chosen to maintain a large amount of information across varying numbers of individual cell measurements. However, given that replicates at lower measurement counts tend to experience significant |Δ*S*| loss at the 300 × 300 grid size, any subsequent analysis may be impaired. Hence, we excluded cPrDs who had at least one replicate with < 2 × 10^4^ individual cell measurements. This left us with 84 cPrDs for subsequent analysis.

### Determining the degree of discontinuity in the transition of a cPrD classified as undergoing a two-state transition

The *R*^2^ value around the expression slice that corresponds to log_10_(*c*_50_) yields information on how well a sum of two Gaussians can capture the 1-dimensional AmFRET histogram at the transition. Thus, we sort the cPrDs by their mean minimum *R*^2^ in the region that corresponds to the expression level slices within the window of log_10_(*c*_50_) −/+1, 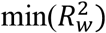, to order them by the degree of discontinuity in the transition. To color the data in plots **Figure 5E**, **Figure 11** and **Figure 12** we take advantage of the fact that one of the criteria for classification of DAmFRET data into the two-state continuous class is that 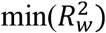 must be less than 0.6. Thus, any cPrD that has a mean 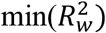 of less than 0.6 is colored red (two-state continuous) and any cPrD that has a mean 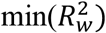 of greater than 0.6 is colored green (two-state discontinuous). Then, the alpha color for each of the two-state cPrDs / cPrD replicates is based on how far the 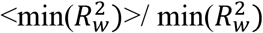 is from the threshold of 0.6. For those cPrDs classified as two-state discontinuous the alpha color is set to 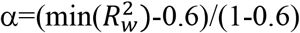. For those cPrDs classified as two-state discontinuous the alpha color is set to 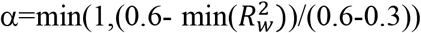. A lower limit of 0.3 rather than 0.0 is used in the latter case given that *R*^2^ values rarely ever drop below 0.3.

### Microscopy

Yeast cells expressing ASC-mEos3.1 were grown overnight in synthetic media containing 2% dextrose while shaken at 30°C. Cells were then loaded into a CellASIC ONIX microfluidic device (Millipore Sigma B04A03). Media containing 2% galactose was flowed through the microfluidic device at a rate of 5kPa from 2 wells at a time. Timelapse images were acquired on an Ultraview Vox (Perkin Elmer) Spinning Disc (Yokogawa CSU-X1). Images were collected with an alpha Plan Apochromat 100x objective (Zeiss, NA 1.4) onto an Orca R2 camera (Hamamatsu, C10600-10B). mEos3.1 was excited with a 488nm laser, and the fluorescence emission was collected through a 525-50nm bandpass filter. Images were collected as z-stacks with 0.5 um steps (41 slices) and a 30ms camera integration time every 5 minutes. Additionally, a single transmitted light image was acquired in the middle of the z stack with an integration time of 200ms. Each time point was sum projected and the resulting time course was registered to reduce movement of the cells. Regions of interest were then drawn in individual cells and the mean intensity of the sum projected fluorescence was measured from the beginning of the time course until the end.

## Acknowledgments

This work was supported by grants from the US National Science Foundation (MCB1614766 and DMR 1729783 to RVP) and US National Institutes of Health (5R01NS056114 and 1R01NS089932 to RVP, and R01GM130927 to RH). We thank Furqan Dar for helpful discussions and Matthew King and Min Kyung Shinn for comments on the manuscript.

