## Supporting Information for "Mechanistic inferences from analysis of measurements of protein phase transitions in live cells"

**Figure S1: The sum of two Gaussian fits to the 1D histogram of $\Omega_{FRET}$ for each expression level slice and the corresponding fraction assembled profiles extracted from these fits for the five synthetic DAmFRET histograms shown in Figure 2**. Displayed are: Panel **A** (One-State: No Assembly); Panel **B** (One-State: All Assembled); Panel **C** (Two-State: Discontinuous); Panel **D** (Two-State: Continuous); and Panel **E** (Three-State: Discontinuous). Each column shows the fits to the 1D histogram of $\Omega_{FRET}$ for a different expression level slice.

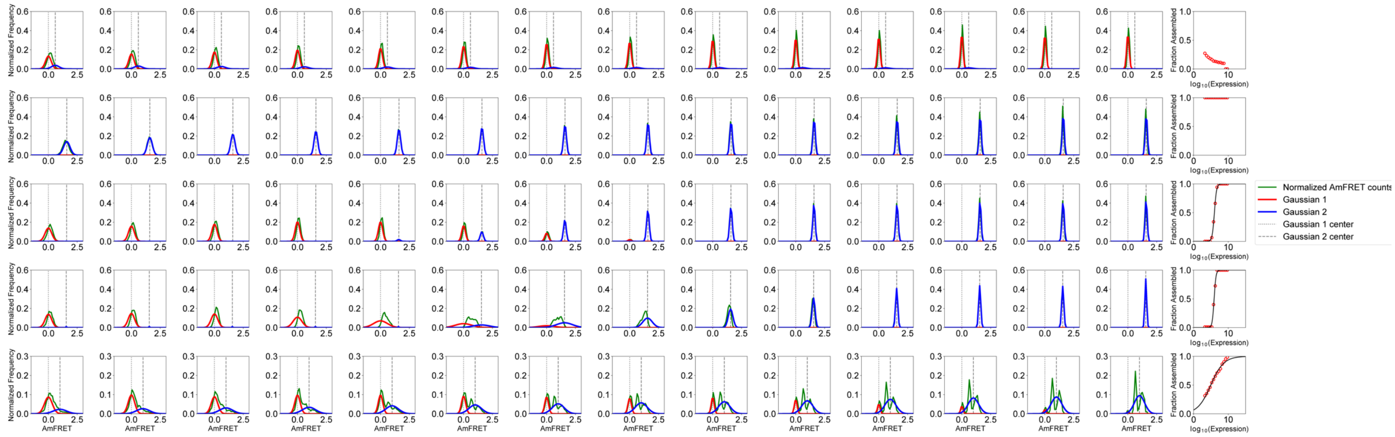

**Figure S2: Synthetic DAmFRET histograms sorted by their classification**. The five classes are: assembled at all expression levels (black class), no assembly at all expression levels (blue class), two-state discontinuous transition (green class), higher order state transition (magenta class), and two-state continuous transition (red class). The number in the top right-hand corner of each histogram indicates the confidence score of the classification. Each histogram is titled according to **Table S3** which describes the parameters used to generate the given synthetic histogram.

Assembled at all expression levels (black) [ 1 of 4 ]

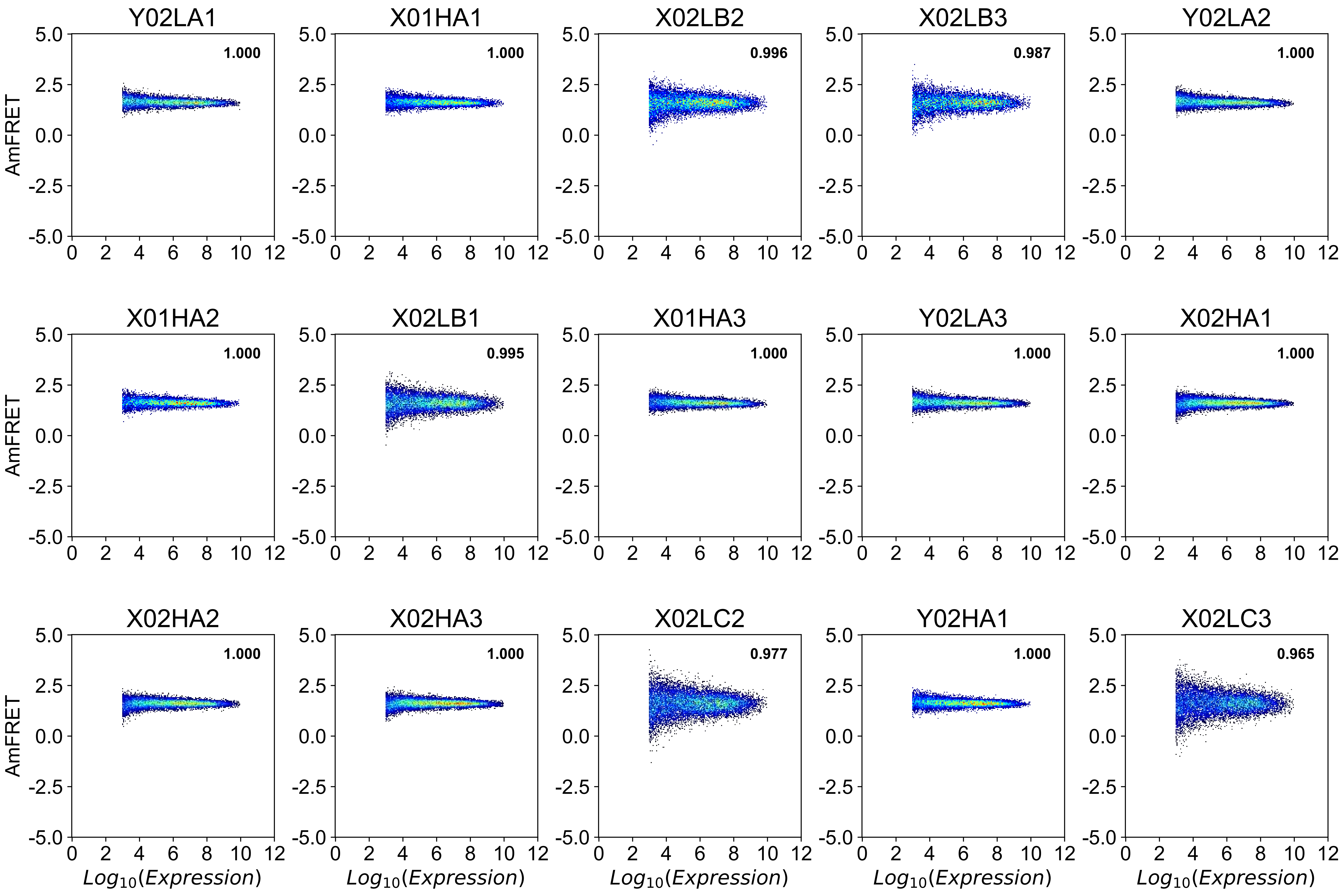

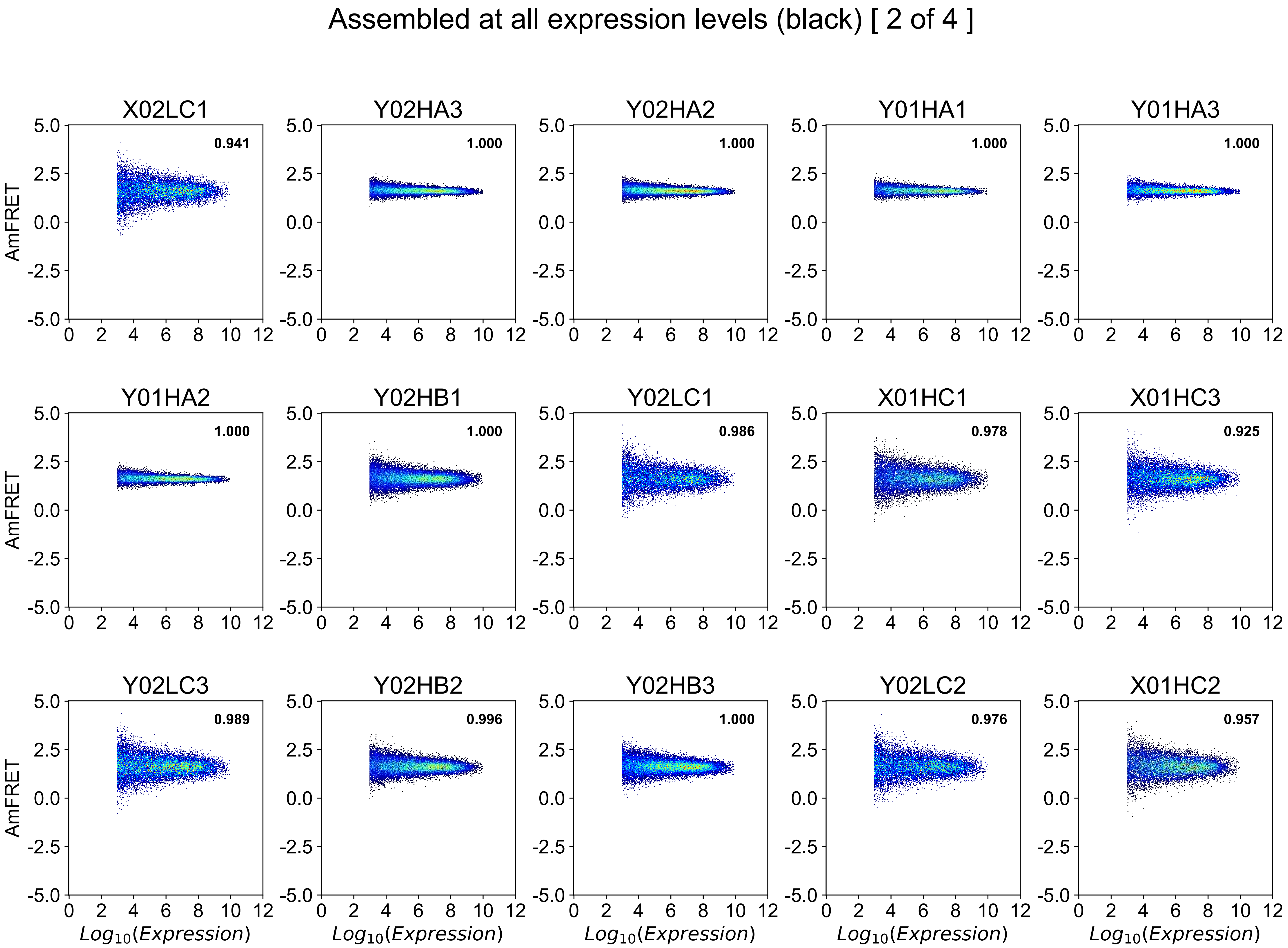
Assembled at all expression levels (black) [ 2 of 4 ]
X02LC1
0.941
Y02HA3
1.000
Y02HA2
1.000
Y01HA1
1.000
Y01HA3
1.000
Y01HA2
1.000
Y02HB1
1.000
Y02LC1
0.986
X01HC1
0.978
X01HC3
0.925
Y02LC3
0.989
Y02HB2
0.996
Y02HB3
1.000
Y02LC2
0.976
X01HC2
0.957
AmFRET
Log10(Expression)

### Assembled at all expression levels (black) [ 3 of 4 ]

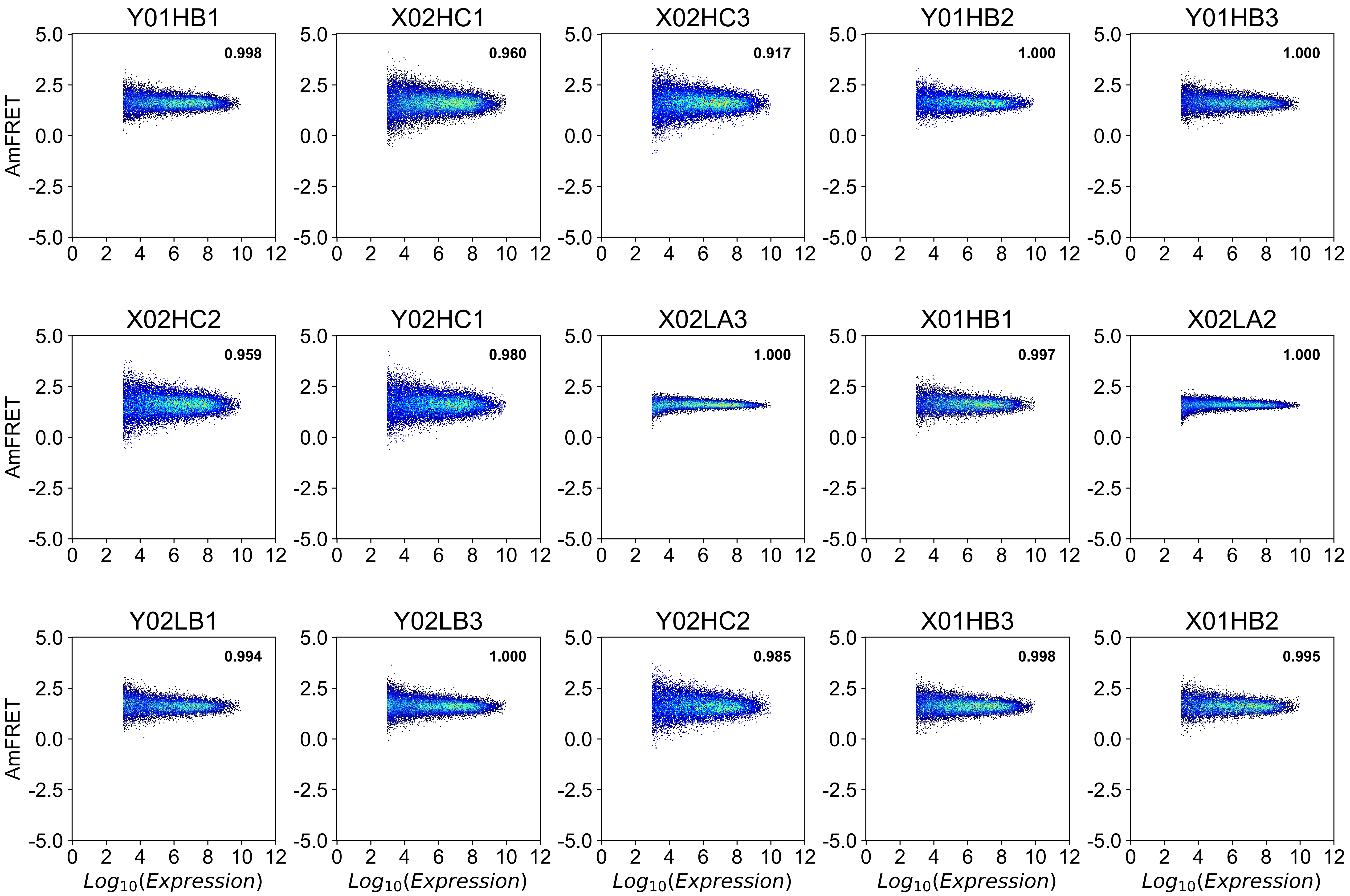

### Assembled at all expression levels (black) [ 4 of 4 ]

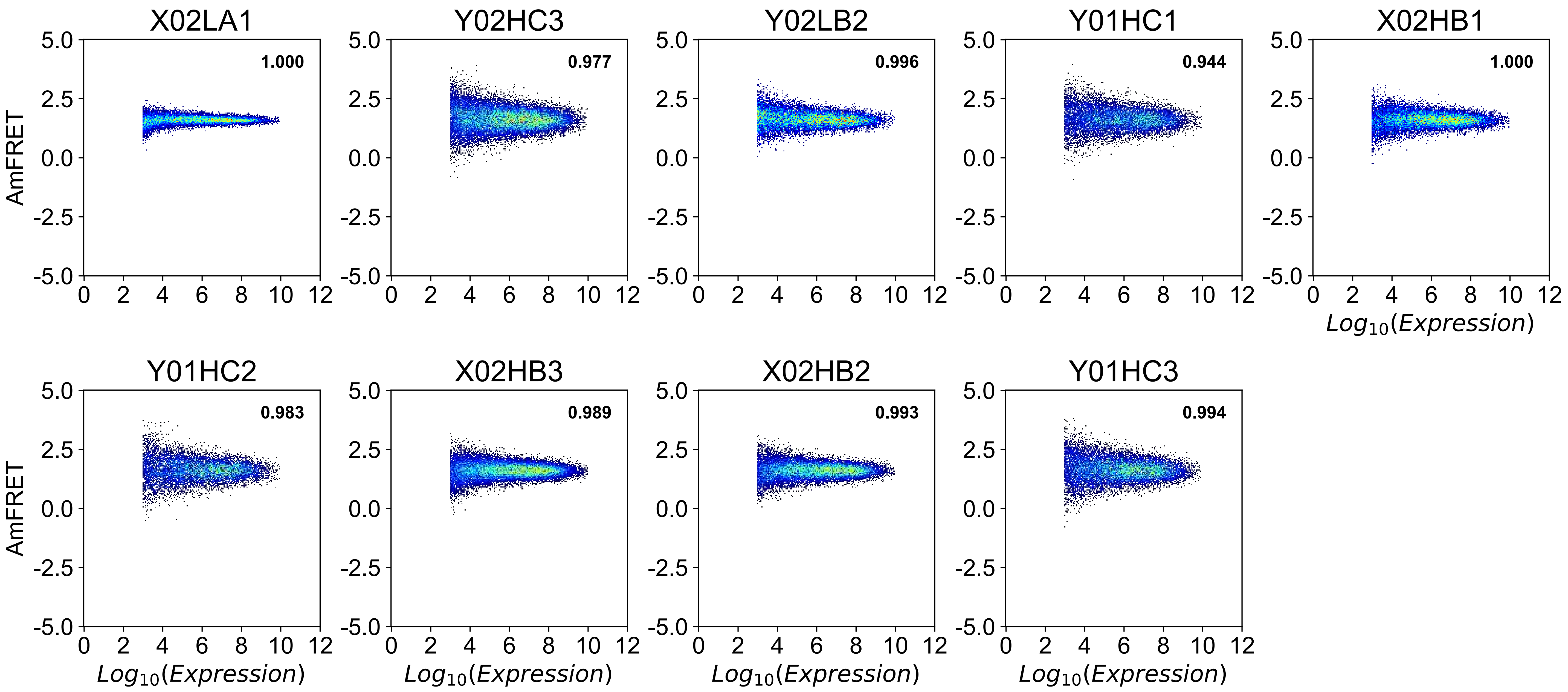

No assembly at all expression levels (blue) [ 1 of 2 ]

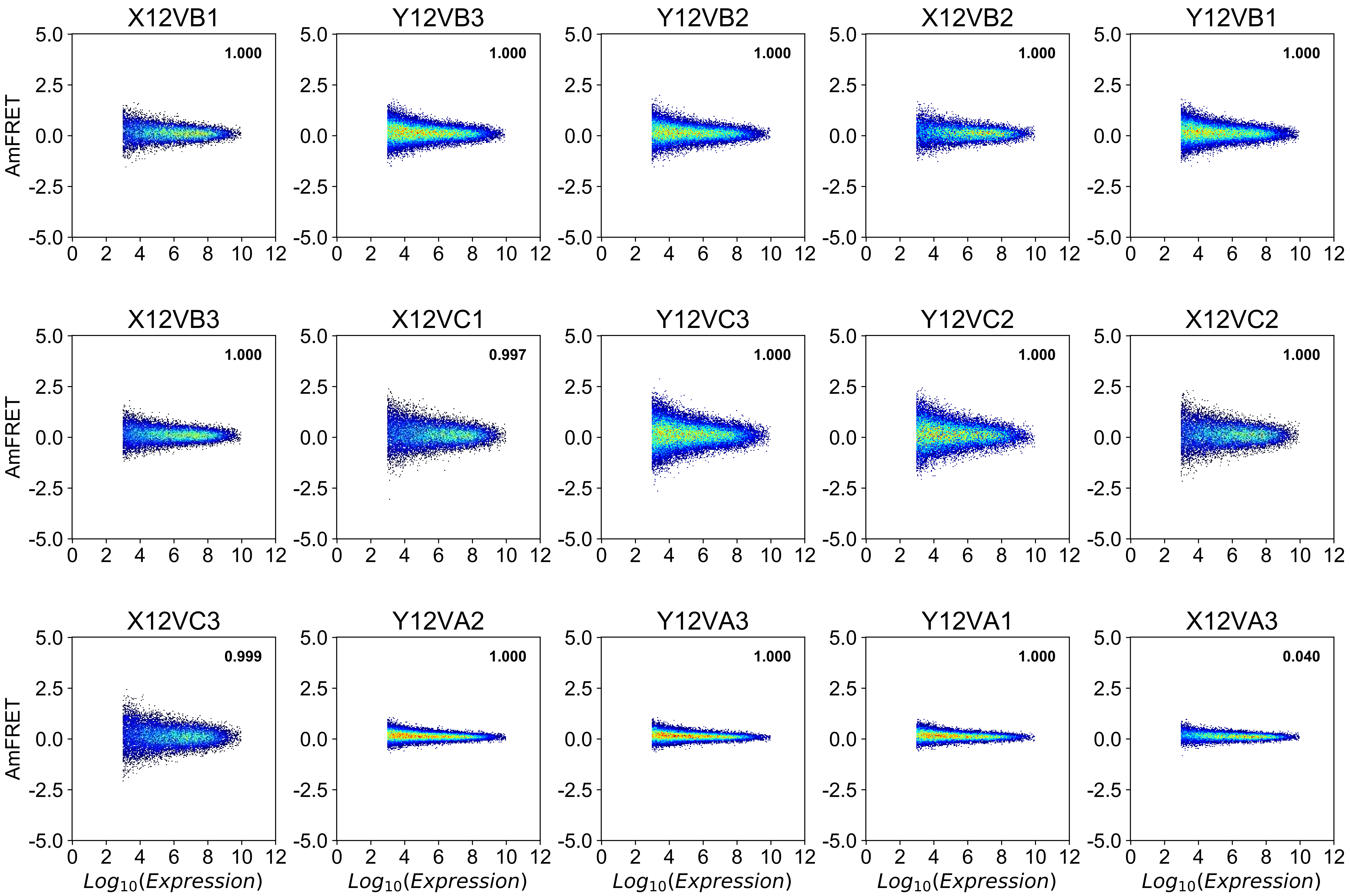

### No assembly at all expression levels (blue) [ 2 of 2 ]

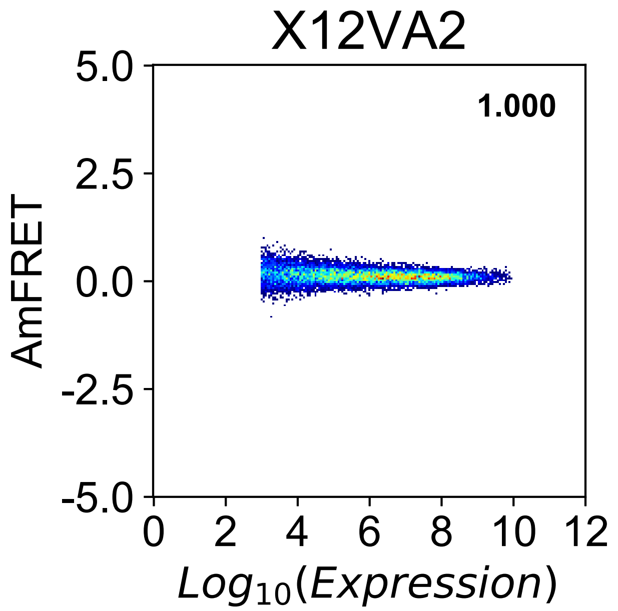

Two-state discontinuous transition (green) [ 1 of 9 ]

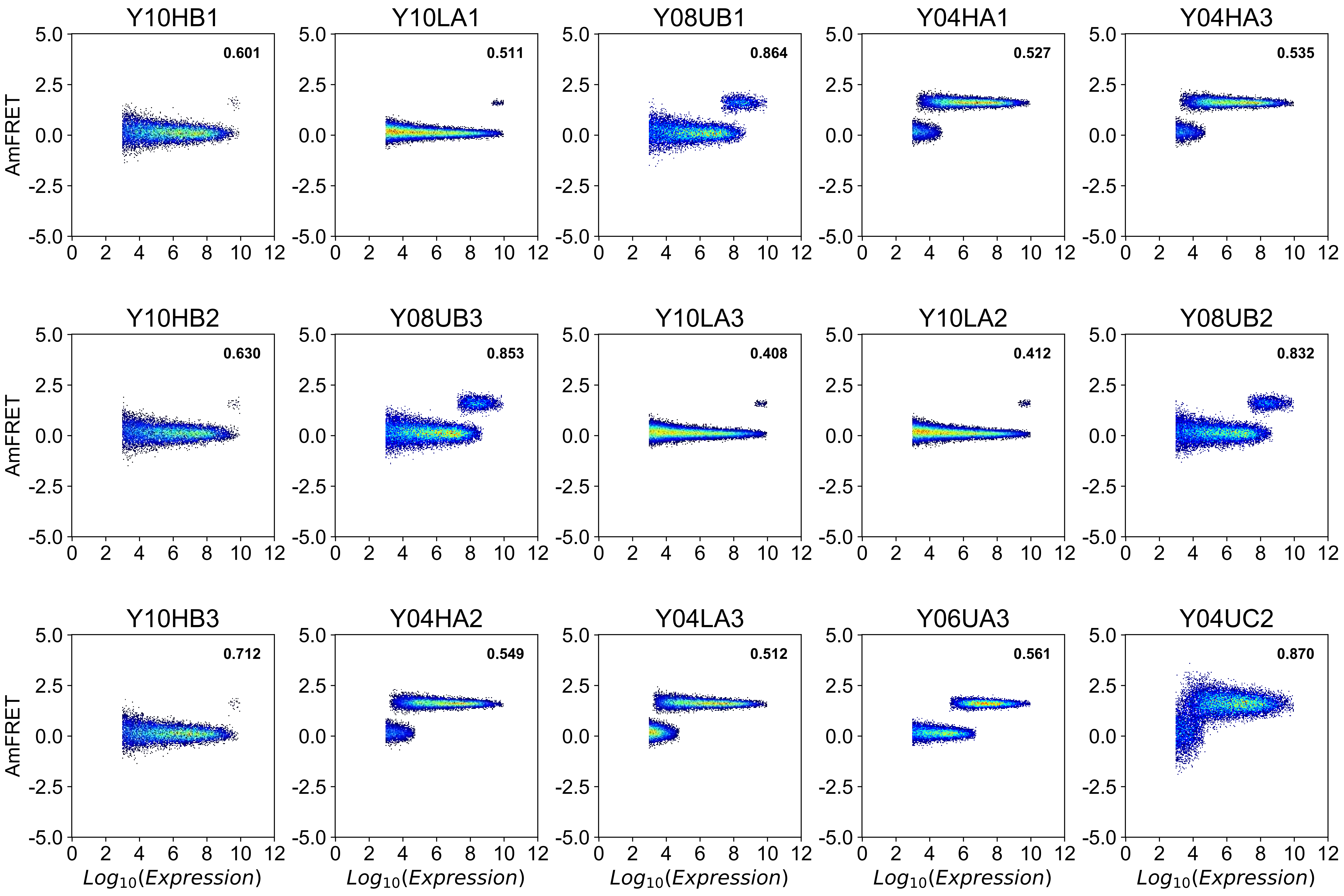

### Two-state discontinuous transition (green) [ 2 of 9 ]

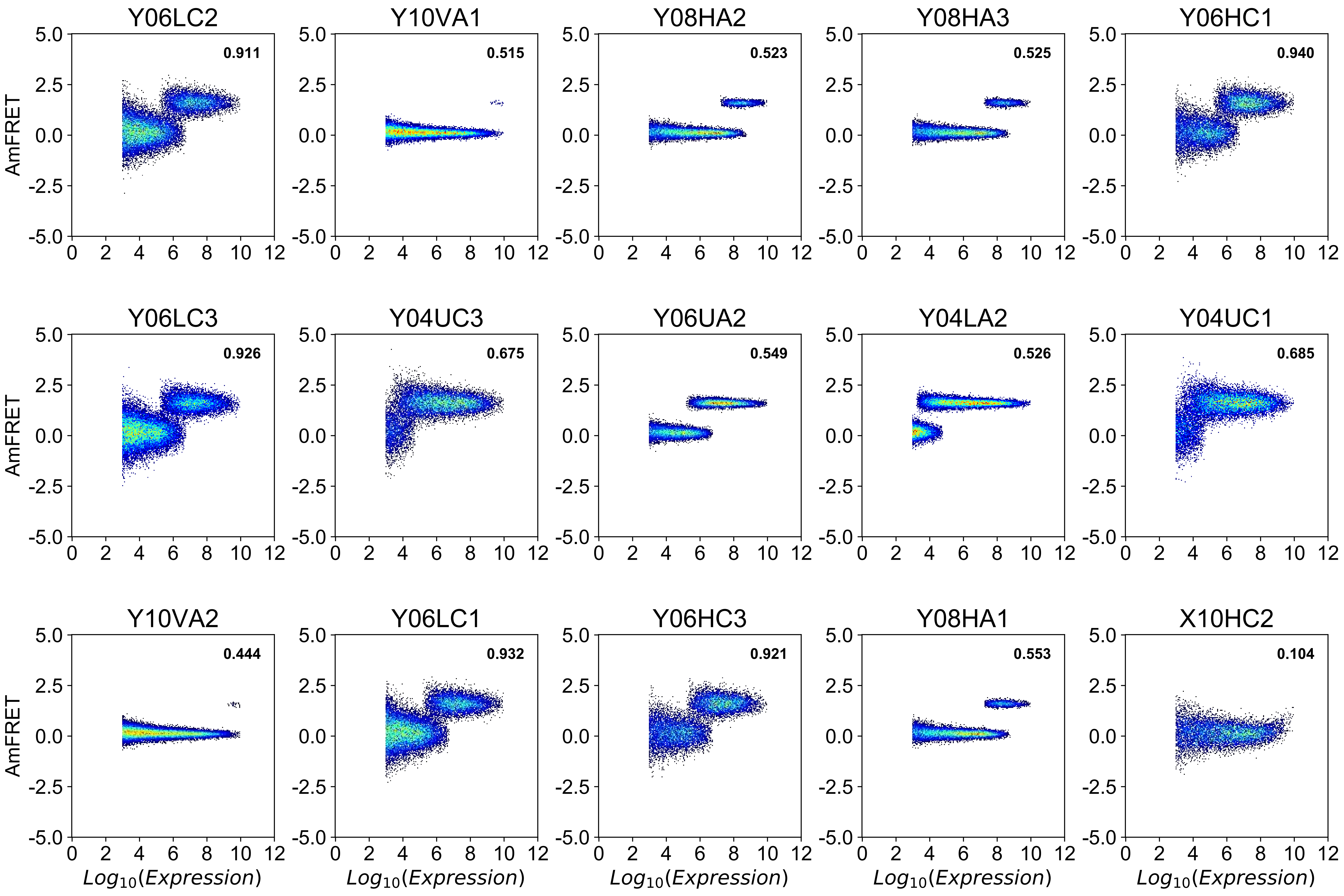

Two-state discontinuous transition (green) [ 3 of 9 ]

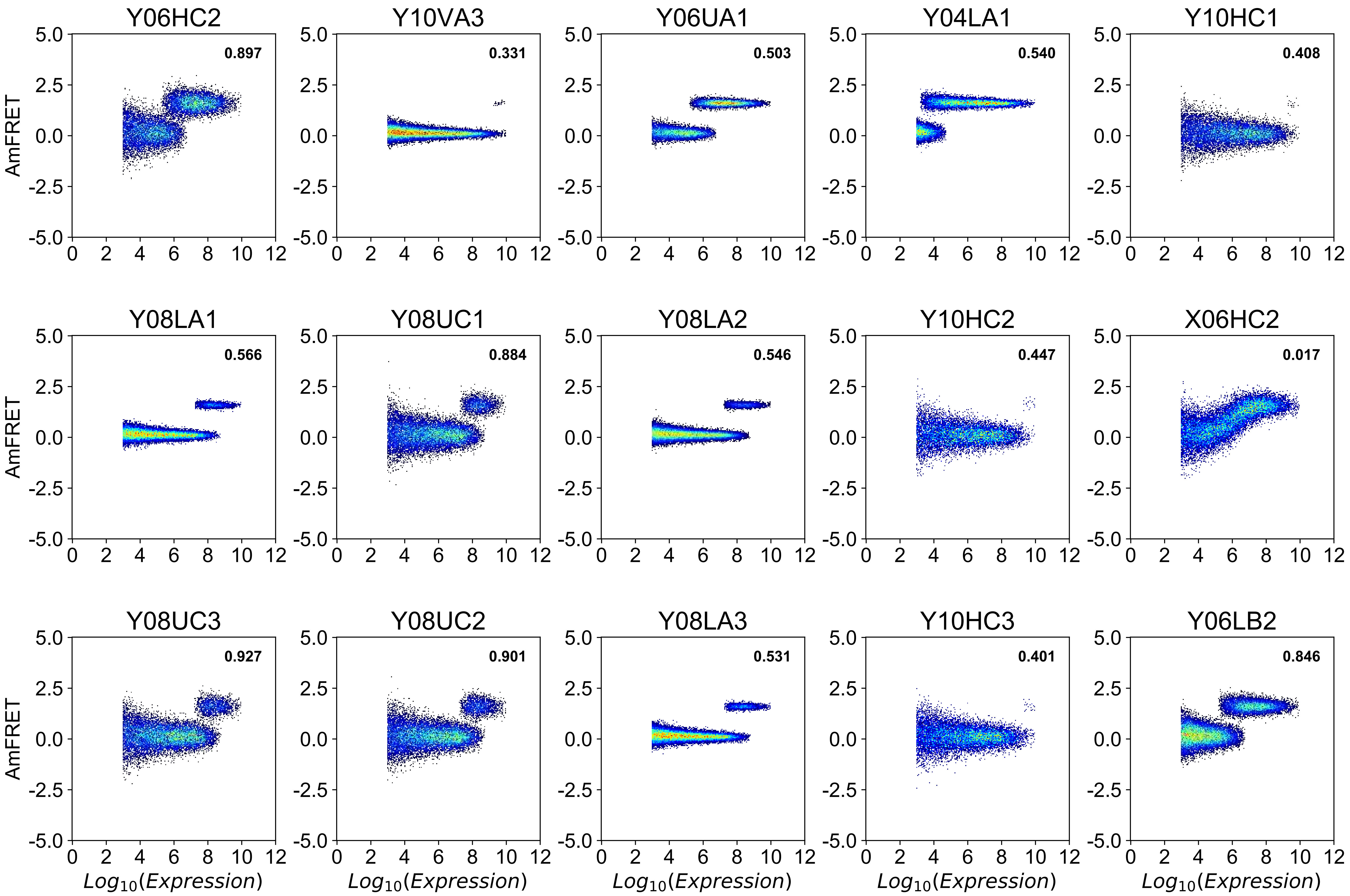

Two-state discontinuous transition (green) [ 4 of 9 ]

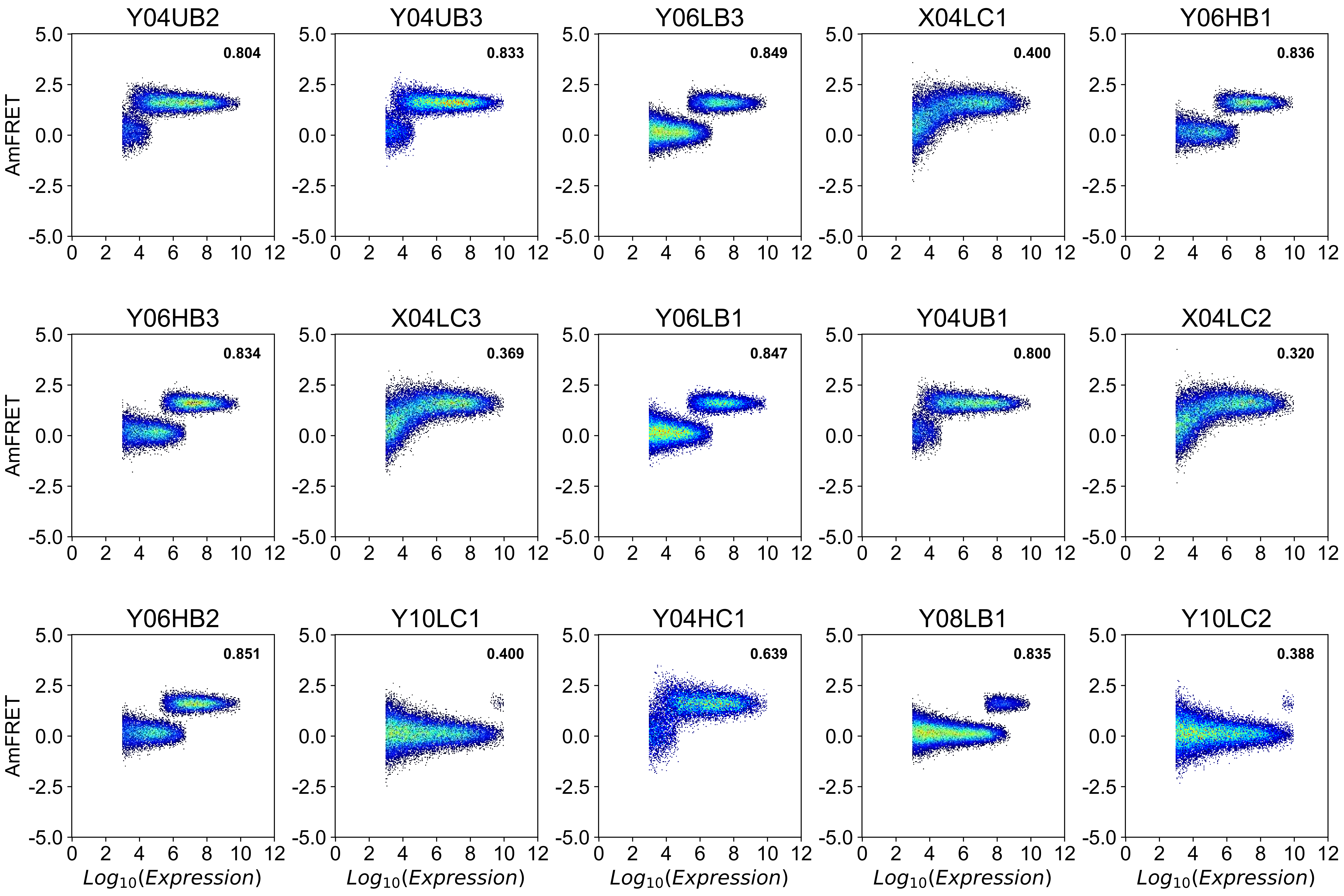

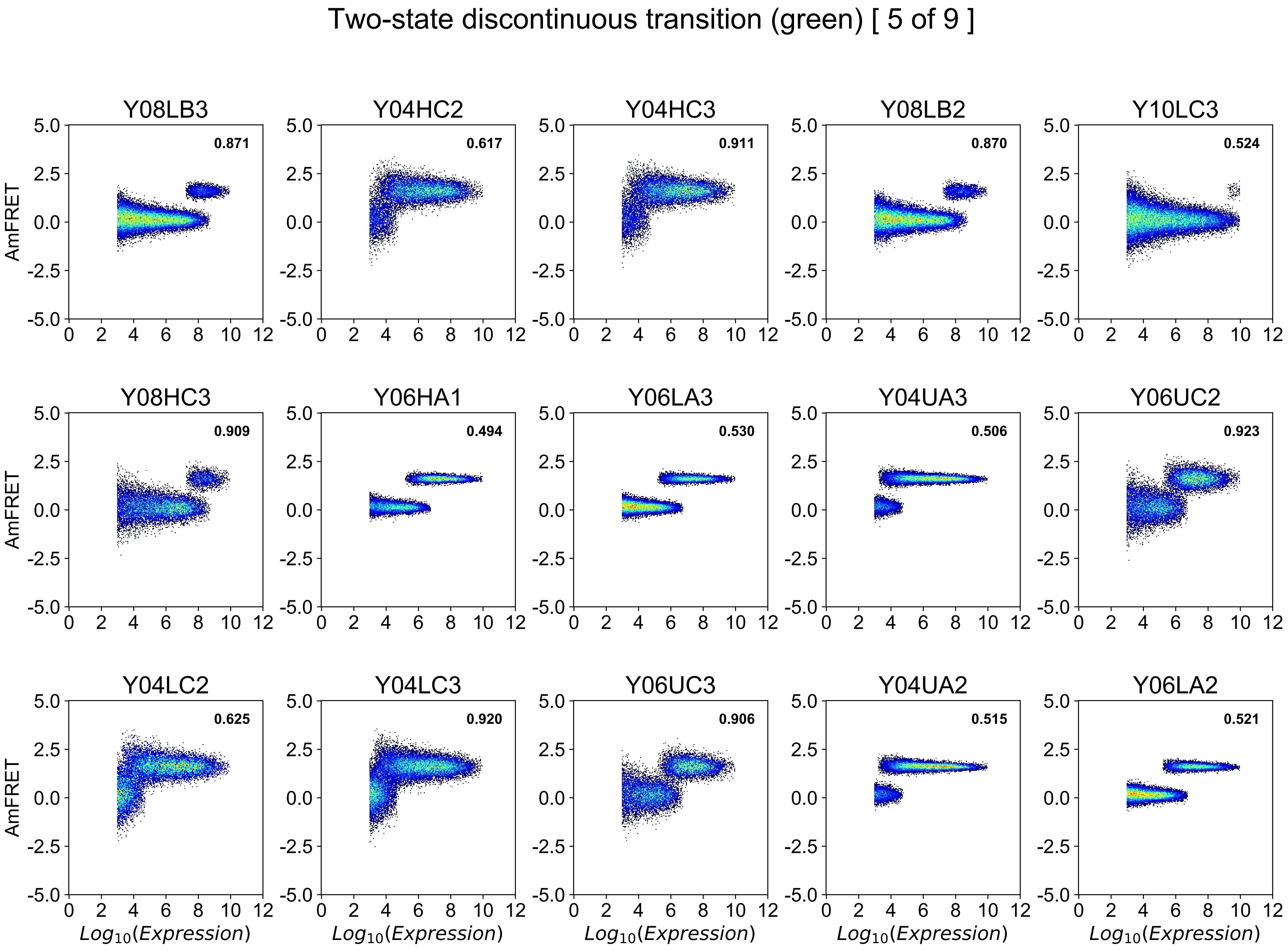

Two-state discontinuous transition (green) [ 5 of 9 ]
Y08LB3
0.871
Y04HC2
0.617
Y04HC3
0.911
Y08LB2
0.870
Y10LC3
0.524
Y08HC3
0.909
Y06HA1
0.494
Y06LA3
0.530
Y04UA3
0.506
Y06UC2
0.923
Y04LC2
0.625
Y04LC3
0.920
Y06UC3
0.906
Y04UA2
0.515
Y06LA2
0.521
AmFRET
Log10(Expression)

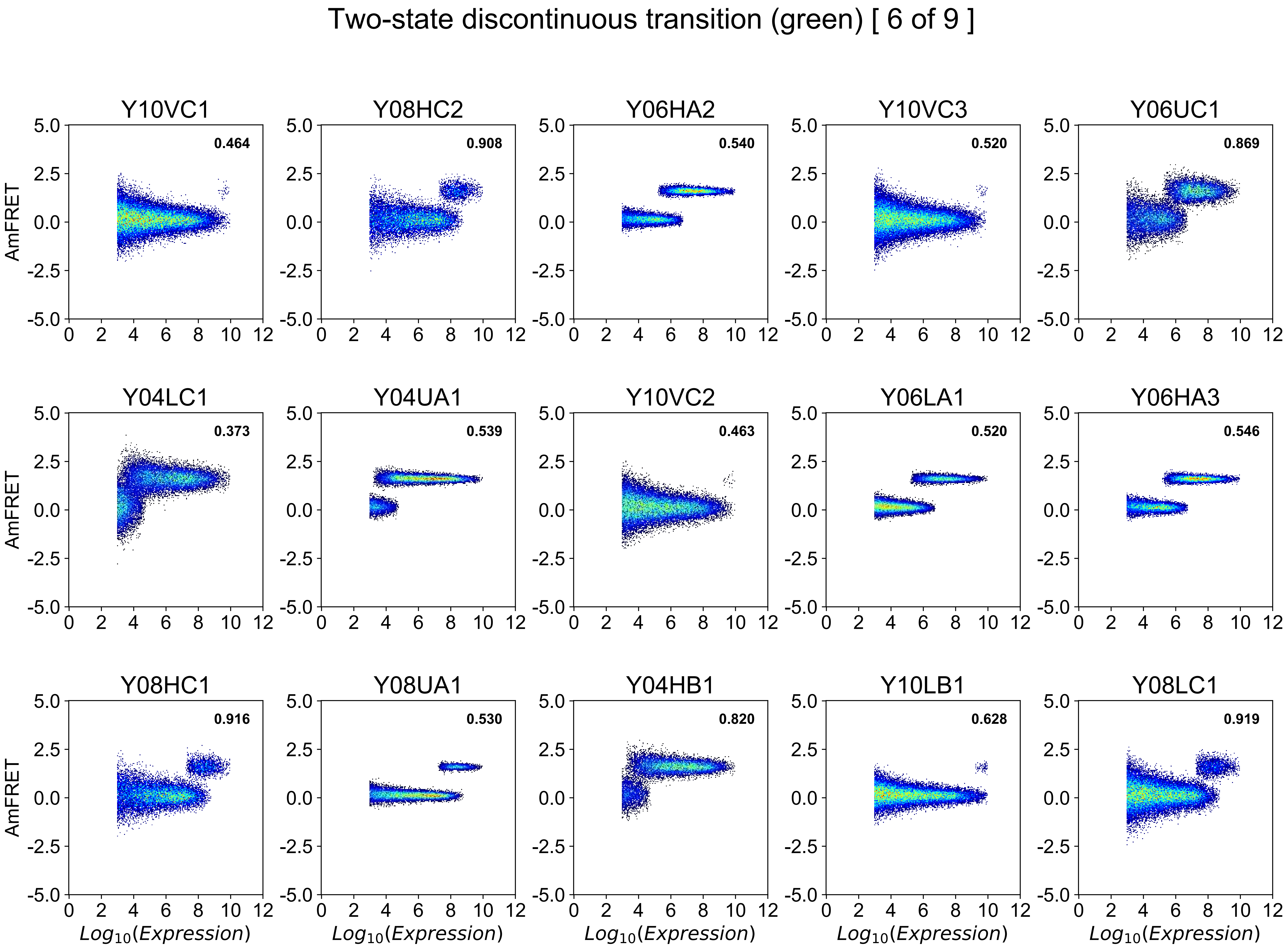
Two-state discontinuous transition (green) [ 6 of 9 ]
Y10VC1
0.464
Y08HC2
0.908
Y06HA2
0.540
Y10VC3
0.520
Y06UC1
0.869
Y04LC1
0.373
Y04UA1
0.539
Y10VC2
0.463
Y06LA1
0.520
Y06HA3
0.546
Y08HC1
0.916
Y08UA1
0.530
Y04HB1
0.820
Y10LB1
0.628
Y08LC1
0.919
AmFRET
Log10(Expression)

Two-state discontinuous transition (green) [ 7 of 9 ]

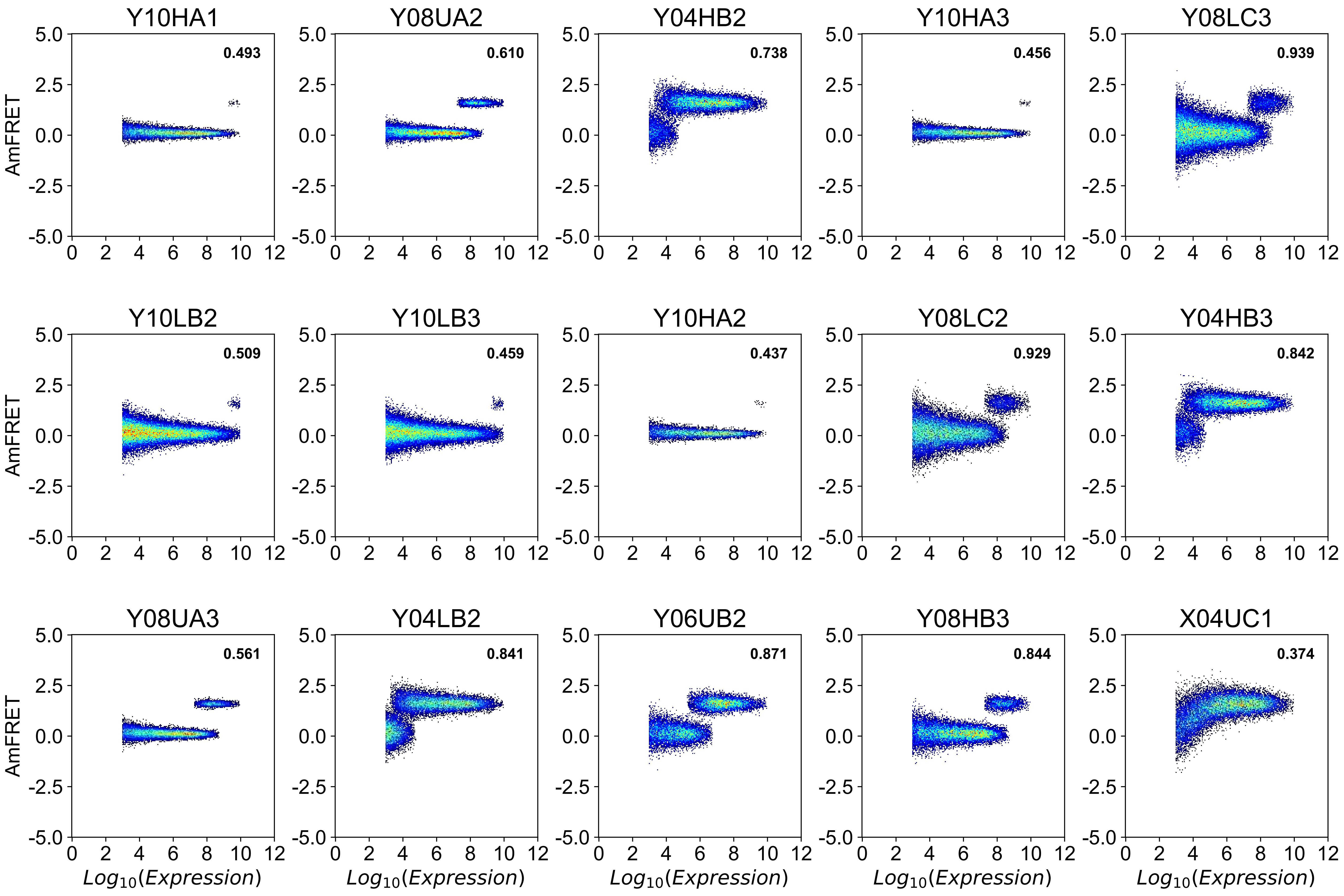

### Two-state discontinuous transition (green) [ 8 of 9 ]

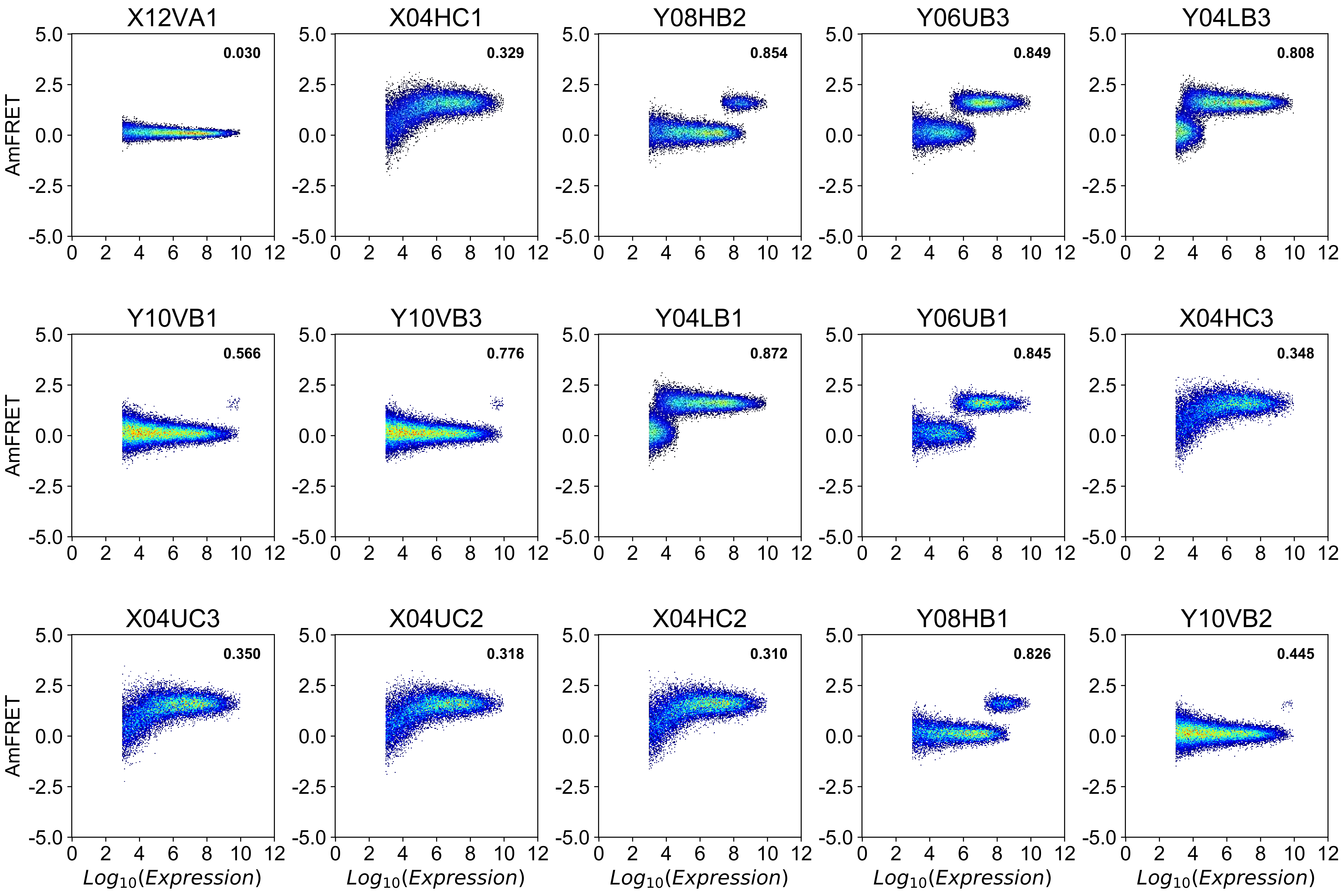

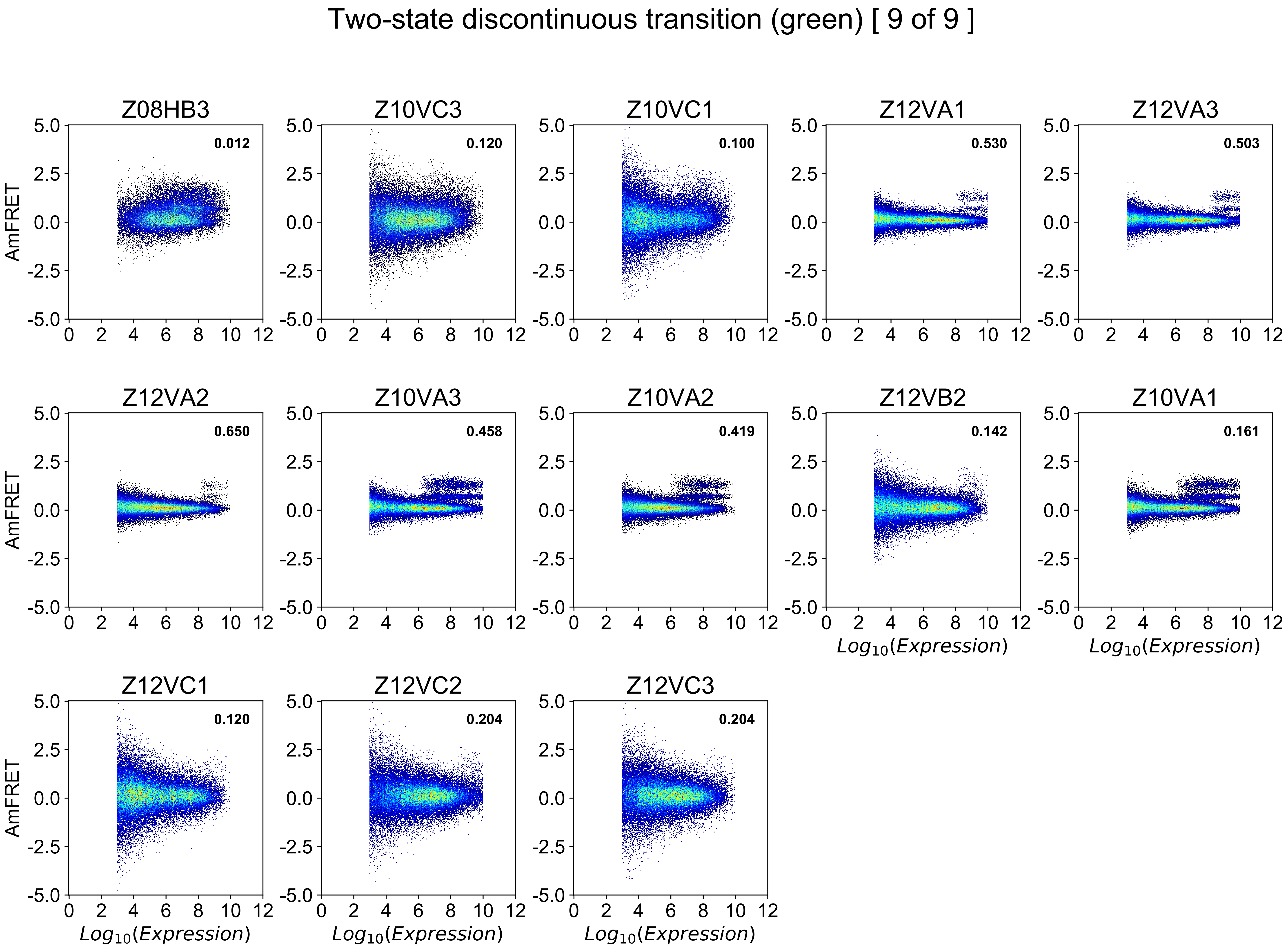

Two-state discontinuous transition (green) [ 9 of 9 ]
Z08HB3
0.012
Z10VC3
0.120
Z10VC1
0.100
Z12VA1
0.530
Z12VA3
0.503
Z12VA2
0.650
Z10VA3
0.458
Z10VA2
0.419
Z12VB2
0.142
Z10VA1
0.161
Z12VC1
0.120
Z12VC2
0.204
Z12VC3
0.204
AmFRET
$Log_{10}(Expression)$

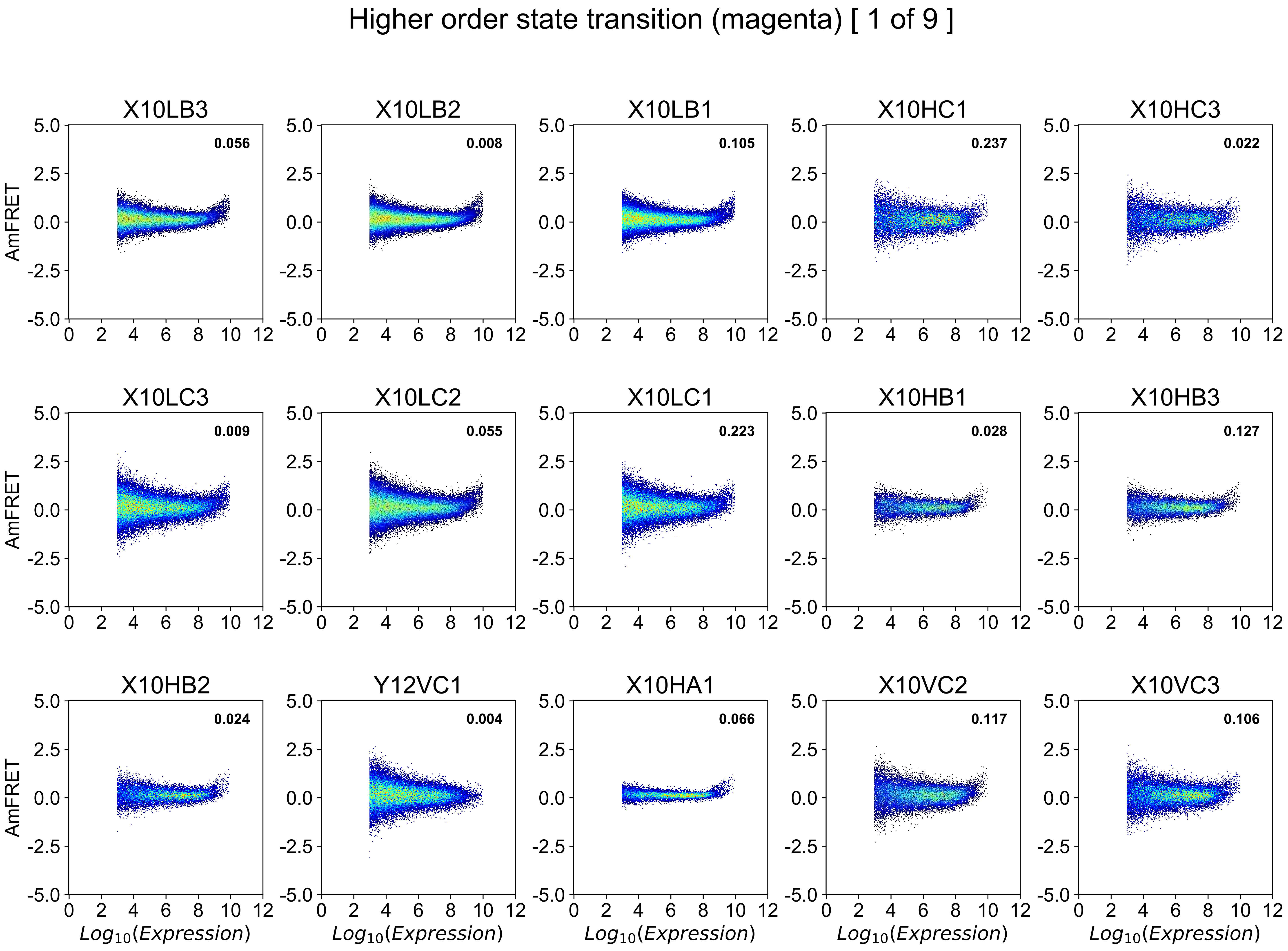
Higher order state transition (magenta) [ 1 of 9 ]
X10LB3
0.056
X10LB2
0.008
X10LB1
0.105
X10HC1
0.237
X10HC3
0.022
X10LC3
0.009
X10LC2
0.055
X10LC1
0.223
X10HB1
0.028
X10HB3
0.127
X10HB2
0.024
Y12VC1
0.004
X10HA1
0.066
X10VC2
0.117
X10VC3
0.106
AmFRET
Log10(Expression)

### Higher order state transition (magenta) [ 2 of 9 ]

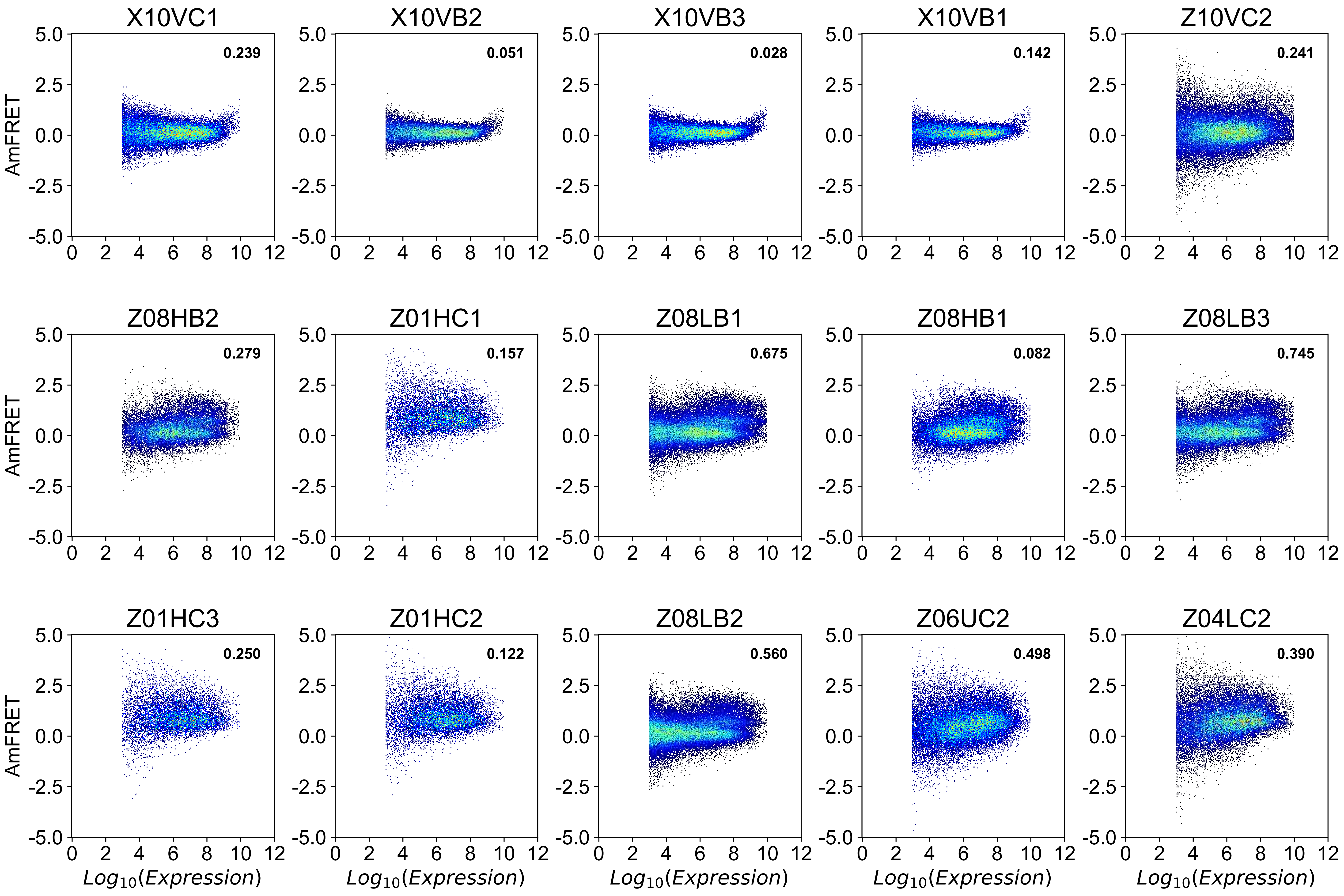

### Higher order state transition (magenta) [ 3 of 9 ]

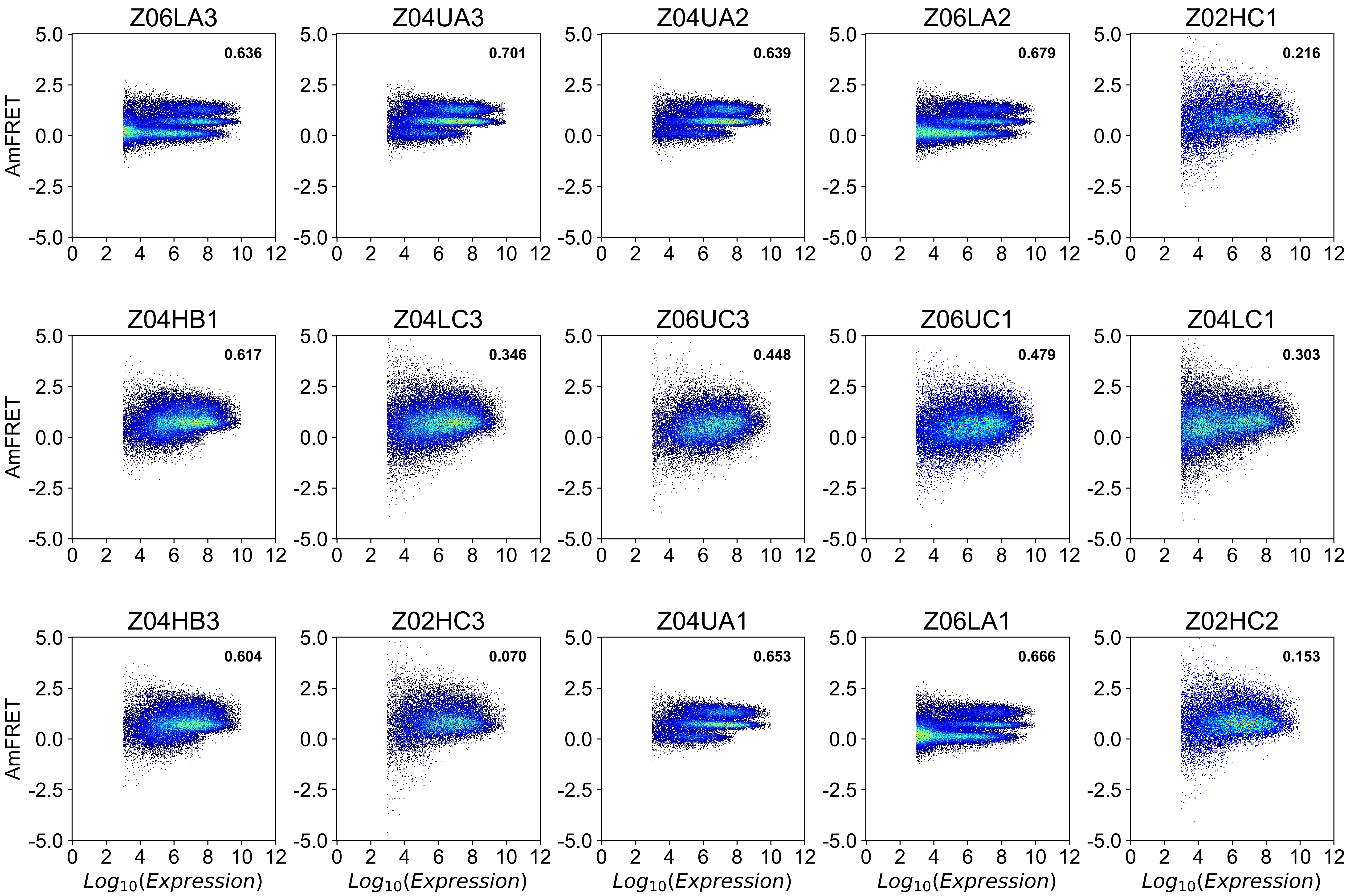

#### Higher order state transition (magenta) [ 4 of 9 ]

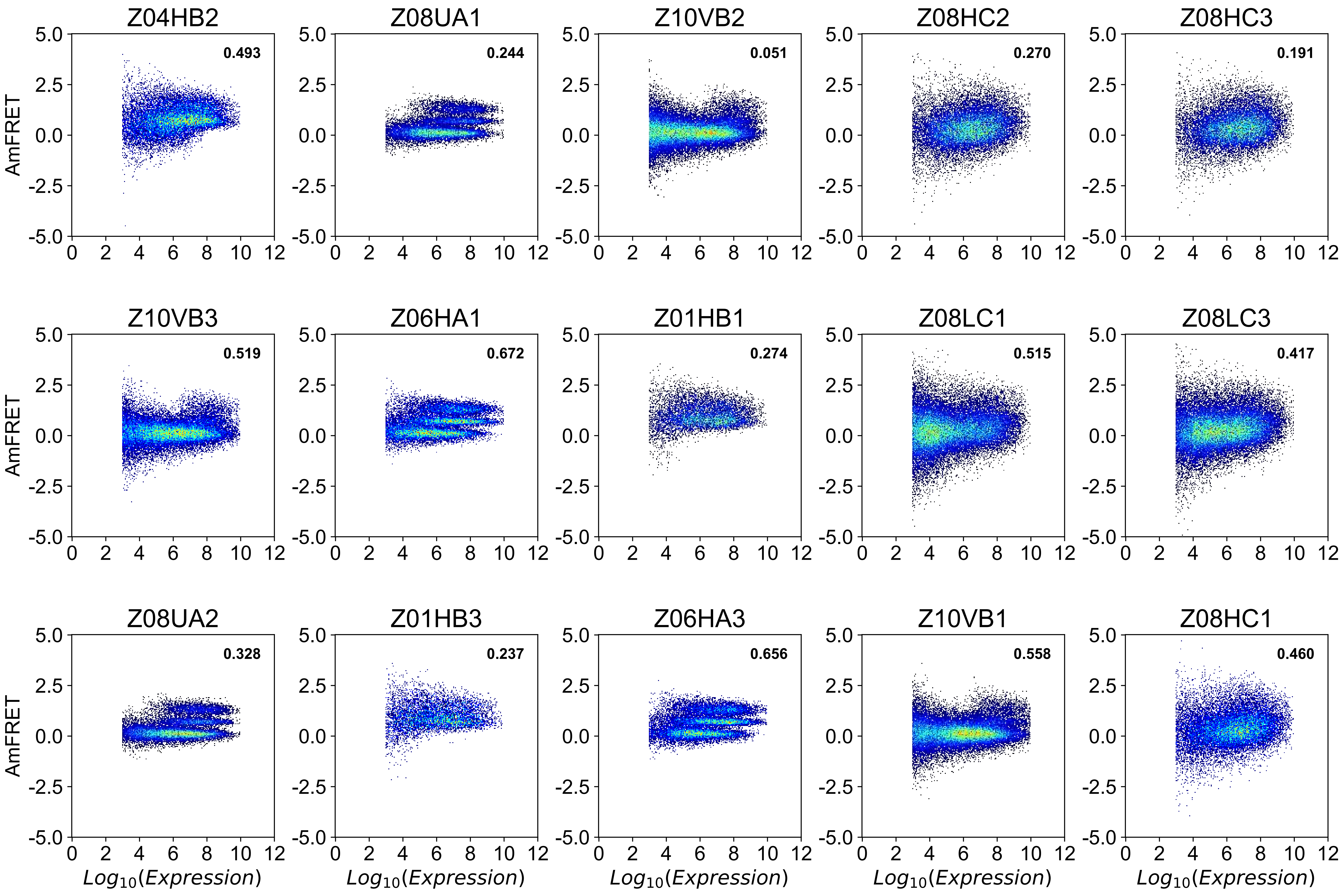

### Higher order state transition (magenta) [ 5 of 9 ]

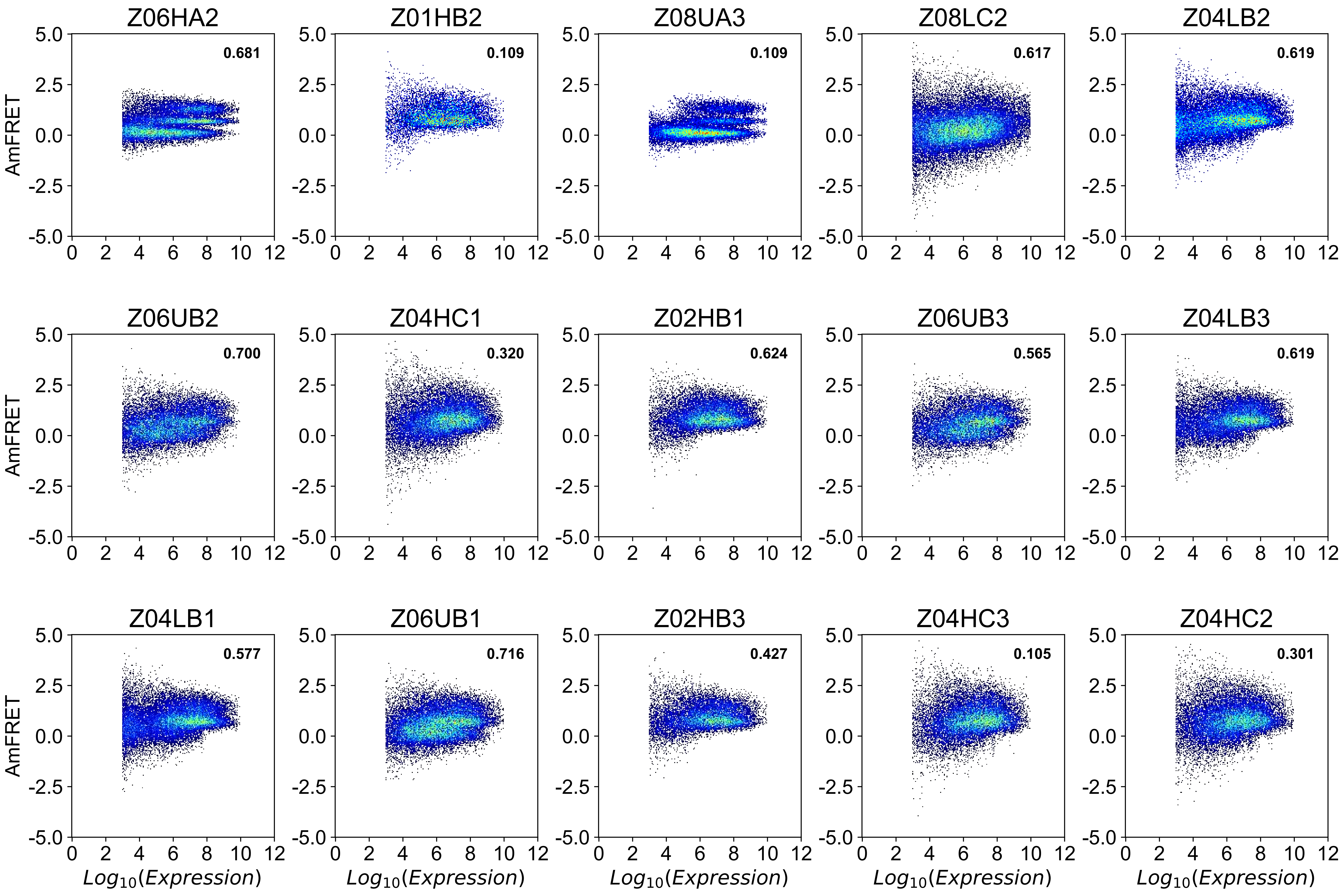

### Higher order state transition (magenta) [ 6 of 9 ]

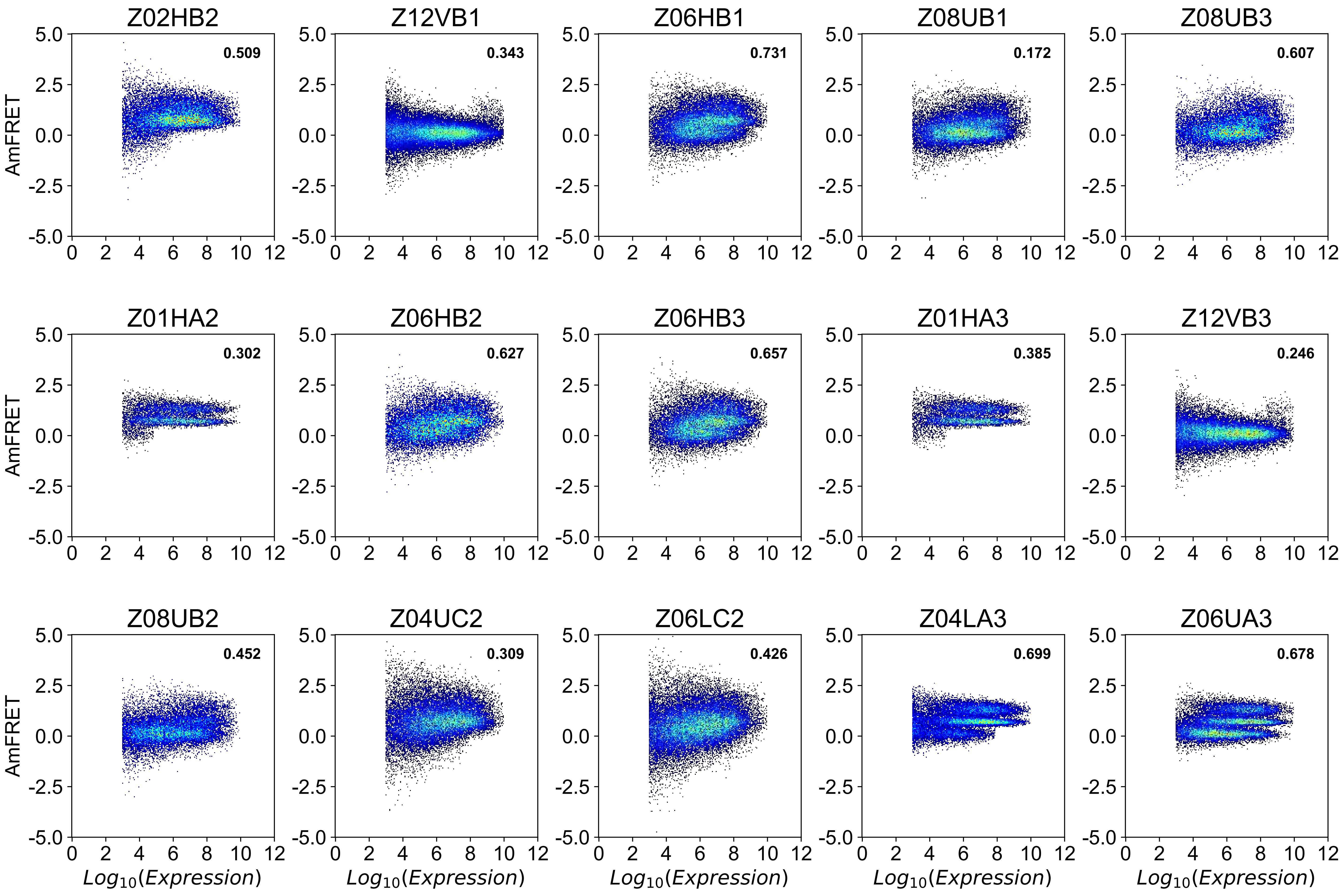

### Higher order state transition (magenta) [ 7 of 9 ]

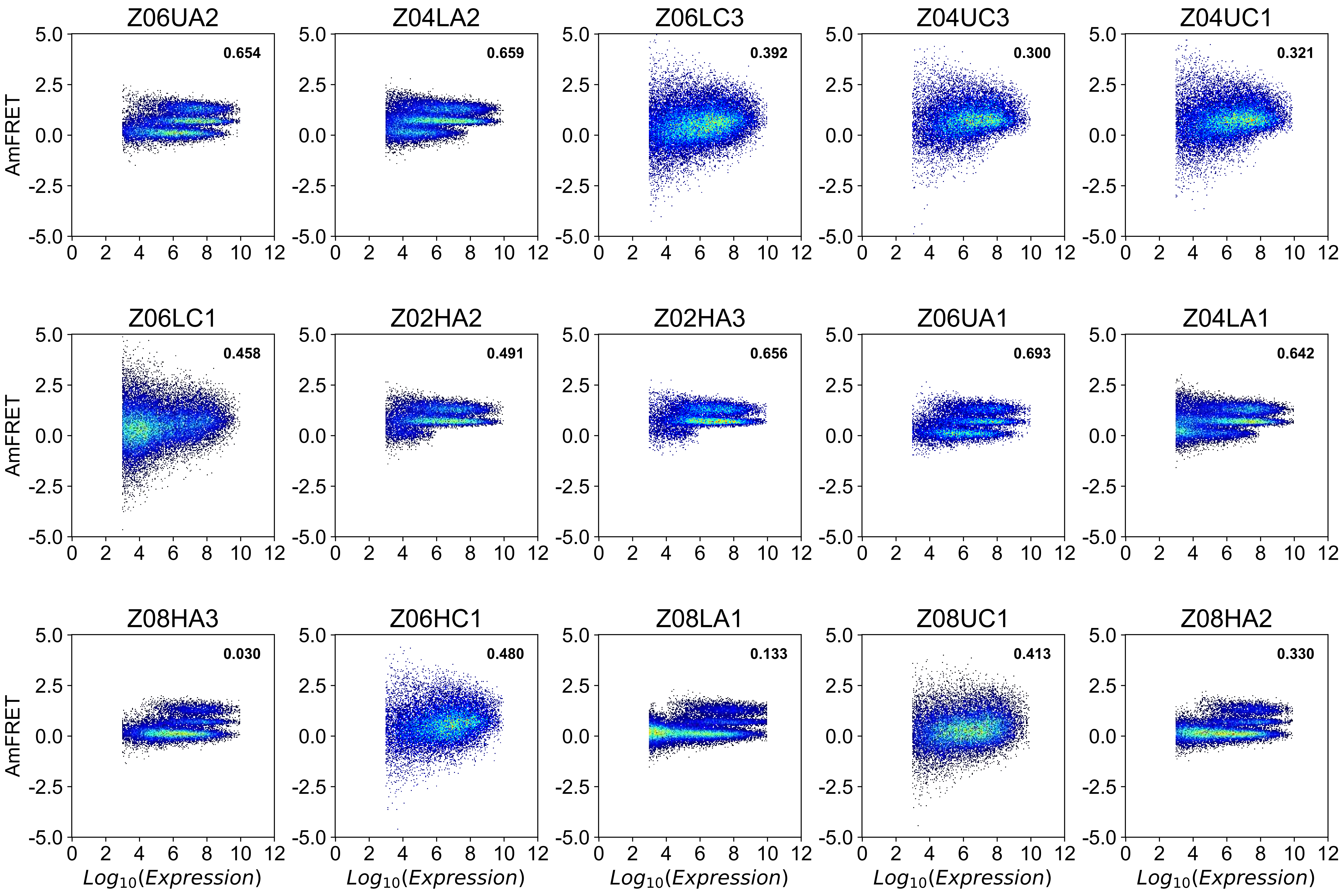

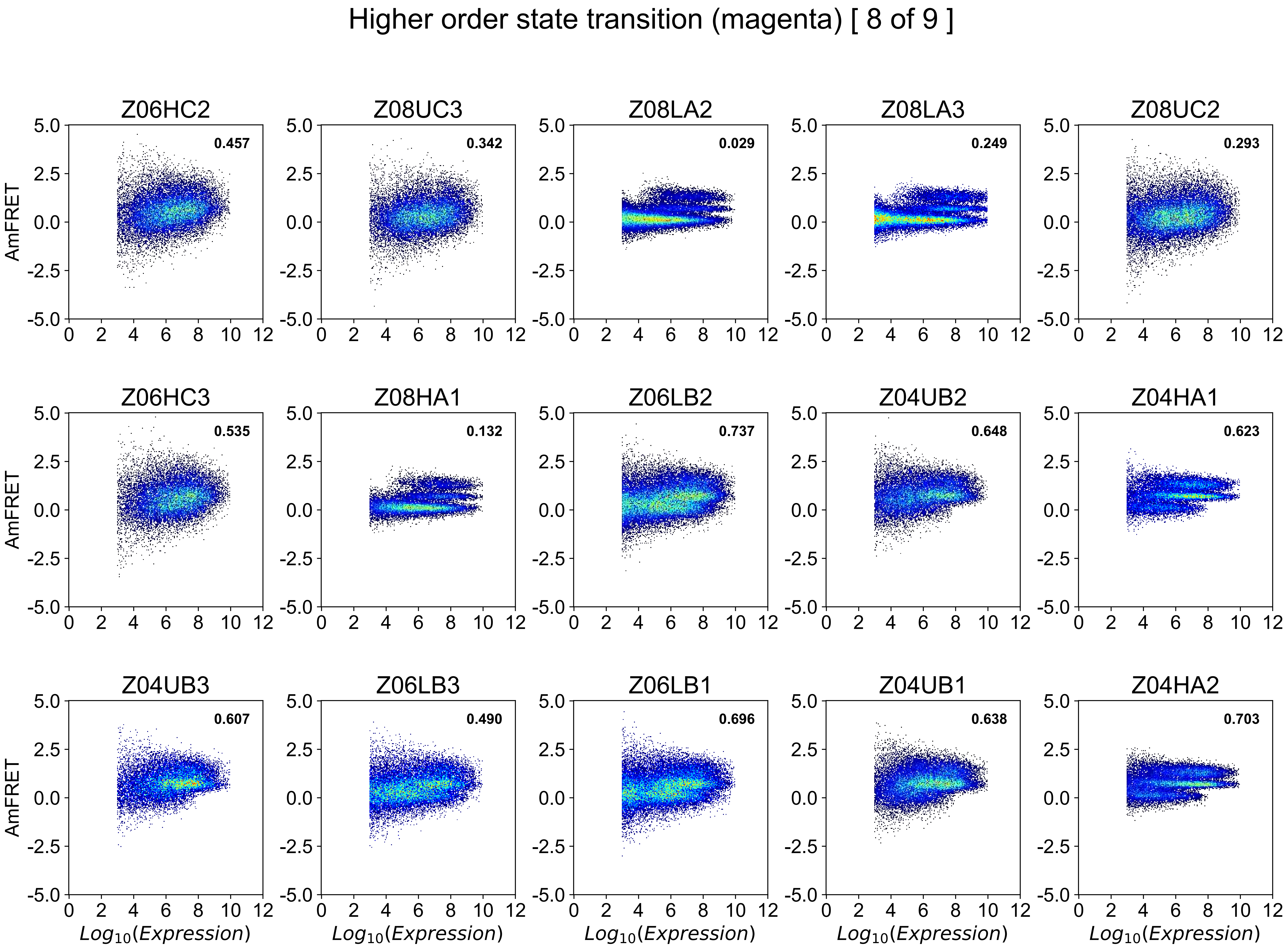
Higher order state transition (magenta) [ 8 of 9 ]
Z06HC2
0.457
Z08UC3
0.342
Z08LA2
0.029
Z08LA3
0.249
Z08UC2
0.293
Z06HC3
0.535
Z08HA1
0.132
Z06LB2
0.737
Z04UB2
0.648
Z04HA1
0.623
Z04UB3
0.607
Z06LB3
0.490
Z06LB1
0.696
Z04UB1
0.638
Z04HA2
0.703
AmFRET
Log10(Expression)
5.0
2.5
0.0
-2.5
-5.0
0
2
4
6
8
10
12

### Higher order state transition (magenta) [ 9 of 9 ]

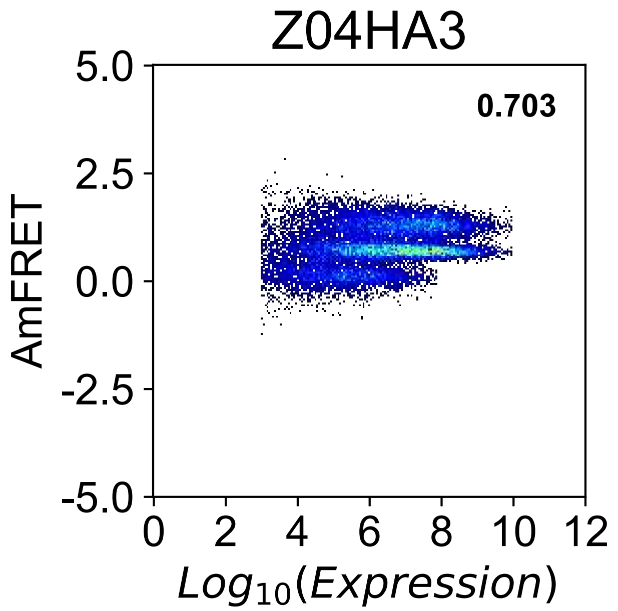

Two-state continuous transition (red) [ 1 of 6 ]

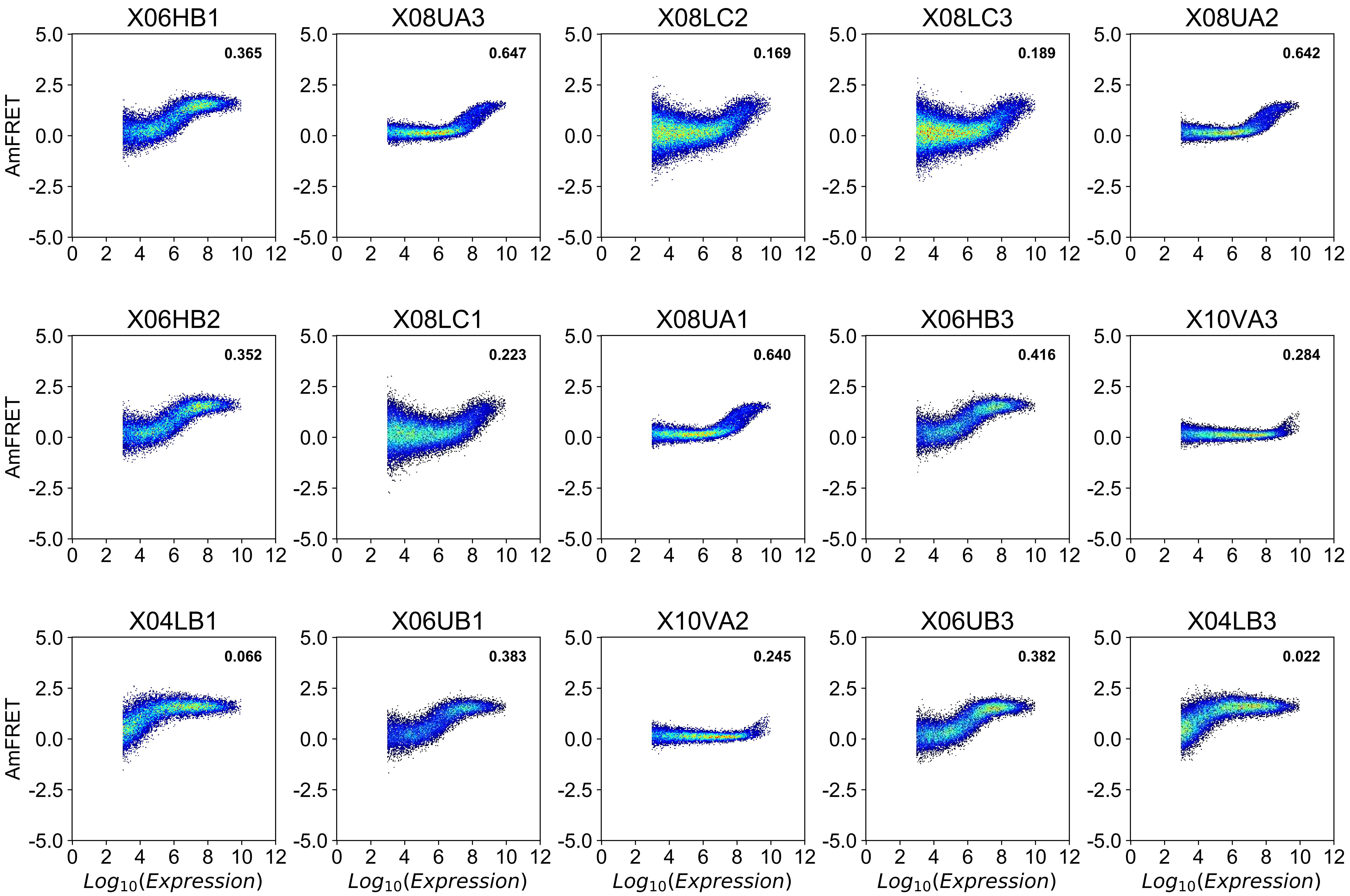

#### Two-state continuous transition (red) [ 2 of 6 ]

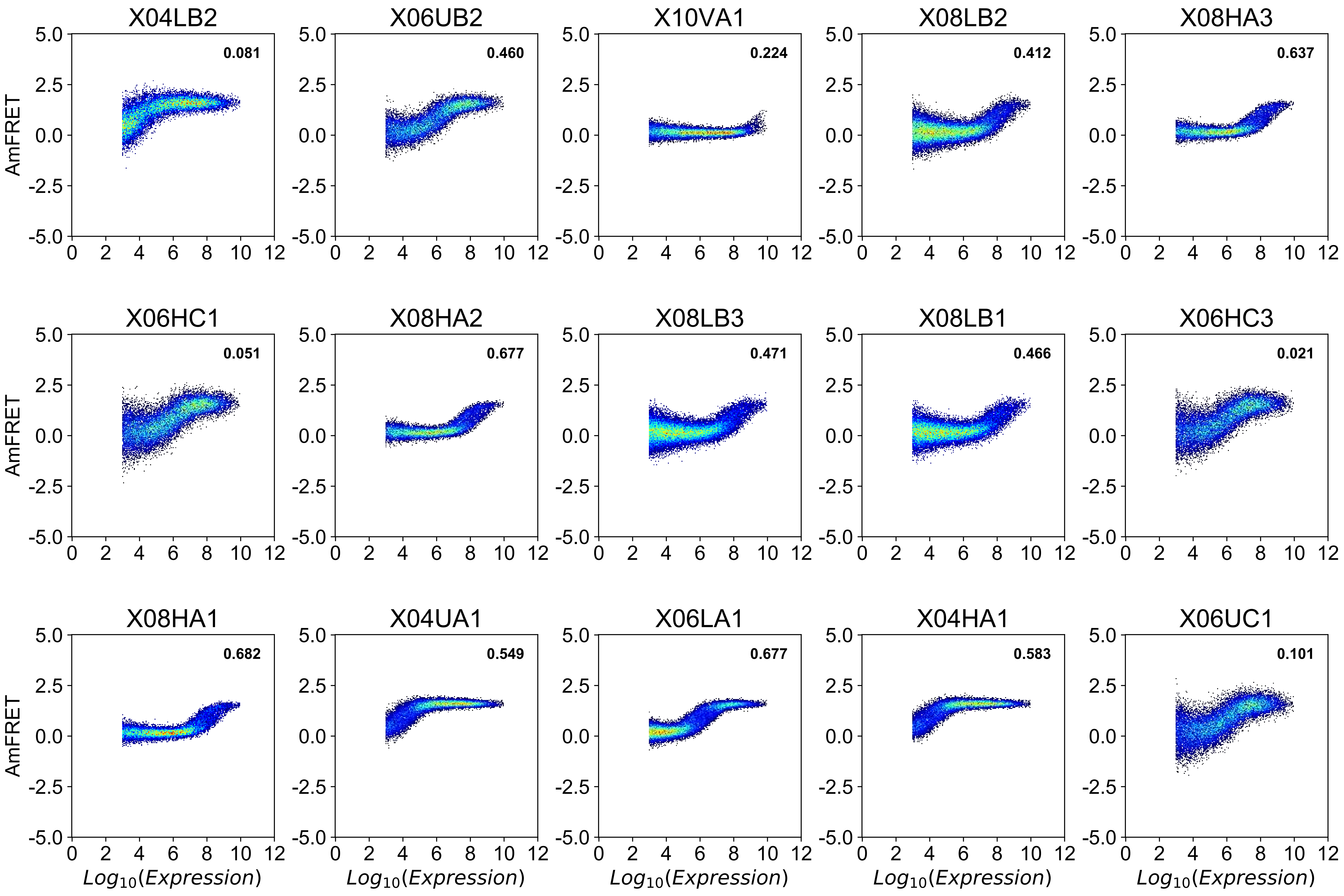

### Two-state continuous transition (red) [ 3 of 6 ]

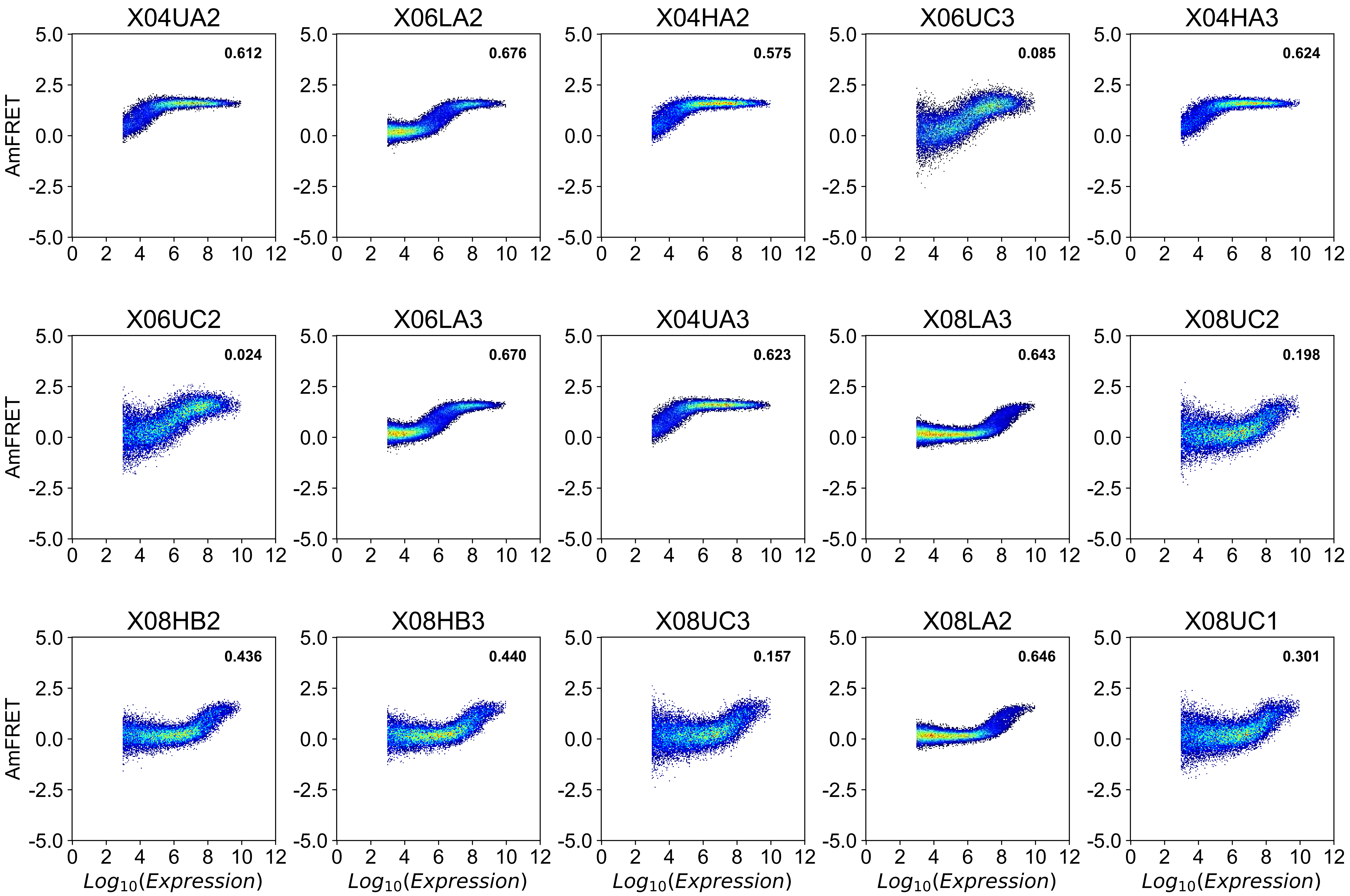

Two-state continuous transition (red) [ 4 of 6 ]

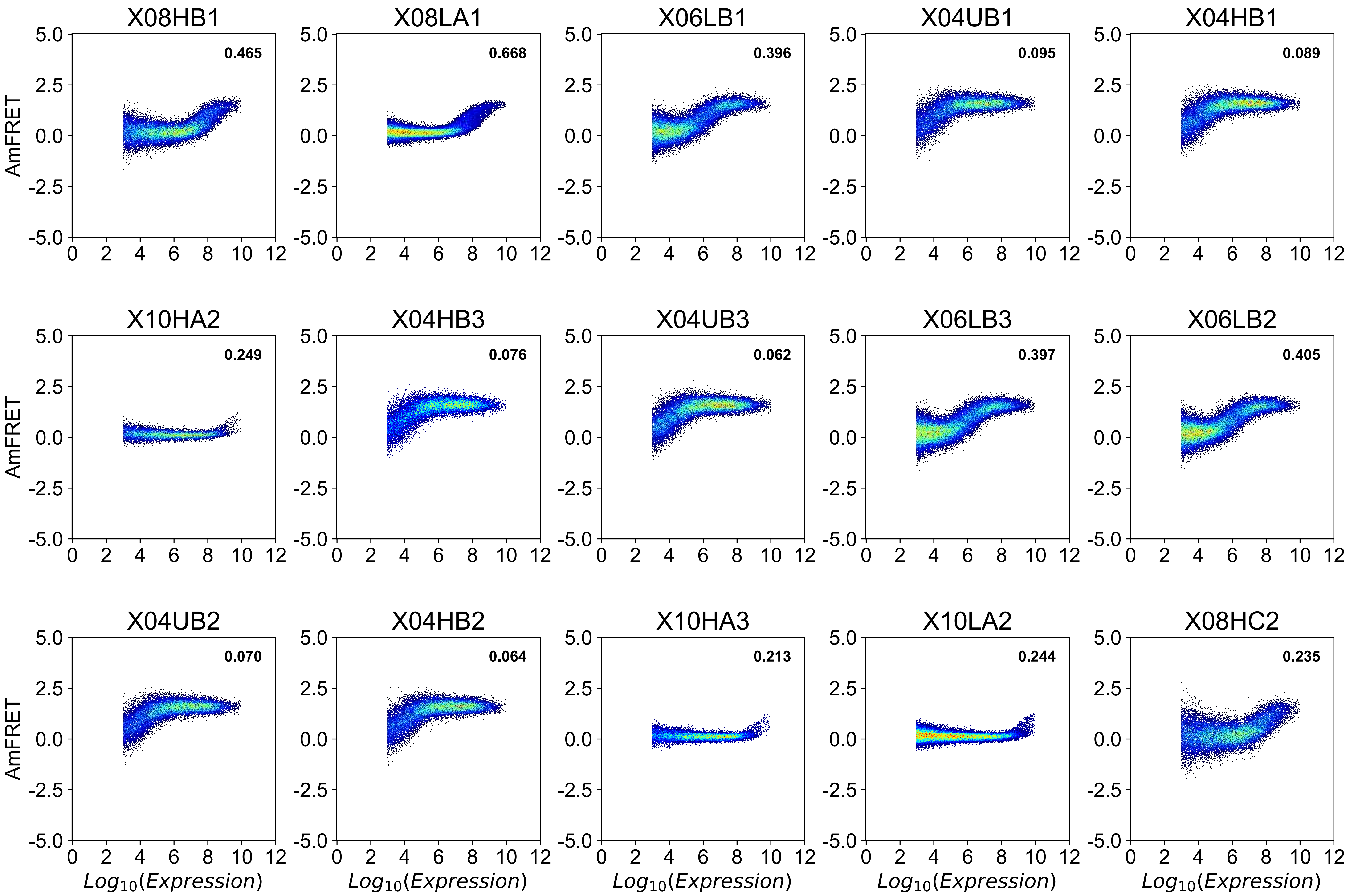

Two-state continuous transition (red) [ 5 of 6 ]

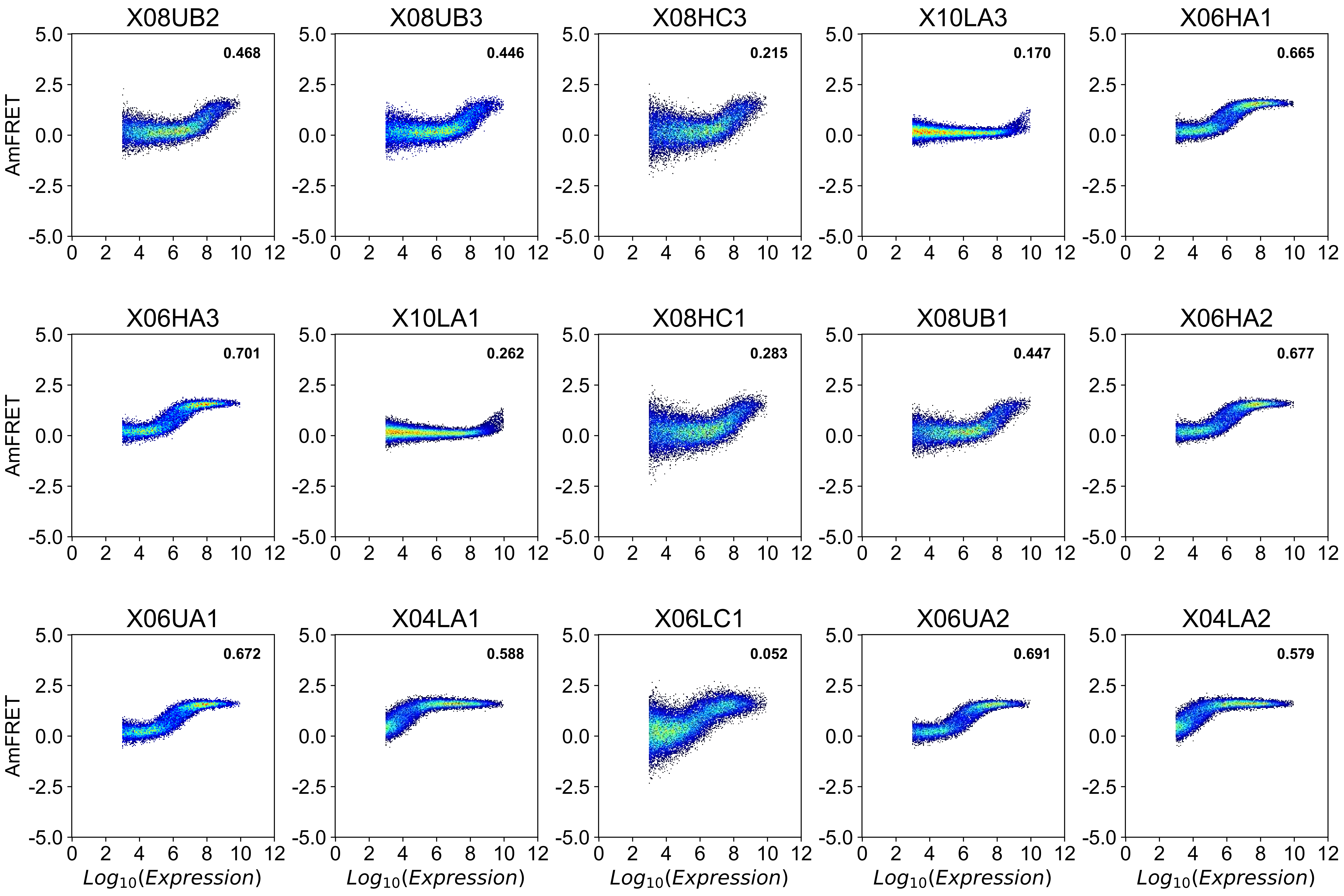

### Two-state continuous transition (red) [ 6 of 6 ]

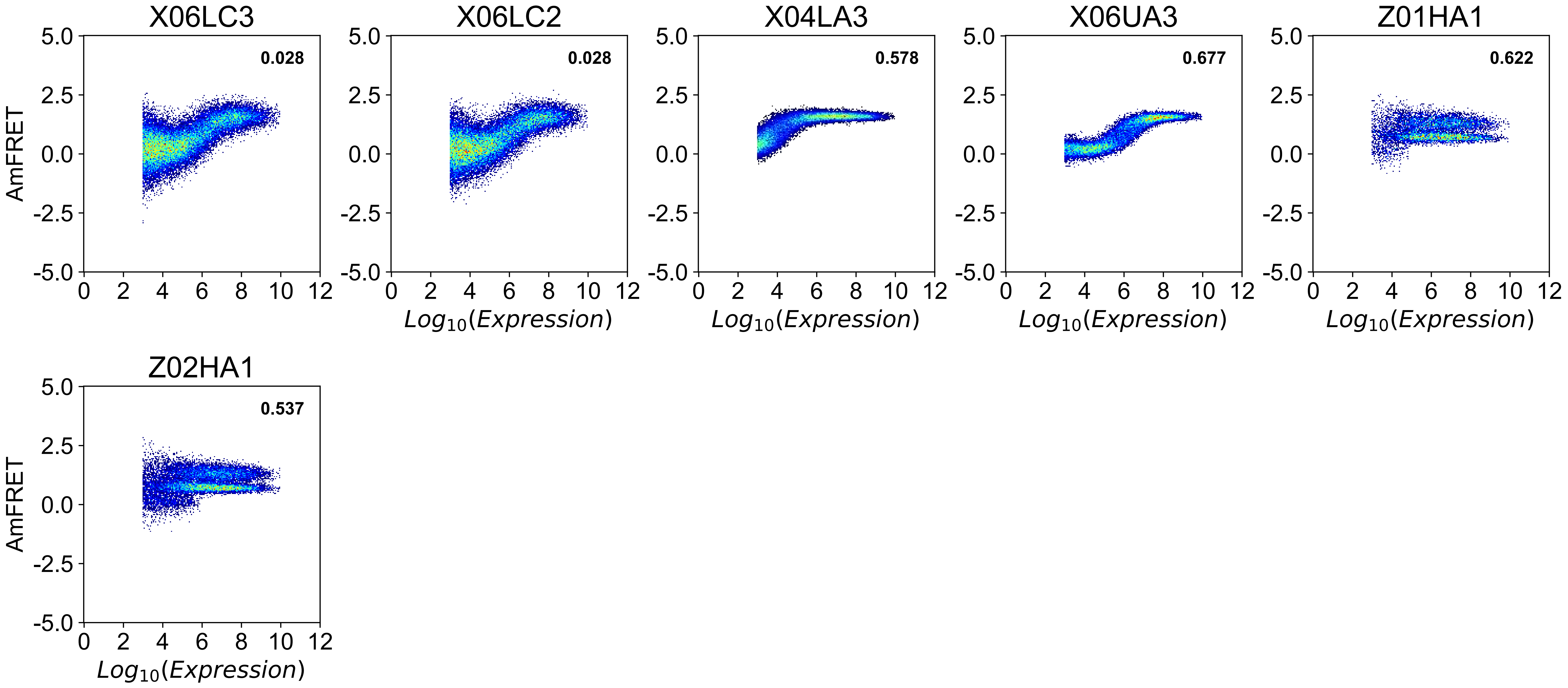

**Figure S3: Classification of synthetic DAmFRET histograms without shading by confidence score**. Each row denotes a different replica. Each column is denoted by the $\log_{10}(c_{50})$ (1.0, 2.0, 4.0, 6.0, 8.0, 10.0, or 12.0), the function (Sigmoid, Step, or 2 Step), and the noise level (Low, Medium, or High) used to generate the synthetic DAmFRET histograms.

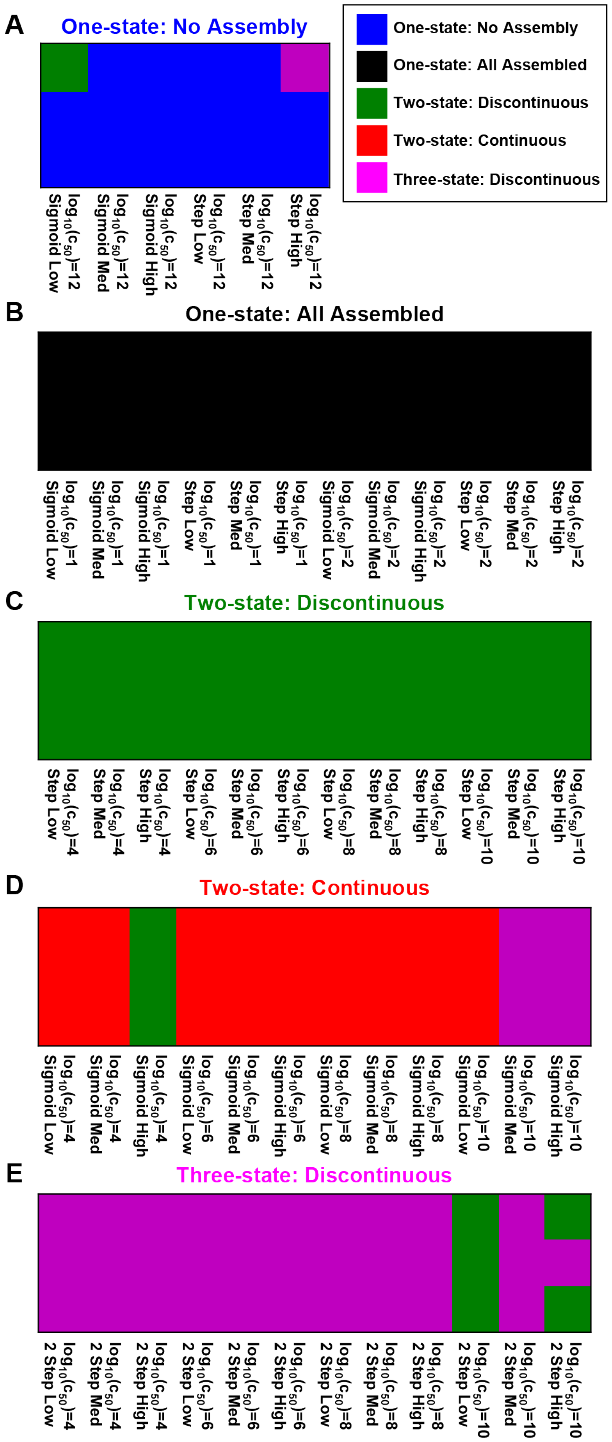

**Figure S4: DAmFRET histograms for each cPrD sorted by their PAPA score.** The border of each histogram is colored by its classification. The following classifications are used: assembled at all expression levels (black); no assembly at all expression levels (blue); continuous two-state transition (red); discontinuous two-state transition (green); higher order state transition (magenta); and infrequent transition (yellow). Also displayed in the inset of each profile is the confidence score of its classification. Although the full profile is plotted, our method utilizes a low cutoff of 1.5 and a high cutoff at 5.0 in $\log_{10}(Expression)$ in our analysis pipeline to reduce the noise contributions at the extrema.

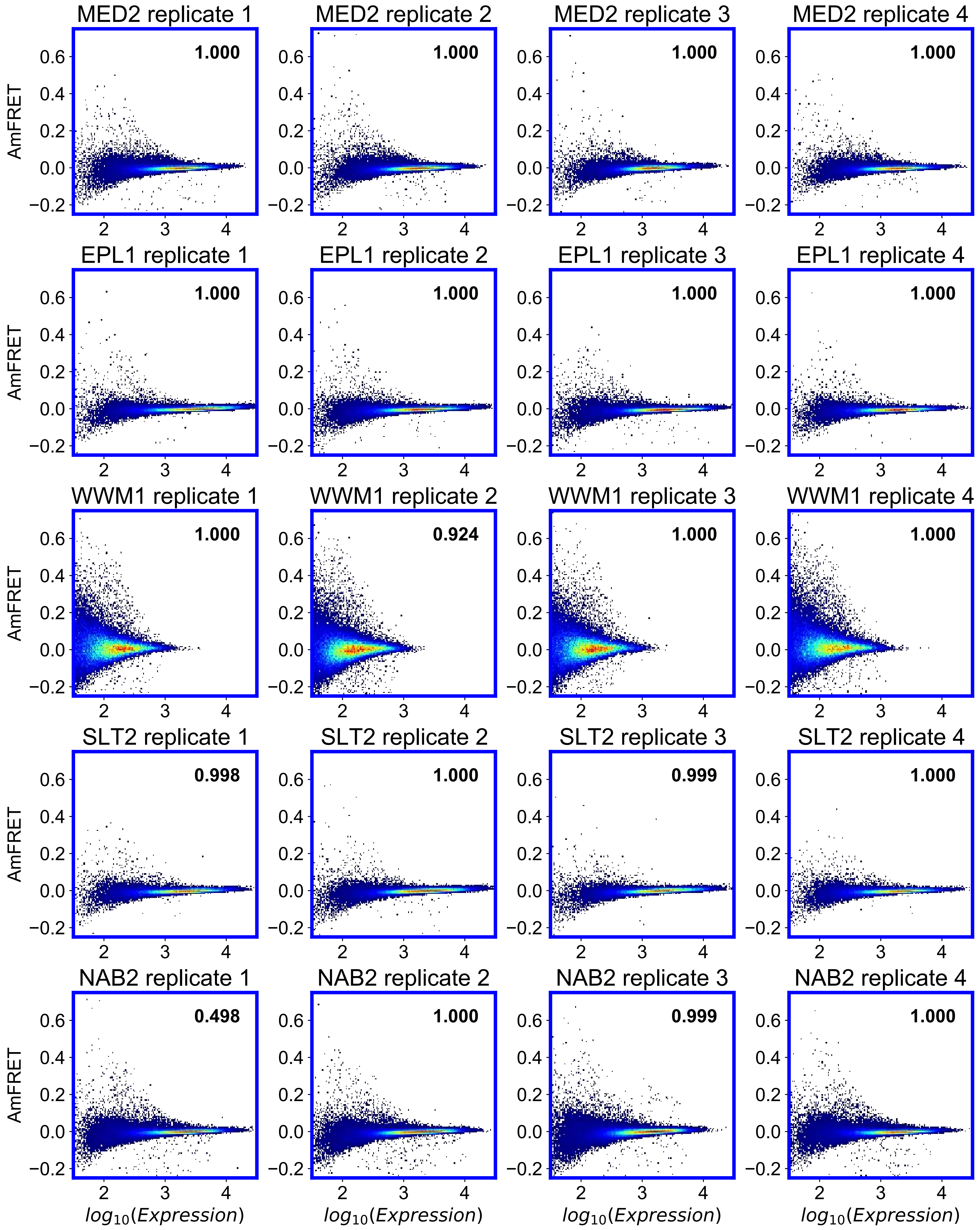

MED2 replicate 1
1.000
MED2 replicate 2
1.000
MED2 replicate 3
1.000
MED2 replicate 4
1.000
EPL1 replicate 1
1.000
EPL1 replicate 2
1.000
EPL1 replicate 3
1.000
EPL1 replicate 4
1.000
WWM1 replicate 1
1.000
WWM1 replicate 2
0.924
WWM1 replicate 3
1.000
WWM1 replicate 4
1.000
SLT2 replicate 1
0.998
SLT2 replicate 2
1.000
SLT2 replicate 3
0.999
SLT2 replicate 4
1.000
NAB2 replicate 1
0.498
NAB2 replicate 2
1.000
NAB2 replicate 3
0.999
NAB2 replicate 4
1.000
AmFRET
$log_{10}(Expression)$

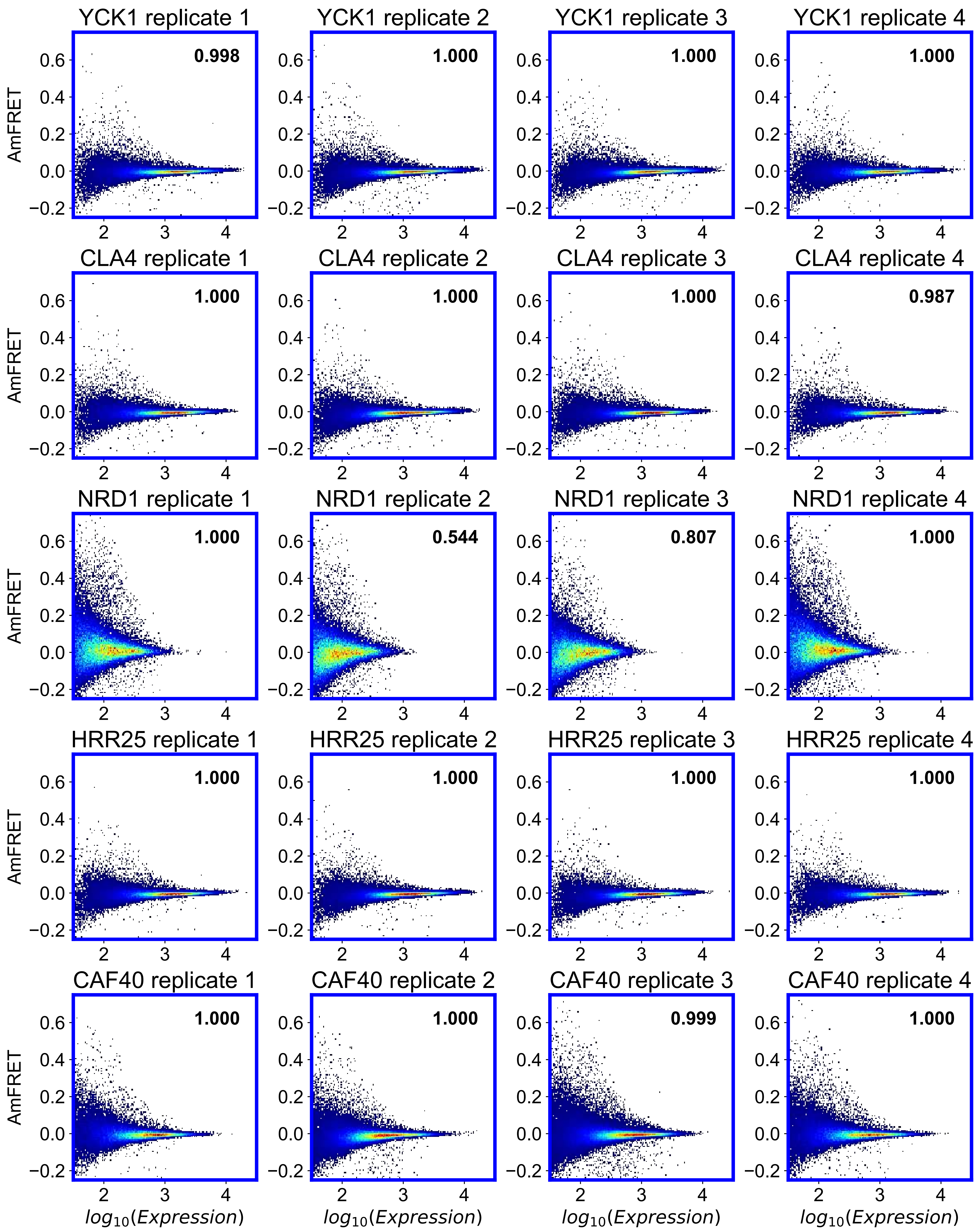

YCK1 replicate 1
0.998
YCK1 replicate 2
1.000
YCK1 replicate 3
1.000
YCK1 replicate 4
1.000
CLA4 replicate 1
1.000
CLA4 replicate 2
1.000
CLA4 replicate 3
1.000
CLA4 replicate 4
0.987
NRD1 replicate 1
1.000
NRD1 replicate 2
0.544
NRD1 replicate 3
0.807
NRD1 replicate 4
1.000
HRR25 replicate 1
1.000
HRR25 replicate 2
1.000
HRR25 replicate 3
1.000
HRR25 replicate 4
1.000
CAF40 replicate 1
1.000
CAF40 replicate 2
1.000
CAF40 replicate 3
0.999
CAF40 replicate 4
1.000
AmFRET
$log_{10}(Expression)$

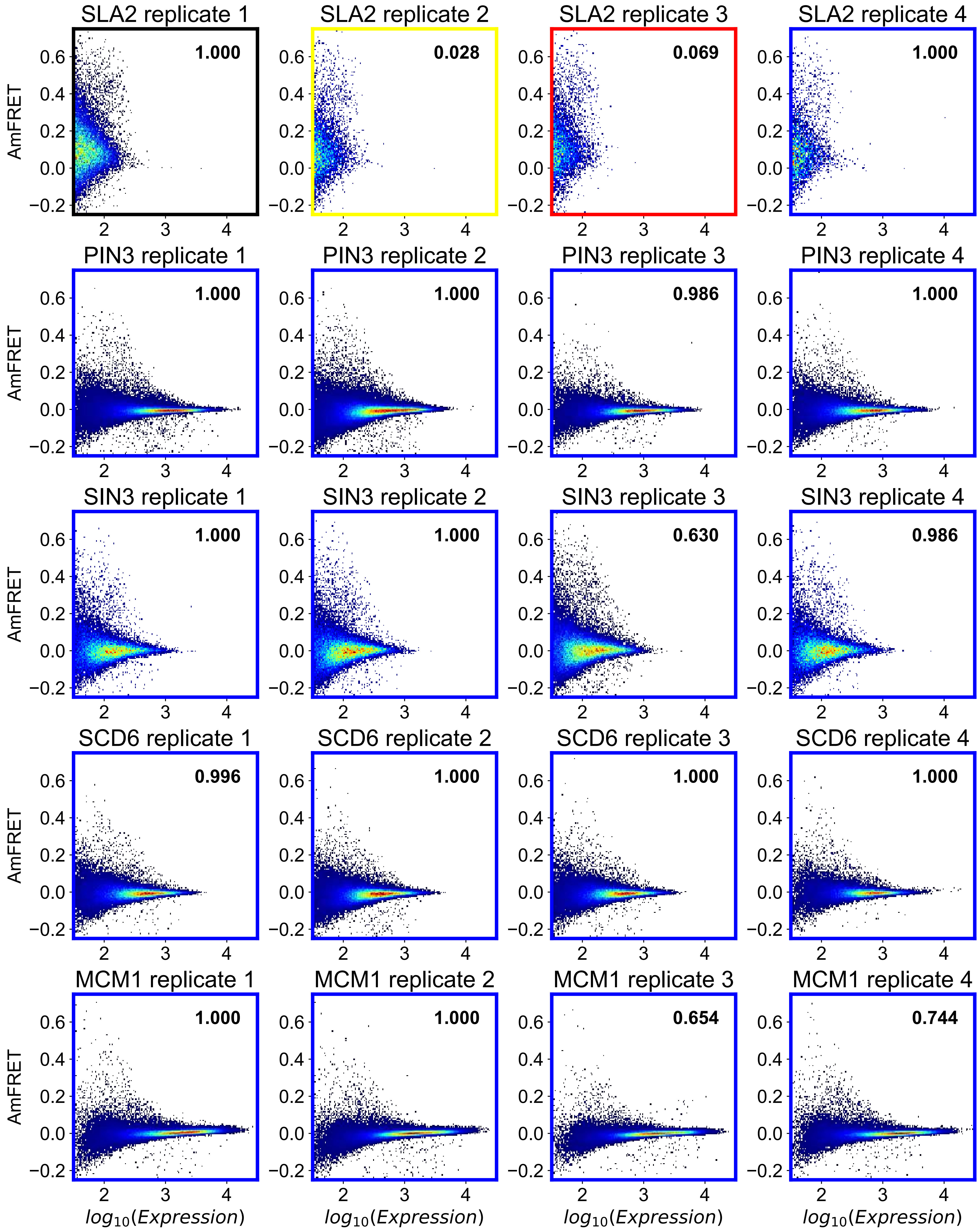
SLA2 replicate 1
1.000
SLA2 replicate 2
0.028
SLA2 replicate 3
0.069
SLA2 replicate 4
1.000
PIN3 replicate 1
1.000
PIN3 replicate 2
1.000
PIN3 replicate 3
0.986
PIN3 replicate 4
1.000
SIN3 replicate 1
1.000
SIN3 replicate 2
1.000
SIN3 replicate 3
0.630
SIN3 replicate 4
0.986
SCD6 replicate 1
0.996
SCD6 replicate 2
1.000
SCD6 replicate 3
1.000
SCD6 replicate 4
1.000
MCM1 replicate 1
1.000
MCM1 replicate 2
1.000
MCM1 replicate 3
0.654
MCM1 replicate 4
0.744
AmFRET
$log_{10}(Expression)$
0.6
0.4
0.2
0.0
−0.2
2
3
4

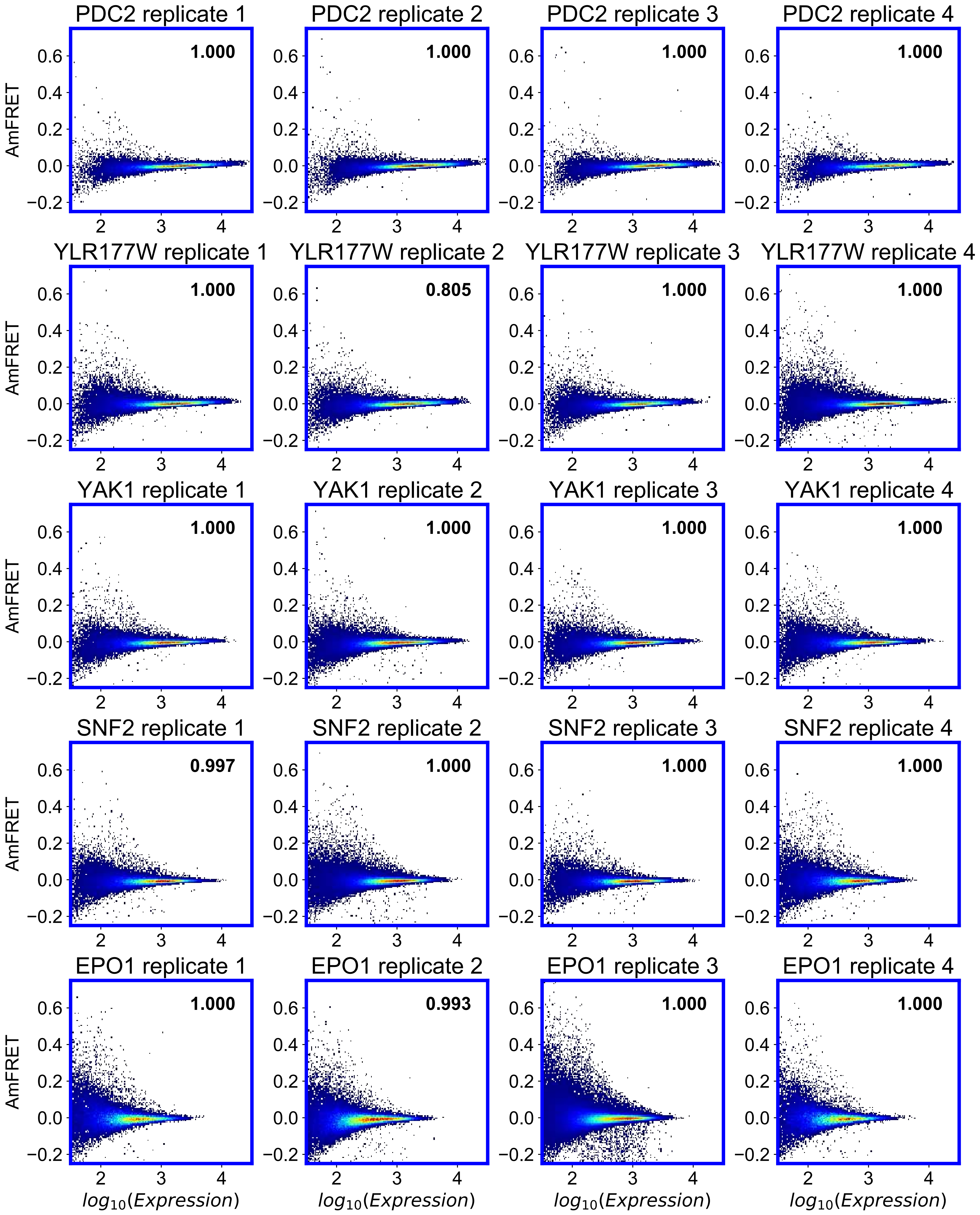
PDC2 replicate 1
1.000
PDC2 replicate 2
1.000
PDC2 replicate 3
1.000
PDC2 replicate 4
1.000
YLR177W replicate 1
1.000
YLR177W replicate 2
0.805
YLR177W replicate 3
1.000
YLR177W replicate 4
1.000
YAK1 replicate 1
1.000
YAK1 replicate 2
1.000
YAK1 replicate 3
1.000
YAK1 replicate 4
1.000
SNF2 replicate 1
0.997
SNF2 replicate 2
1.000
SNF2 replicate 3
1.000
SNF2 replicate 4
1.000
EPO1 replicate 1
1.000
EPO1 replicate 2
0.993
EPO1 replicate 3
1.000
EPO1 replicate 4
1.000
AmFRET
0.6
0.4
0.2
0.0
−0.2
2
3
4
$log_{10}(Expression)$

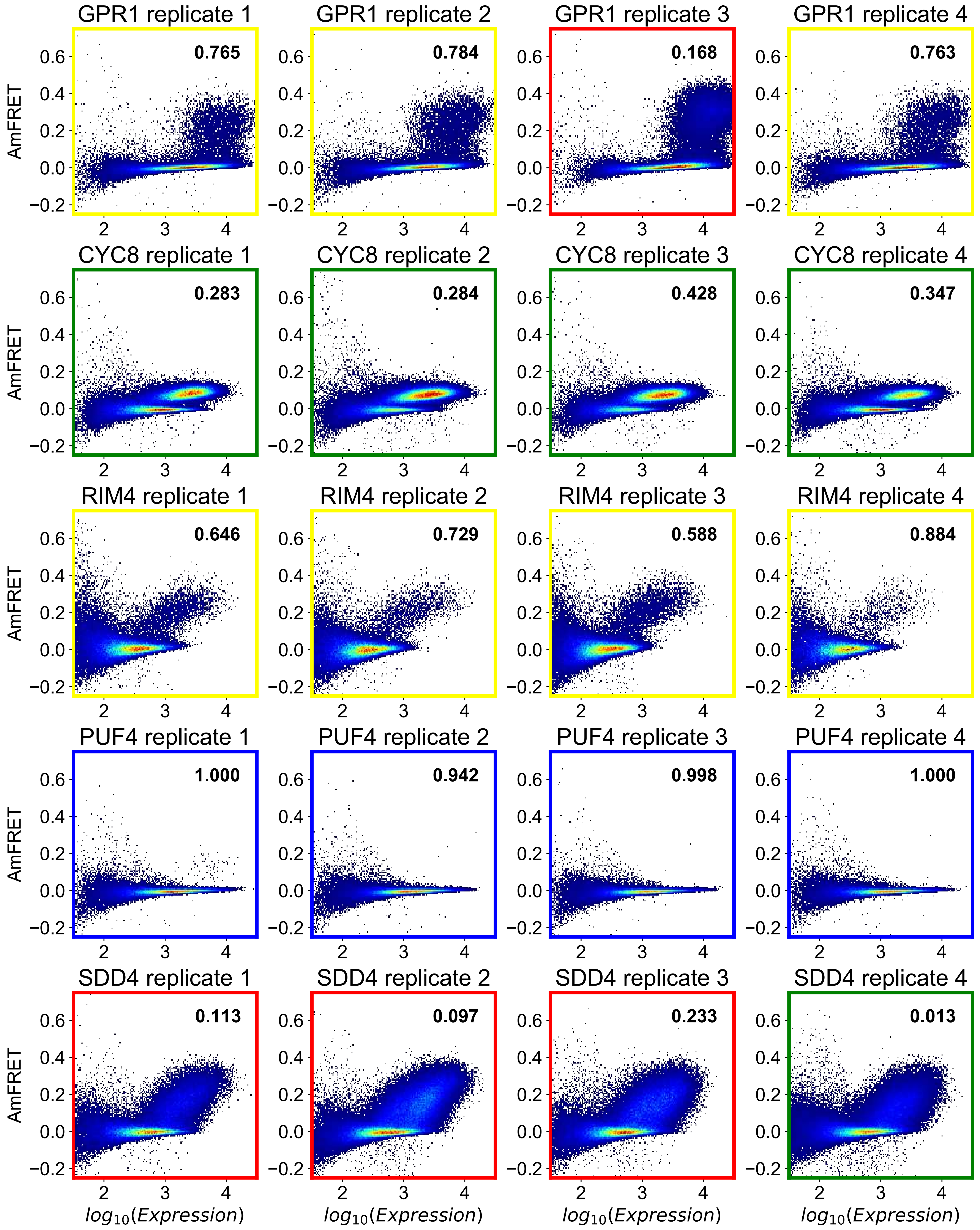

GPR1 replicate 1
0.765
GPR1 replicate 2
0.784
GPR1 replicate 3
0.168
GPR1 replicate 4
0.763
CYC8 replicate 1
0.283
CYC8 replicate 2
0.284
CYC8 replicate 3
0.428
CYC8 replicate 4
0.347
RIM4 replicate 1
0.646
RIM4 replicate 2
0.729
RIM4 replicate 3
0.588
RIM4 replicate 4
0.884
PUF4 replicate 1
1.000
PUF4 replicate 2
0.942
PUF4 replicate 3
0.998
PUF4 replicate 4
1.000
SDD4 replicate 1
0.113
SDD4 replicate 2
0.097
SDD4 replicate 3
0.233
SDD4 replicate 4
0.013
AmFRET
$log_{10}(Expression)$

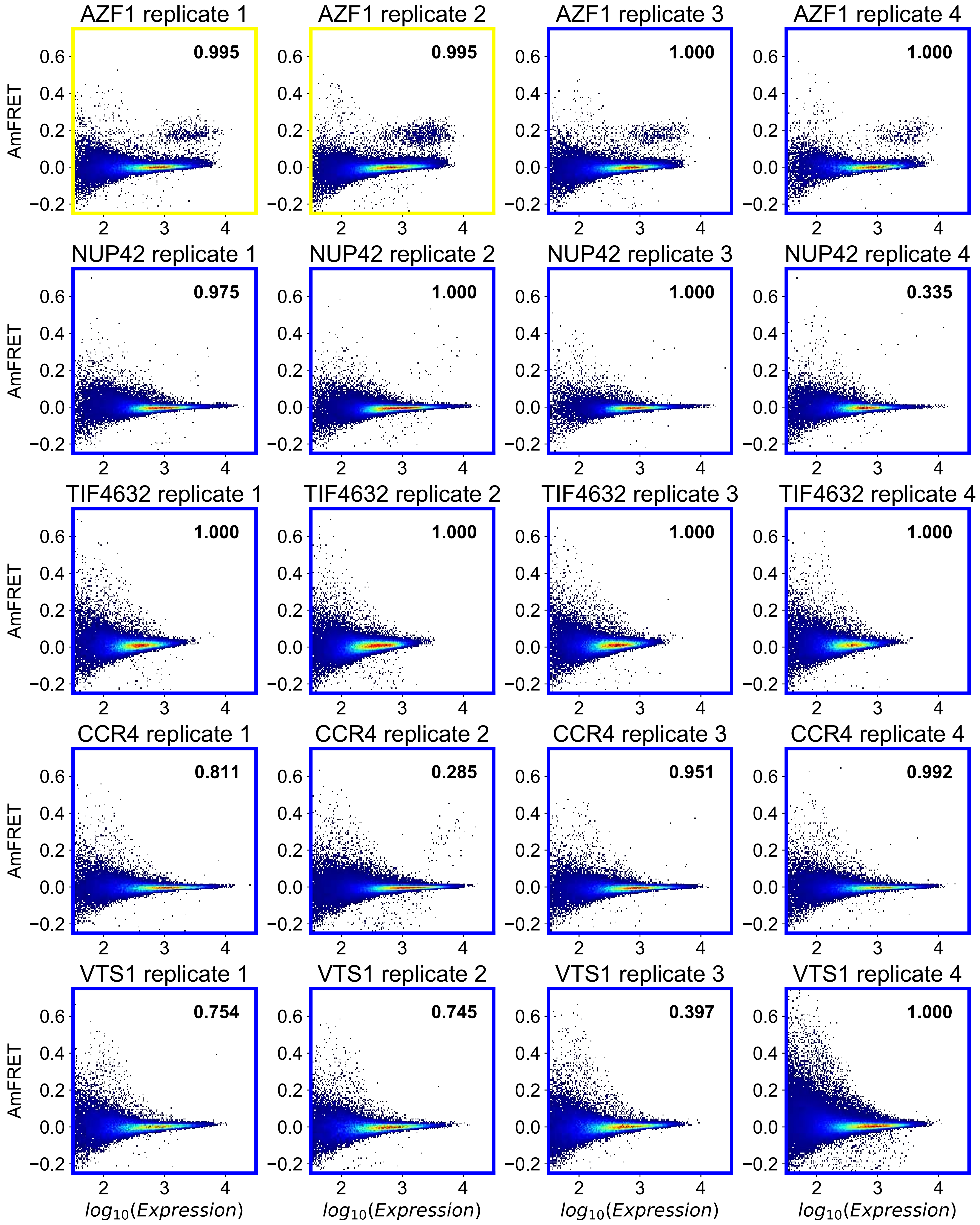
AZF1 replicate 1
0.995
AZF1 replicate 2
0.995
AZF1 replicate 3
1.000
AZF1 replicate 4
1.000
NUP42 replicate 1
0.975
NUP42 replicate 2
1.000
NUP42 replicate 3
1.000
NUP42 replicate 4
0.335
TIF4632 replicate 1
1.000
TIF4632 replicate 2
1.000
TIF4632 replicate 3
1.000
TIF4632 replicate 4
1.000
CCR4 replicate 1
0.811
CCR4 replicate 2
0.285
CCR4 replicate 3
0.951
CCR4 replicate 4
0.992
VTS1 replicate 1
0.754
VTS1 replicate 2
0.745
VTS1 replicate 3
0.397
VTS1 replicate 4
1.000
AmFRET
$log_{10}(Expression)$

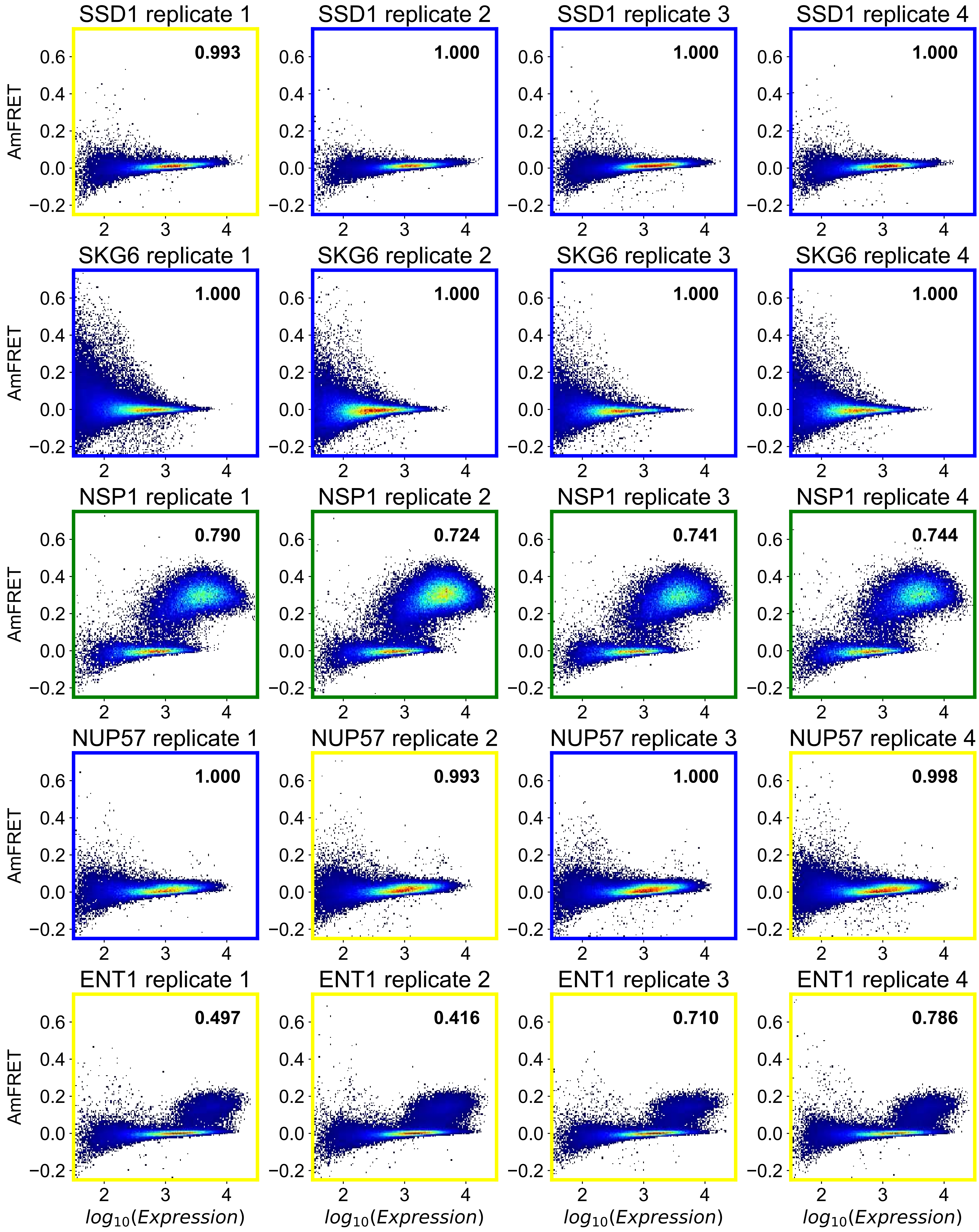
SSD1 replicate 1
0.993
SSD1 replicate 2
1.000
SSD1 replicate 3
1.000
SSD1 replicate 4
1.000
SKG6 replicate 1
1.000
SKG6 replicate 2
1.000
SKG6 replicate 3
1.000
SKG6 replicate 4
1.000
NSP1 replicate 1
0.790
NSP1 replicate 2
0.724
NSP1 replicate 3
0.741
NSP1 replicate 4
0.744
NUP57 replicate 1
1.000
NUP57 replicate 2
0.993
NUP57 replicate 3
1.000
NUP57 replicate 4
0.998
ENT1 replicate 1
0.497
ENT1 replicate 2
0.416
ENT1 replicate 3
0.710
ENT1 replicate 4
0.786
AmFRET
$log_{10}(Expression)$

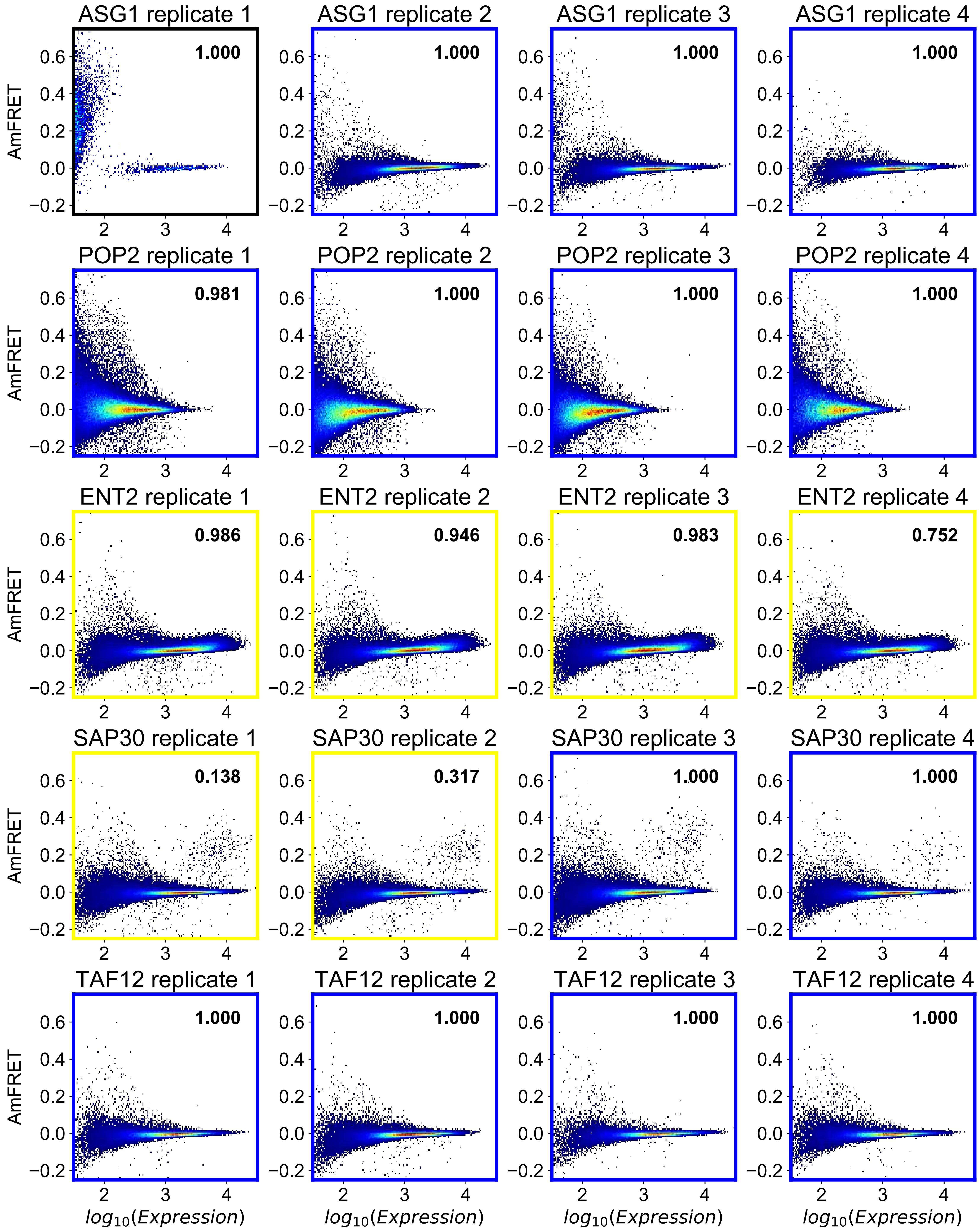

ASG1 replicate 1
1.000
ASG1 replicate 2
1.000
ASG1 replicate 3
1.000
ASG1 replicate 4
1.000
POP2 replicate 1
0.981
POP2 replicate 2
1.000
POP2 replicate 3
1.000
POP2 replicate 4
1.000
ENT2 replicate 1
0.986
ENT2 replicate 2
0.946
ENT2 replicate 3
0.983
ENT2 replicate 4
0.752
SAP30 replicate 1
0.138
SAP30 replicate 2
0.317
SAP30 replicate 3
1.000
SAP30 replicate 4
1.000
TAF12 replicate 1
1.000
TAF12 replicate 2
1.000
TAF12 replicate 3
1.000
TAF12 replicate 4
1.000
AmFRET
0.6
0.4
0.2
0.0
−0.2
2
3
4
$log_{10}(Expression)$

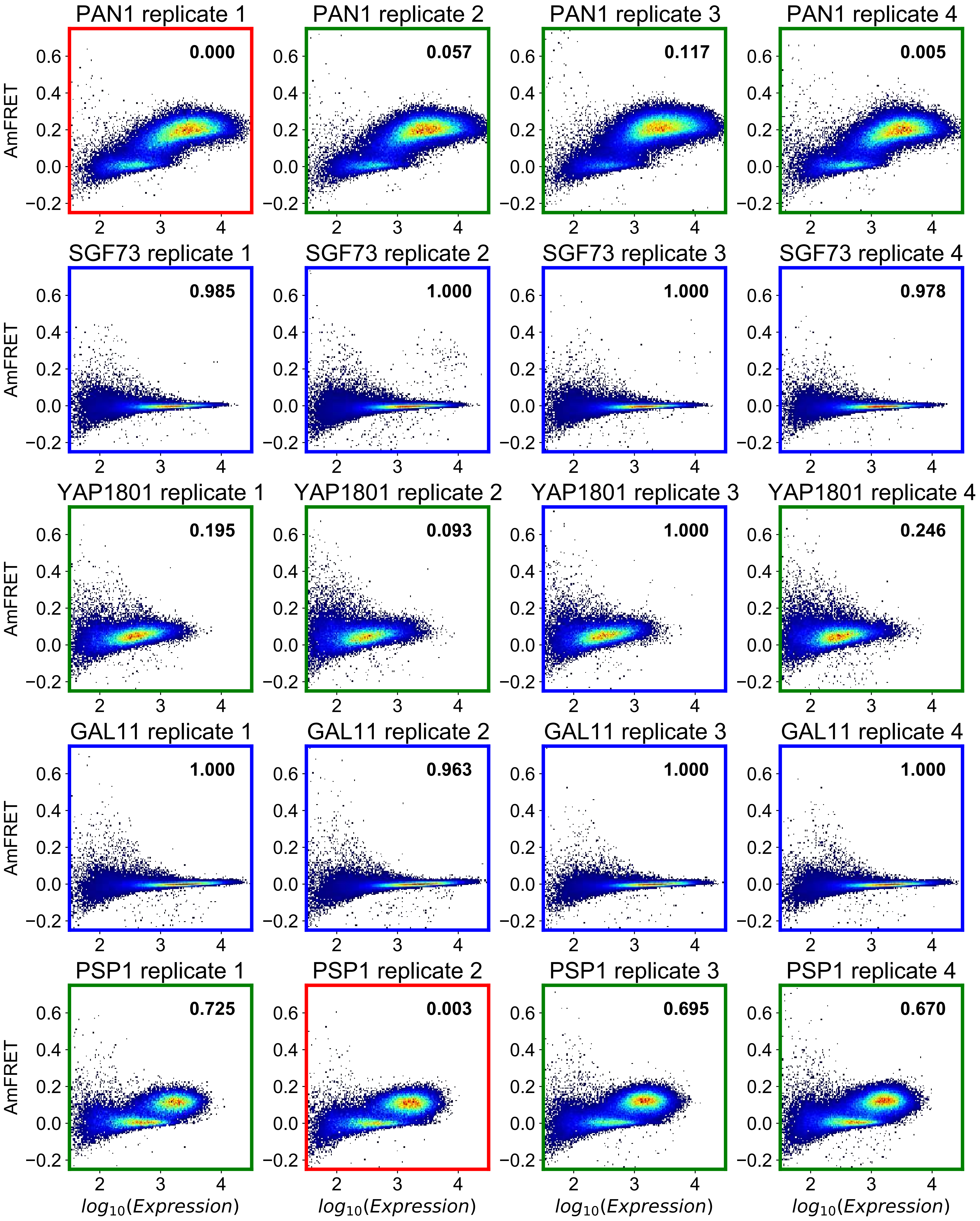
PAN1 replicate 1
0.000
PAN1 replicate 2
0.057
PAN1 replicate 3
0.117
PAN1 replicate 4
0.005
SGF73 replicate 1
0.985
SGF73 replicate 2
1.000
SGF73 replicate 3
1.000
SGF73 replicate 4
0.978
YAP1801 replicate 1
0.195
YAP1801 replicate 2
0.093
YAP1801 replicate 3
1.000
YAP1801 replicate 4
0.246
GAL11 replicate 1
1.000
GAL11 replicate 2
0.963
GAL11 replicate 3
1.000
GAL11 replicate 4
1.000
PSP1 replicate 1
0.725
PSP1 replicate 2
0.003
PSP1 replicate 3
0.695
PSP1 replicate 4
0.670
AmFRET
$log_{10}(Expression)$

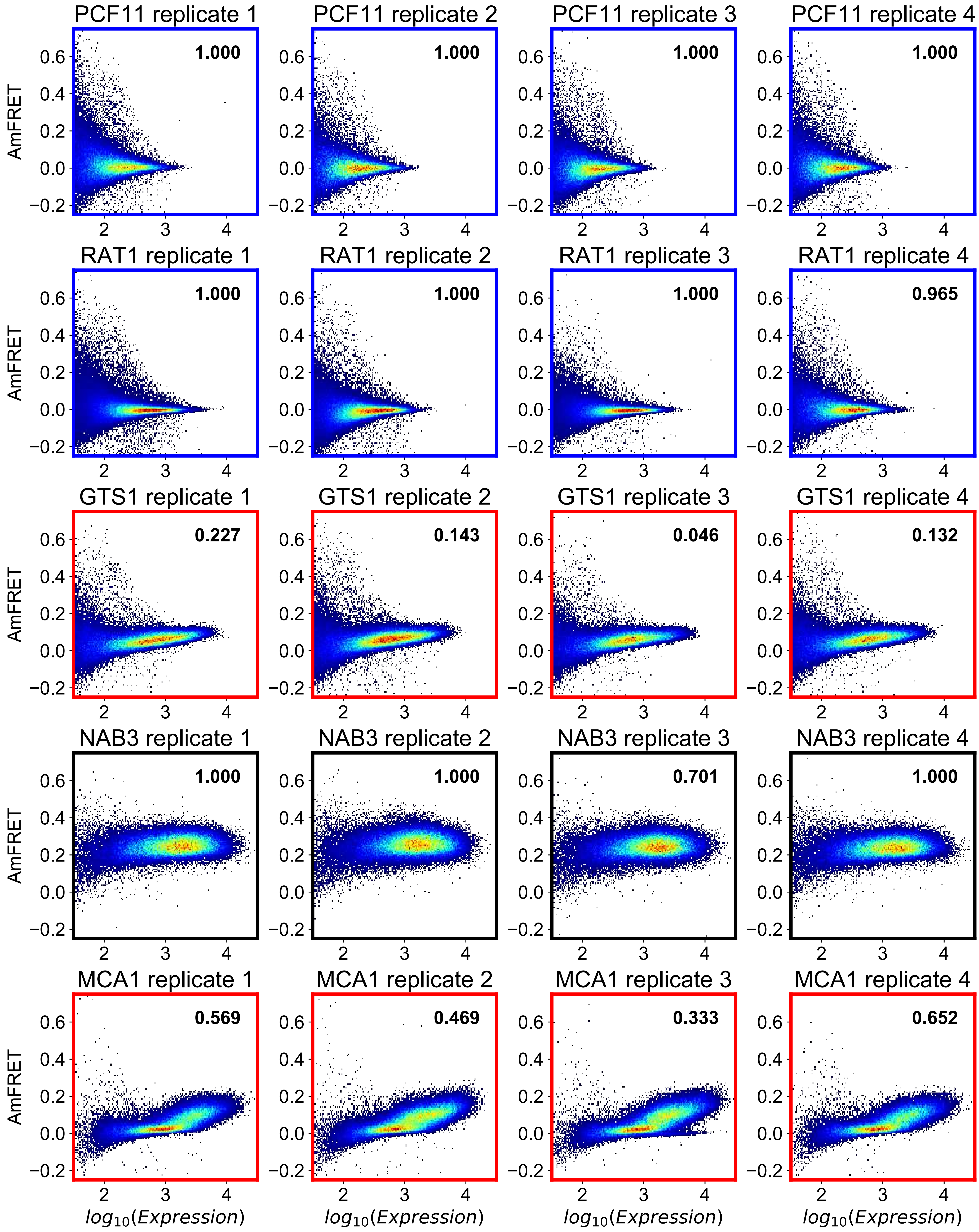
PCF11 replicate 1
1.000
PCF11 replicate 2
1.000
PCF11 replicate 3
1.000
PCF11 replicate 4
1.000
RAT1 replicate 1
1.000
RAT1 replicate 2
1.000
RAT1 replicate 3
1.000
RAT1 replicate 4
0.965
GTS1 replicate 1
0.227
GTS1 replicate 2
0.143
GTS1 replicate 3
0.046
GTS1 replicate 4
0.132
NAB3 replicate 1
1.000
NAB3 replicate 2
1.000
NAB3 replicate 3
0.701
NAB3 replicate 4
1.000
MCA1 replicate 1
0.569
MCA1 replicate 2
0.469
MCA1 replicate 3
0.333
MCA1 replicate 4
0.652
AmFRET
$log_{10}(Expression)$

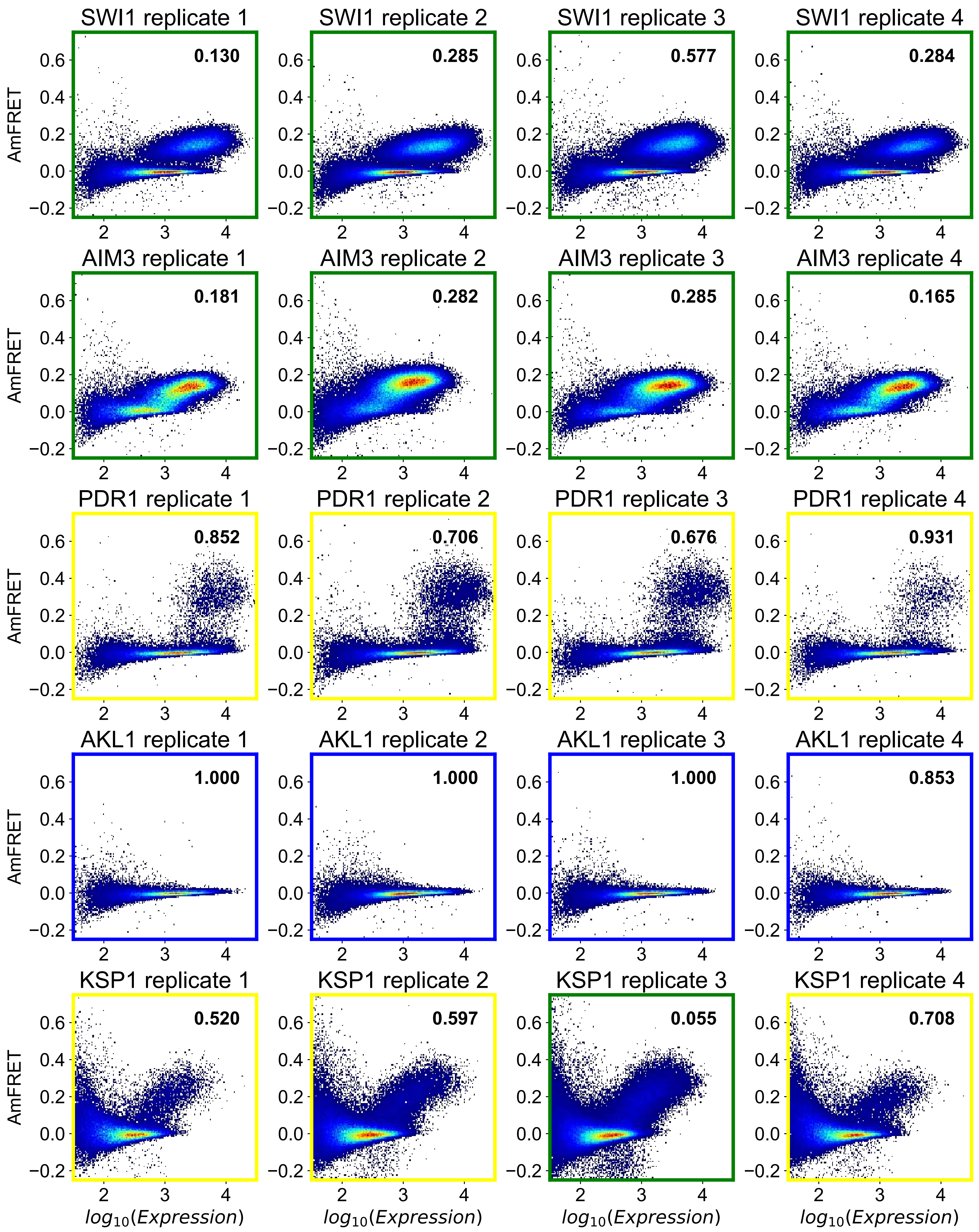
SWI1 replicate 1
0.130
SWI1 replicate 2
0.285
SWI1 replicate 3
0.577
SWI1 replicate 4
0.284
AIM3 replicate 1
0.181
AIM3 replicate 2
0.282
AIM3 replicate 3
0.285
AIM3 replicate 4
0.165
PDR1 replicate 1
0.852
PDR1 replicate 2
0.706
PDR1 replicate 3
0.676
PDR1 replicate 4
0.931
AKL1 replicate 1
1.000
AKL1 replicate 2
1.000
AKL1 replicate 3
1.000
AKL1 replicate 4
0.853
KSP1 replicate 1
0.520
KSP1 replicate 2
0.597
KSP1 replicate 3
0.055
KSP1 replicate 4
0.708
AmFRET
$log_{10}(Expression)$

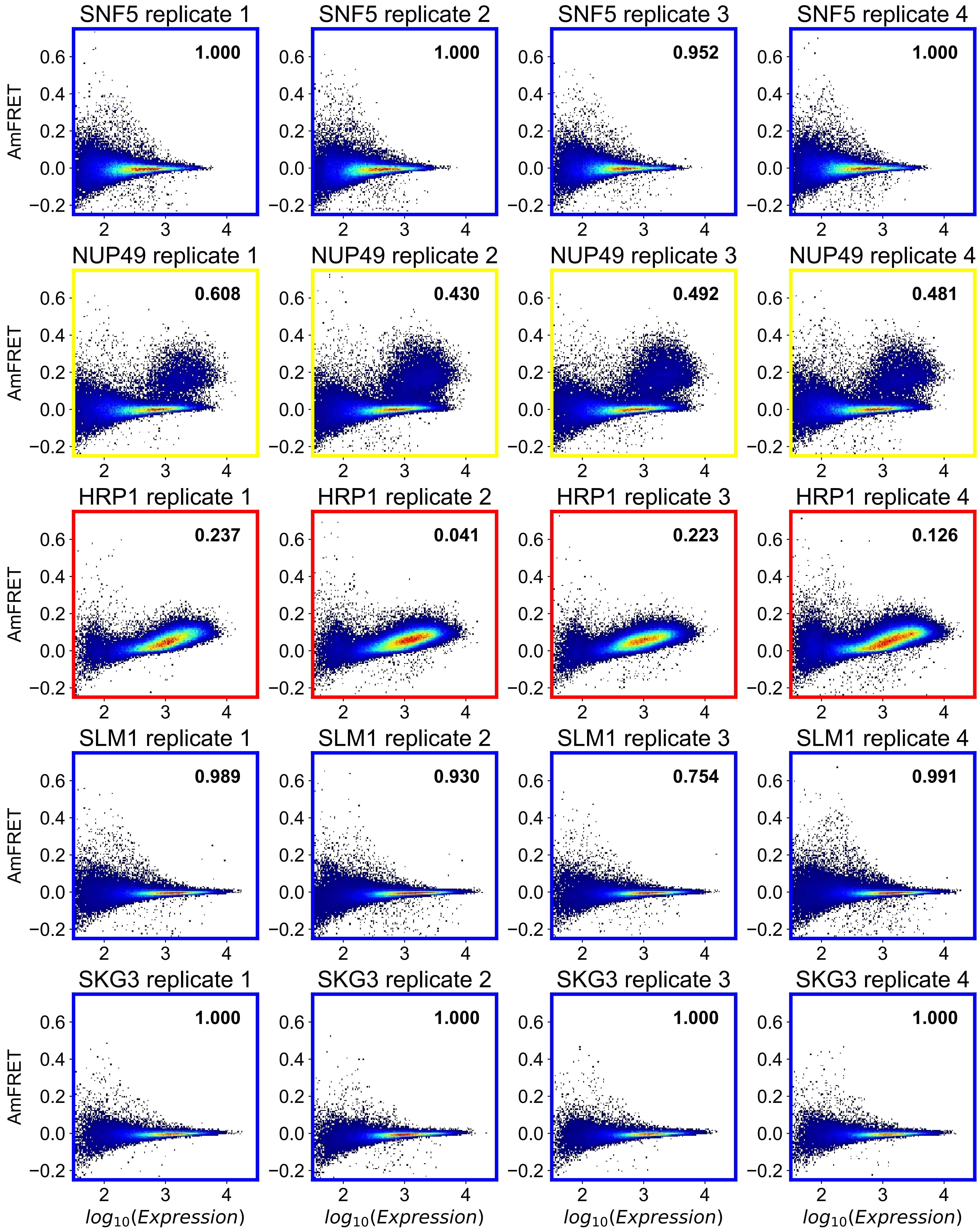

SNF5 replicate 1
1.000
SNF5 replicate 2
1.000
SNF5 replicate 3
0.952
SNF5 replicate 4
1.000
NUP49 replicate 1
0.608
NUP49 replicate 2
0.430
NUP49 replicate 3
0.492
NUP49 replicate 4
0.481
HRP1 replicate 1
0.237
HRP1 replicate 2
0.041
HRP1 replicate 3
0.223
HRP1 replicate 4
0.126
SLM1 replicate 1
0.989
SLM1 replicate 2
0.930
SLM1 replicate 3
0.754
SLM1 replicate 4
0.991
SKG3 replicate 1
1.000
SKG3 replicate 2
1.000
SKG3 replicate 3
1.000
SKG3 replicate 4
1.000
AmFRET
$log_{10}(Expression)$

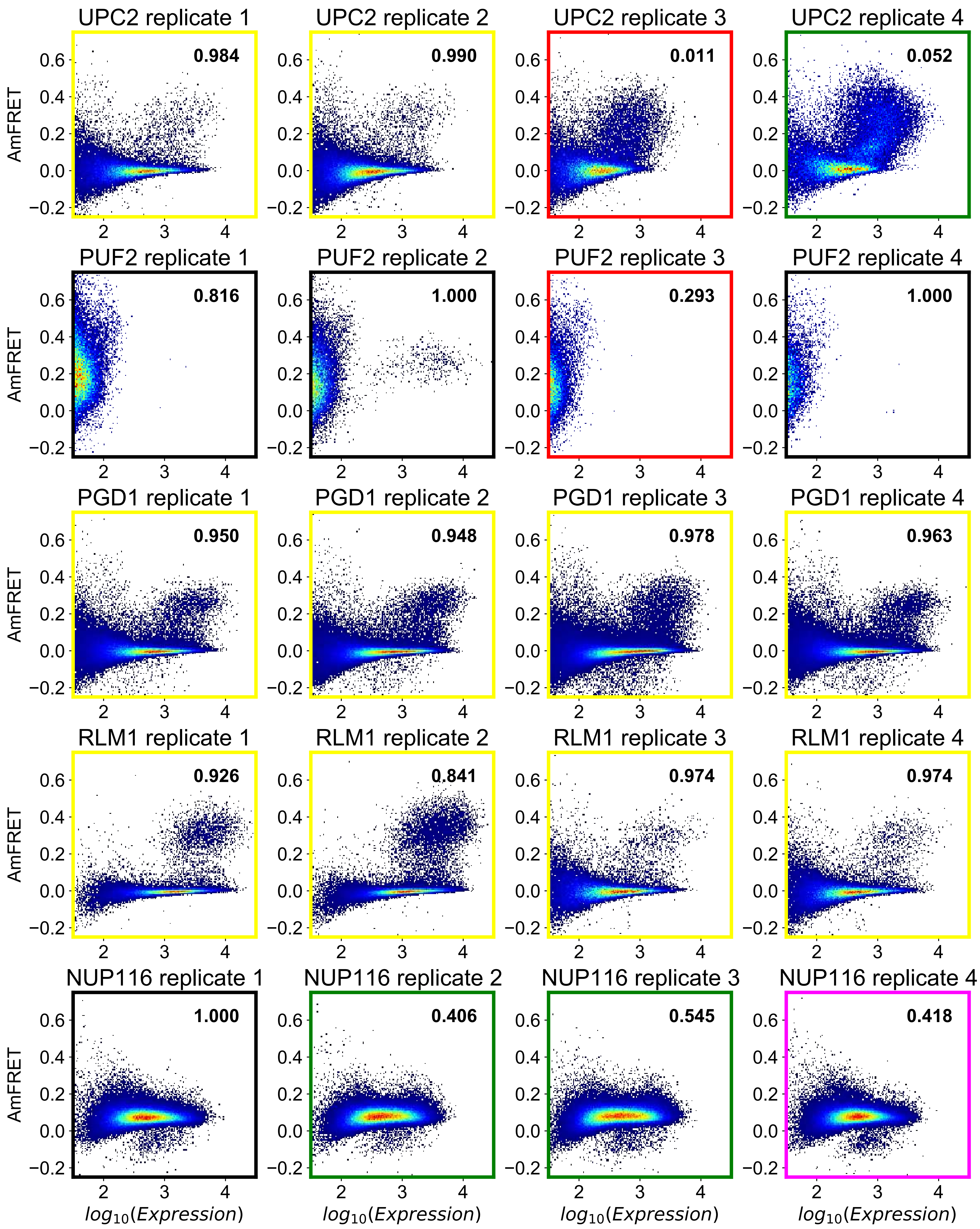

UPC2 replicate 1
0.984
UPC2 replicate 2
0.990
UPC2 replicate 3
0.011
UPC2 replicate 4
0.052
PUF2 replicate 1
0.816
PUF2 replicate 2
1.000
PUF2 replicate 3
0.293
PUF2 replicate 4
1.000
PGD1 replicate 1
0.950
PGD1 replicate 2
0.948
PGD1 replicate 3
0.978
PGD1 replicate 4
0.963
RLM1 replicate 1
0.926
RLM1 replicate 2
0.841
RLM1 replicate 3
0.974
RLM1 replicate 4
0.974
NUP116 replicate 1
1.000
NUP116 replicate 2
0.406
NUP116 replicate 3
0.545
NUP116 replicate 4
0.418
AmFRET
0.6
0.4
0.2
0.0
−0.2
2
3
4
$log_{10}(Expression)$

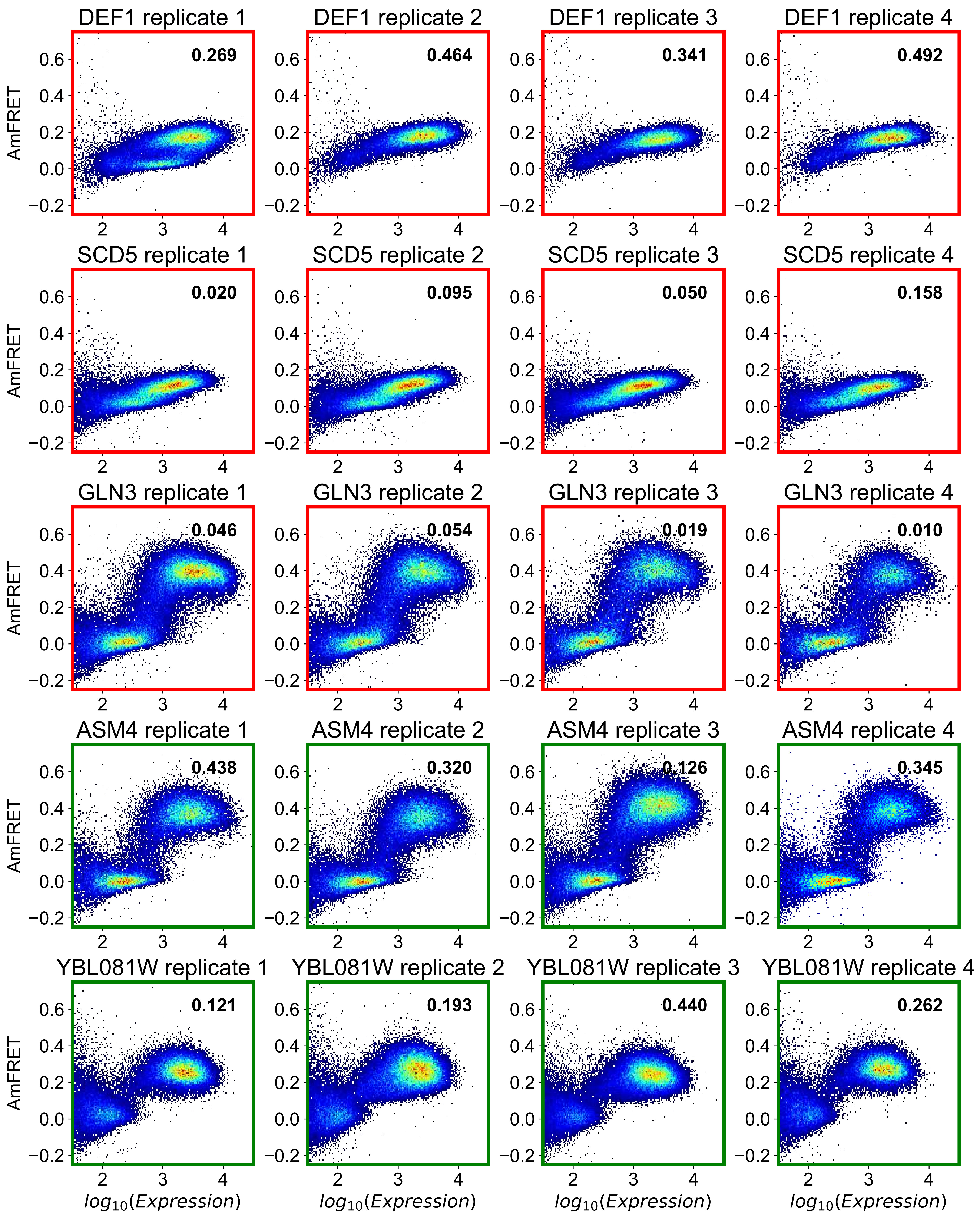
DEF1 replicate 1
0.269
DEF1 replicate 2
0.464
DEF1 replicate 3
0.341
DEF1 replicate 4
0.492
SCD5 replicate 1
0.020
SCD5 replicate 2
0.095
SCD5 replicate 3
0.050
SCD5 replicate 4
0.158
GLN3 replicate 1
0.046
GLN3 replicate 2
0.054
GLN3 replicate 3
0.019
GLN3 replicate 4
0.010
ASM4 replicate 1
0.438
ASM4 replicate 2
0.320
ASM4 replicate 3
0.126
ASM4 replicate 4
0.345
YBL081W replicate 1
0.121
YBL081W replicate 2
0.193
YBL081W replicate 3
0.440
YBL081W replicate 4
0.262
AmFRET
$log_{10}(Expression)$

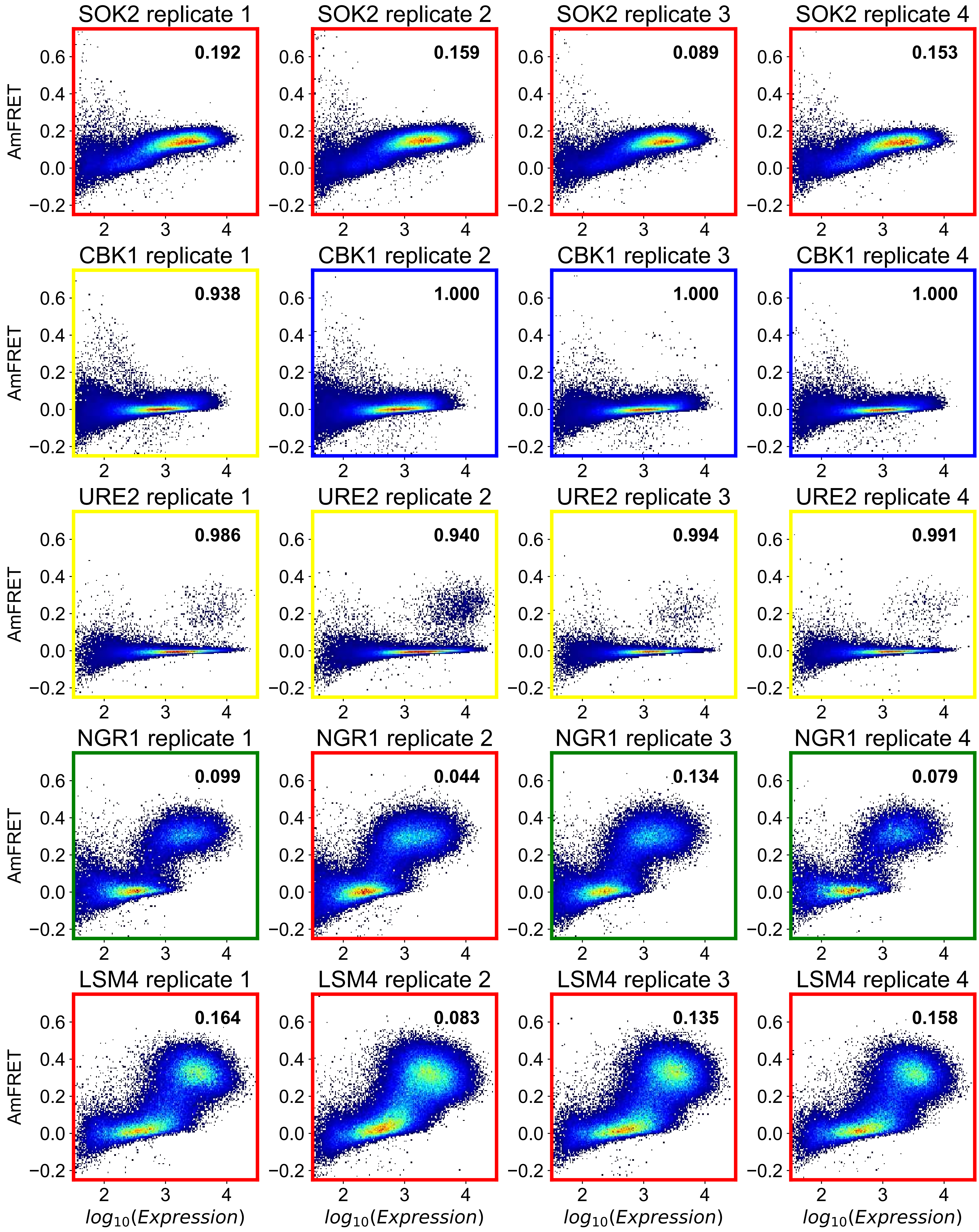

SOK2 replicate 1
0.192
SOK2 replicate 2
0.159
SOK2 replicate 3
0.089
SOK2 replicate 4
0.153
CBK1 replicate 1
0.938
CBK1 replicate 2
1.000
CBK1 replicate 3
1.000
CBK1 replicate 4
1.000
URE2 replicate 1
0.986
URE2 replicate 2
0.940
URE2 replicate 3
0.994
URE2 replicate 4
0.991
NGR1 replicate 1
0.099
NGR1 replicate 2
0.044
NGR1 replicate 3
0.134
NGR1 replicate 4
0.079
LSM4 replicate 1
0.164
LSM4 replicate 2
0.083
LSM4 replicate 3
0.135
LSM4 replicate 4
0.158
AmFRET
$log_{10}(Expression)$

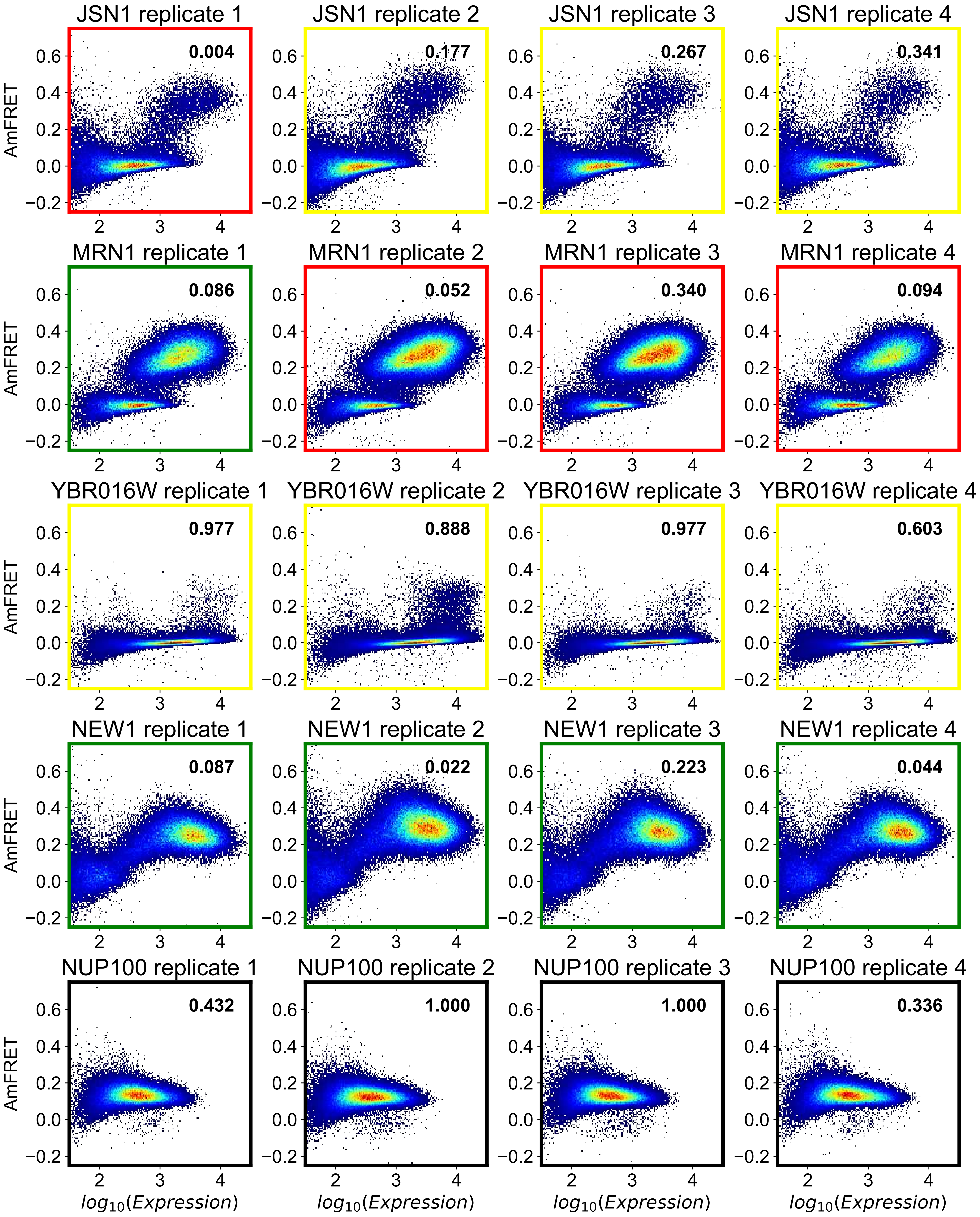

JSN1 replicate 1
0.004
JSN1 replicate 2
0.177
JSN1 replicate 3
0.267
JSN1 replicate 4
0.341
MRN1 replicate 1
0.086
MRN1 replicate 2
0.052
MRN1 replicate 3
0.340
MRN1 replicate 4
0.094
YBR016W replicate 1
0.977
YBR016W replicate 2
0.888
YBR016W replicate 3
0.977
YBR016W replicate 4
0.603
NEW1 replicate 1
0.087
NEW1 replicate 2
0.022
NEW1 replicate 3
0.223
NEW1 replicate 4
0.044
NUP100 replicate 1
0.432
NUP100 replicate 2
1.000
NUP100 replicate 3
1.000
NUP100 replicate 4
0.336
AmFRET
$log_{10}(Expression)$

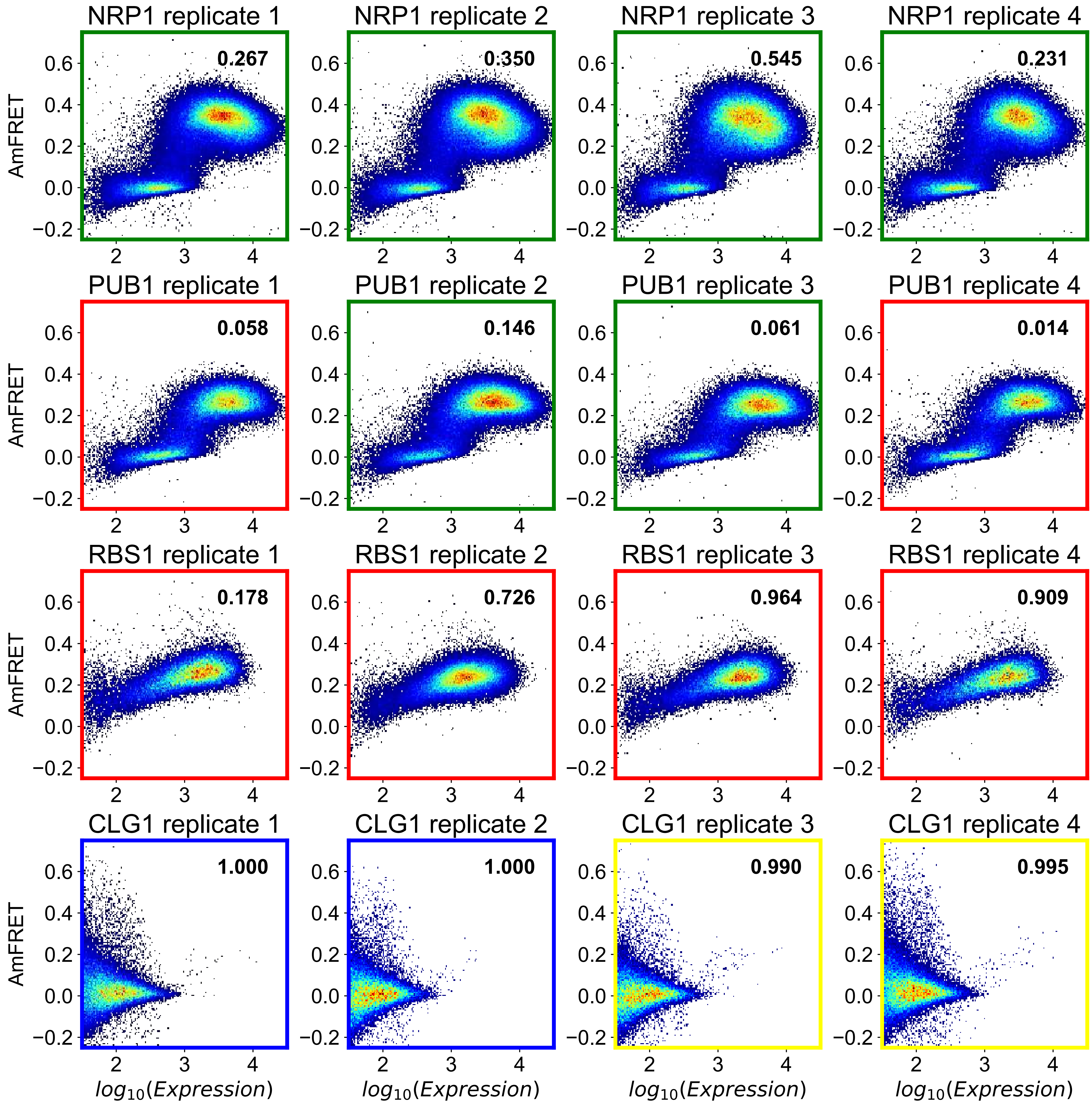

NRP1 replicate 1
0.267
NRP1 replicate 2
0.350
NRP1 replicate 3
0.545
NRP1 replicate 4
0.231
PUB1 replicate 1
0.058
PUB1 replicate 2
0.146
PUB1 replicate 3
0.061
PUB1 replicate 4
0.014
RBS1 replicate 1
0.178
RBS1 replicate 2
0.726
RBS1 replicate 3
0.964
RBS1 replicate 4
0.909
CLG1 replicate 1
1.000
CLG1 replicate 2
1.000
CLG1 replicate 3
0.990
CLG1 replicate 4
0.995
AmFRET
log10(Expression)

**Figure S5: Classification of DAmFRET histograms of 84 cPrDs previously examined by Alberti et al. without shading by confidence score**. cPrDs are sorted by their PAPA scores shown in parentheses. Each column denotes a different experimental replicate.

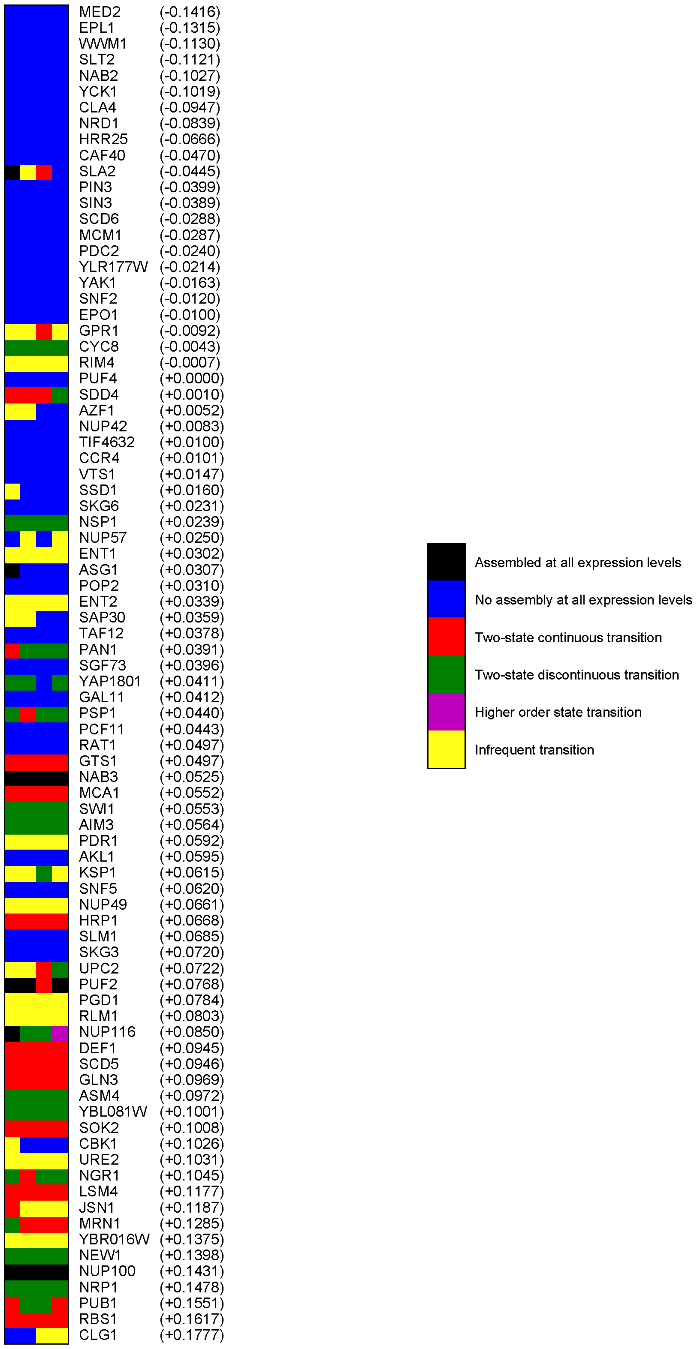

**Figure S6: Correlation between amino acid frequency and degree of discontinuity for all 20 amino acids and all 23 cPrDs classified as undergoing a two-state transition.** Each point corresponds to an experimental replicate. The color of each point denotes the degree of discontinuity of the transition. Numbers indicate the Pearson r-values used to quantify positive or negative correlations.

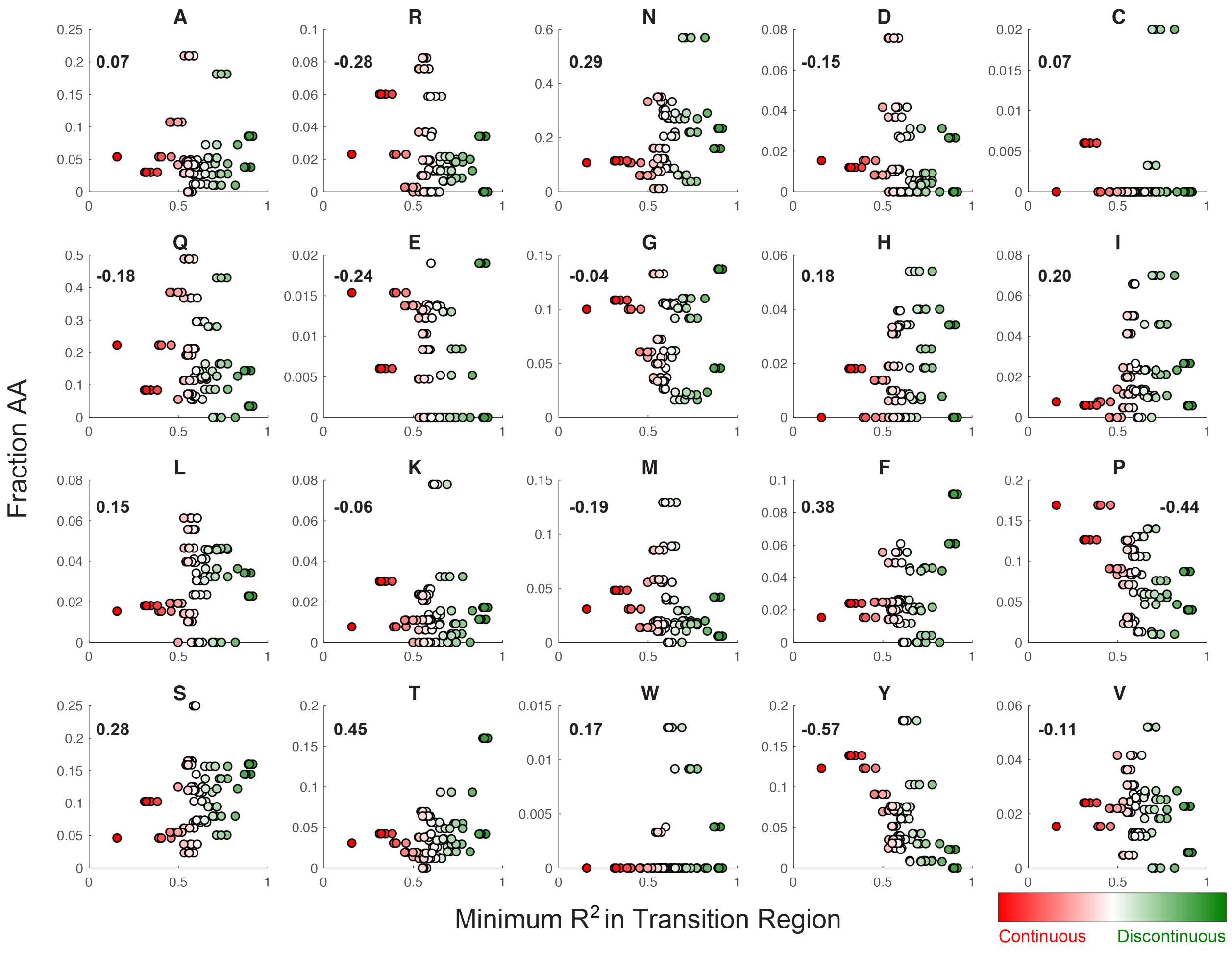

**Figure S7: Distribution of the number of individual cell measurements collected for the set of 94 cPrDs from the Alberti et al. dataset**. Bin widths were set to 5,000.

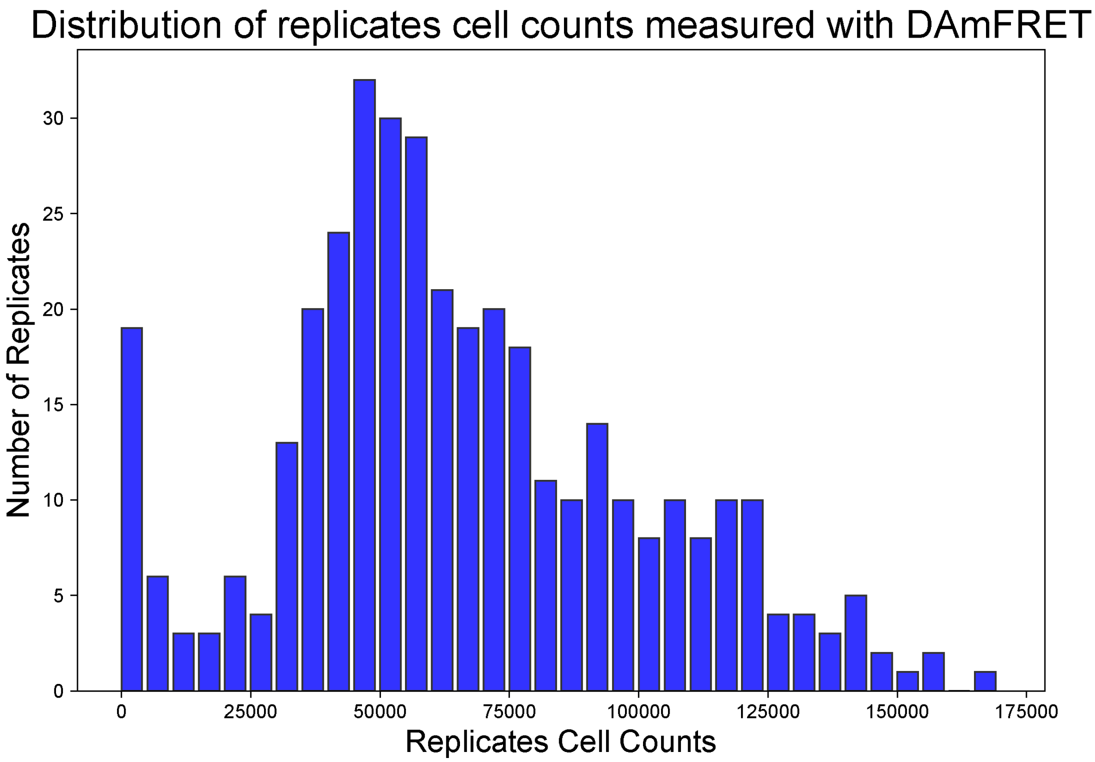

**Figure S8: The Shannon Entropy was used to identify a suitable range of acceptable grid sizes that could be applied across an entire collection of DAmFRET data**. To determine an acceptable grid size which could be applied across a majority of replicates for subsequent analysis, we examined the information density quantified by the Shannon Entropy, $S = -\sum_{i=1}^{n_x}\sum_{j=1}^{n_y} p_{ij}\log p_{ij}$, and its change as a function of increasing grid size, on a subset of replicates. We identified 6 replicates with the following cell measurements: $1\times10^4$, $2\times10^4$, $5\times10^4$, $8\times10^4$, $10\times10^4$, and $12\times10^4$ as these adequately sample the range in the number of cell measurements within the extrema. Each grid is generated from a sum-normalized 2D histogram of each replicate along the x-axis (Expression) and y-axis (AmFRET / Expression i.e. $\Omega_{FRET}$), which is binned into the same number of $n_x$ and $n_y$ bins. Asymmetric grid sizes were not chosen.

In (A), as the grid size increases, the Shannon Entropy increases in a logarithmic fashion until it reaches a maximum that is dependent on the dataset size. As this change is non-linear, the turning points along each curve define lower and upper limits for possible grid sizes for replicates with a similar number of cell measurements. The location of the second turning point along the x-axis in which the entropy approaches its maximum defines an upper range in grid size as beyond that limit the information gained is minimized. Similarly, examining the location of the first turning point along each Shannon Entropy curve provides insight into the location of a lower limit of the grid size which can be applied on a given dataset.

These insights, while informative, do not substantially narrow the range of possible grid sizes. Instead, by examining the numerical derivative of the Shannon Entropy curves as a function of grid size, we can better identify the grid sizes at which the information density change is maximized. However, as the entropy change can be positive or negative, we instead examine the absolute value of the numerical derivative of the Shannon Entropy curves, $|\Delta S|$.

In (B), the peak of each absolute derivative curve $|\Delta S|$, identified by a grey line, indicates the grid size at which the information density gained or lost is maximized or minimized. Contextually, larger values in $|\Delta S|$ correspond to significant changes in sparsity of the grid at a given size. This suggests that beyond the peak, the information density gained or lost by increasing the sparsity of a grid is smaller overall. Hence, grid sizes chosen at or near to this value would be ideal.

For each replicate, the grid size corresponding to the maximum $|\Delta S|$ is different but not necessarily unique. In general, the replicates with lower cell measurements ≤ $8\times10^4$ peak at much smaller grid sizes, 46. The replicates with more cell measurements peak at grid sizes of 215 and 464. Thus, these values define an acceptable range of possible grid values which could be applied across all replicates.

However, any grid size chosen is mediated by being able to distinguish features within the grids. Grids generated at the grid size corresponding to the smallest grid size peak (46) produce featureless representations, while grids generated at the larger grid sizes (215 and 464) are more featurefull and sparse. As we desire to better distinguish inherent features within the landscapes, values closer to the peak at the largest grid size is more ideal.

Thus, we chose a grid size of $300\times300$ as it can be applied to replicates with at least $5\times10^4$ cell measurements and is more featureful. Most of the replicates contain cell measurements above that number. Given that replicates at lower counts tend to experience significant $|\Delta S|$ loss at the $300 \times 300$ grid size, any subsequent analysis may be impaired. Hence, we excluded cPrDs with replicates < $2 \times 10^4$ cell measurements from our analysis. The cPrDs that fell below this threshold are: PEX13, TIF4631, NPL3, RGT1, PSP2, YAP1802, SLA1, MOT3, DDR48, and SFP1.

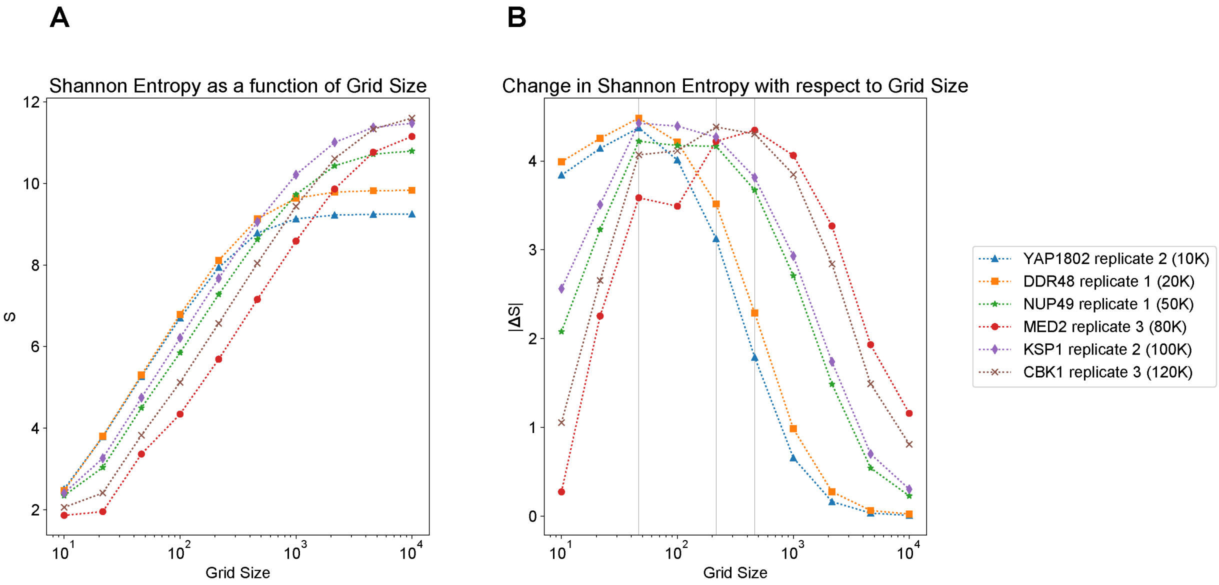

**Table S1: Synthetic Data Classifications.** Noise 1 implies low noise, Noise 2 implies medium noise, and Noise 3 implies high noise. Saturation denotes how points were added to the dataset and Confidence-Score denotes how confident we are in the classification. These variables are described in the Materials and Methods.

| log10(c50) | Generating-Function | Replicate | Saturation | Noise-Profile | Classification | Confidence-Score |
|---|---|---|---|---|---|---|
| 1 | 3 State | 1 | High | Noise 1 | Higher order state transition (magenta) | 0.621546 |
| 1 | 3 State | 2 | High | Noise 1 | Higher order state transition (magenta) | 0.302117 |
| 1 | 3 State | 3 | High | Noise 1 | Higher order state transition (magenta) | 0.384786 |
| 1 | 3 State | 1 | High | Noise 2 | Higher order state transition (magenta) | 0.274063 |
| 1 | 3 State | 2 | High | Noise 2 | Higher order state transition (magenta) | 0.108933 |
| 1 | 3 State | 3 | High | Noise 2 | Higher order state transition (magenta) | 0.236509 |
| 1 | 3 State | 1 | High | Noise 3 | Higher order state transition (magenta) | 0.157181 |
| 1 | 3 State | 2 | High | Noise 3 | Higher order state transition (magenta) | 0.121959 |
| 1 | 3 State | 3 | High | Noise 3 | Higher order state transition (magenta) | 0.250389 |
| 2 | 3 State | 1 | High | Noise 1 | Higher order state transition (magenta) | 0.536798 |
| 2 | 3 State | 2 | High | Noise 1 | Higher order state transition (magenta) | 0.49149 |
| 2 | 3 State | 3 | High | Noise 1 | Higher order state transition (magenta) | 0.655625 |
| 2 | 3 State | 1 | High | Noise 2 | Higher order state transition (magenta) | 0.623963 |
| 2 | 3 State | 2 | High | Noise 2 | Higher order state transition (magenta) | 0.5088 |
| 2 | 3 State | 3 | High | Noise 2 | Higher order state transition (magenta) | 0.427051 |
| 2 | 3 State | 1 | High | Noise 3 | Higher order state transition (magenta) | 0.215512 |
| 2 | 3 State | 2 | High | Noise 3 | Higher order state transition (magenta) | 0.153058 |
| 2 | 3 State | 3 | High | Noise 3 | Higher order state transition (magenta) | 0.0699569 |
| 4 | 3 State | 1 | High | Noise 1 | Higher order state transition (magenta) | 0.622594 |
| 4 | 3 State | 2 | High | Noise 1 | Higher order state transition (magenta) | 0.702502 |
| 4 | 3 State | 3 | High | Noise 1 | Higher order state transition (magenta) | 0.702645 |
| 4 | 3 State | 1 | High | Noise 2 | Higher order state transition (magenta) | 0.617313 |
| 4 | 3 State | 2 | High | Noise 2 | Higher order state transition (magenta) | 0.493173 |
| 4 | 3 State | 3 | High | Noise 2 | Higher order state transition (magenta) | 0.604476 |
| 4 | 3 State | 1 | High | Noise 3 | Higher order state transition (magenta) | 0.320242 |
| 4 | 3 State | 2 | High | Noise 3 | Higher order state transition (magenta) | 0.301434 |
| 4 | 3 State | 3 | High | Noise 3 | Higher order state transition (magenta) | 0.10474 |
| 4 | 3 State | 1 | Low | Noise 1 | Higher order state transition (magenta) | 0.642313 |
| 4 | 3 State | 2 | Low | Noise 1 | Higher order state transition (magenta) | 0.658769 |
| 4 | 3 State | 3 | Low | Noise 1 | Higher order state transition (magenta) | 0.698815 |
| 4 | 3 State | 1 | Low | Noise 2 | Higher order state transition (magenta) | 0.577048 |
| 4 | 3 State | 2 | Low | Noise 2 | Higher order state transition (magenta) | 0.619435 |

| log10(c50) | Generating-Function | Replicate | Saturation | Noise-Profile | Classification | Confidence-Score |
|---|---|---|---|---|---|---|
| 4 | 3 State | 3 | Low | Noise 2 | Higher order state transition (magenta) | 0.619054 |
| 4 | 3 State | 1 | Low | Noise 3 | Higher order state transition (magenta) | 0.303053 |
| 4 | 3 State | 2 | Low | Noise 3 | Higher order state transition (magenta) | 0.389778 |
| 4 | 3 State | 3 | Low | Noise 3 | Higher order state transition (magenta) | 0.346246 |
| 4 | 3 State | 1 | Uniform 1 | Noise 1 | Higher order state transition (magenta) | 0.653063 |
| 4 | 3 State | 2 | Uniform 1 | Noise 1 | Higher order state transition (magenta) | 0.638787 |
| 4 | 3 State | 3 | Uniform 1 | Noise 1 | Higher order state transition (magenta) | 0.700642 |
| 4 | 3 State | 1 | Uniform 1 | Noise 2 | Higher order state transition (magenta) | 0.638101 |
| 4 | 3 State | 2 | Uniform 1 | Noise 2 | Higher order state transition (magenta) | 0.648085 |
| 4 | 3 State | 3 | Uniform 1 | Noise 2 | Higher order state transition (magenta) | 0.606547 |
| 4 | 3 State | 1 | Uniform 1 | Noise 3 | Higher order state transition (magenta) | 0.320728 |
| 4 | 3 State | 2 | Uniform 1 | Noise 3 | Higher order state transition (magenta) | 0.309023 |
| 4 | 3 State | 3 | Uniform 1 | Noise 3 | Higher order state transition (magenta) | 0.299869 |
| 6 | 3 State | 1 | High | Noise 1 | Higher order state transition (magenta) | 0.671944 |
| 6 | 3 State | 2 | High | Noise 1 | Higher order state transition (magenta) | 0.681096 |
| 6 | 3 State | 3 | High | Noise 1 | Higher order state transition (magenta) | 0.655612 |
| 6 | 3 State | 1 | High | Noise 2 | Higher order state transition (magenta) | 0.731309 |
| 6 | 3 State | 2 | High | Noise 2 | Higher order state transition (magenta) | 0.626682 |
| 6 | 3 State | 3 | High | Noise 2 | Higher order state transition (magenta) | 0.656763 |
| 6 | 3 State | 1 | High | Noise 3 | Higher order state transition (magenta) | 0.480278 |
| 6 | 3 State | 2 | High | Noise 3 | Higher order state transition (magenta) | 0.45685 |
| 6 | 3 State | 3 | High | Noise 3 | Higher order state transition (magenta) | 0.535074 |
| 6 | 3 State | 1 | Low | Noise 1 | Higher order state transition (magenta) | 0.666124 |
| 6 | 3 State | 2 | Low | Noise 1 | Higher order state transition (magenta) | 0.679163 |
| 6 | 3 State | 3 | Low | Noise 1 | Higher order state transition (magenta) | 0.635655 |
| 6 | 3 State | 1 | Low | Noise 2 | Higher order state transition (magenta) | 0.695505 |
| 6 | 3 State | 2 | Low | Noise 2 | Higher order state transition (magenta) | 0.737014 |
| 6 | 3 State | 3 | Low | Noise 2 | Higher order state transition (magenta) | 0.489918 |
| 6 | 3 State | 1 | Low | Noise 3 | Higher order state transition (magenta) | 0.458197 |
| 6 | 3 State | 2 | Low | Noise 3 | Higher order state transition (magenta) | 0.4259 |
| 6 | 3 State | 3 | Low | Noise 3 | Higher order state transition (magenta) | 0.391514 |
| 6 | 3 State | 1 | Uniform 1 | Noise 1 | Higher order state transition (magenta) | 0.693316 |

| log10(c50) | Generating-Function | Replicate | Saturation | Noise-Profile | Classification | Confidence-Score |
|---|---|---|---|---|---|---|
| 6 | 3 State | 2 | Uniform 1 | Noise 1 | Higher order state transition (magenta) | 0.653983 |
| 6 | 3 State | 3 | Uniform 1 | Noise 1 | Higher order state transition (magenta) | 0.677854 |
| 6 | 3 State | 1 | Uniform 1 | Noise 2 | Higher order state transition (magenta) | 0.715761 |
| 6 | 3 State | 2 | Uniform 1 | Noise 2 | Higher order state transition (magenta) | 0.700392 |
| 6 | 3 State | 3 | Uniform 1 | Noise 2 | Higher order state transition (magenta) | 0.564828 |
| 6 | 3 State | 1 | Uniform 1 | Noise 3 | Higher order state transition (magenta) | 0.479303 |
| 6 | 3 State | 2 | Uniform 1 | Noise 3 | Higher order state transition (magenta) | 0.497872 |
| 6 | 3 State | 3 | Uniform 1 | Noise 3 | Higher order state transition (magenta) | 0.44843 |
| 8 | 3 State | 1 | High | Noise 1 | Higher order state transition (magenta) | 0.132153 |
| 8 | 3 State | 2 | High | Noise 1 | Higher order state transition (magenta) | 0.330193 |
| 8 | 3 State | 3 | High | Noise 1 | Higher order state transition (magenta) | 0.0295417 |
| 8 | 3 State | 1 | High | Noise 2 | Higher order state transition (magenta) | 0.081692 |
| 8 | 3 State | 2 | High | Noise 2 | Higher order state transition (magenta) | 0.278568 |
| 8 | 3 State | 3 | High | Noise 2 | Two-state discontinuous transition (green) | 0.0117776 |
| 8 | 3 State | 1 | High | Noise 3 | Higher order state transition (magenta) | 0.459571 |
| 8 | 3 State | 2 | High | Noise 3 | Higher order state transition (magenta) | 0.269888 |
| 8 | 3 State | 3 | High | Noise 3 | Higher order state transition (magenta) | 0.190652 |
| 8 | 3 State | 1 | Low | Noise 1 | Higher order state transition (magenta) | 0.132774 |
| 8 | 3 State | 2 | Low | Noise 1 | Higher order state transition (magenta) | 0.0294251 |
| 8 | 3 State | 3 | Low | Noise 1 | Higher order state transition (magenta) | 0.248775 |
| 8 | 3 State | 1 | Low | Noise 2 | Higher order state transition (magenta) | 0.675253 |
| 8 | 3 State | 2 | Low | Noise 2 | Higher order state transition (magenta) | 0.560303 |
| 8 | 3 State | 3 | Low | Noise 2 | Higher order state transition (magenta) | 0.744742 |
| 8 | 3 State | 1 | Low | Noise 3 | Higher order state transition (magenta) | 0.515401 |
| 8 | 3 State | 2 | Low | Noise 3 | Higher order state transition (magenta) | 0.61651 |
| 8 | 3 State | 3 | Low | Noise 3 | Higher order state transition (magenta) | 0.416882 |
| 8 | 3 State | 1 | Uniform 1 | Noise 1 | Higher order state transition (magenta) | 0.243616 |
| 8 | 3 State | 2 | Uniform 1 | Noise 1 | Higher order state transition (magenta) | 0.327942 |
| 8 | 3 State | 3 | Uniform 1 | Noise 1 | Higher order state transition (magenta) | 0.109243 |
| 8 | 3 State | 1 | Uniform 1 | Noise 2 | Higher order state transition (magenta) | 0.172307 |
| 8 | 3 State | 2 | Uniform 1 | Noise 2 | Higher order state transition (magenta) | 0.452379 |
| 8 | 3 State | 3 | Uniform 1 | Noise 2 | Higher order state transition (magenta) | 0.606731 |

| log10(c50) | Generating-Function | Replicate | Saturation | Noise-Profile | Classification | Confidence-Score |
|---|---|---|---|---|---|---|
| 8 | 3 State | 1 | Uniform 1 | Noise 3 | Higher order state transition (magenta) | 0.412813 |
| 8 | 3 State | 2 | Uniform 1 | Noise 3 | Higher order state transition (magenta) | 0.292569 |
| 8 | 3 State | 3 | Uniform 1 | Noise 3 | Higher order state transition (magenta) | 0.341973 |
| 10 | 3 State | 1 | Uniform 2 | Noise 1 | Two-state discontinuous transition (green) | 0.160875 |
| 10 | 3 State | 2 | Uniform 2 | Noise 1 | Two-state discontinuous transition (green) | 0.41912 |
| 10 | 3 State | 3 | Uniform 2 | Noise 1 | Two-state discontinuous transition (green) | 0.458148 |
| 10 | 3 State | 1 | Uniform 2 | Noise 2 | Higher order state transition (magenta) | 0.558226 |
| 10 | 3 State | 2 | Uniform 2 | Noise 2 | Higher order state transition (magenta) | 0.0509299 |
| 10 | 3 State | 3 | Uniform 2 | Noise 2 | Higher order state transition (magenta) | 0.518843 |
| 10 | 3 State | 1 | Uniform 2 | Noise 3 | Two-state discontinuous transition (green) | 0.100431 |
| 10 | 3 State | 2 | Uniform 2 | Noise 3 | Higher order state transition (magenta) | 0.240856 |
| 10 | 3 State | 3 | Uniform 2 | Noise 3 | Two-state discontinuous transition (green) | 0.120229 |
| 12 | 3 State | 1 | Uniform 2 | Noise 1 | Two-state discontinuous transition (green) | 0.530231 |
| 12 | 3 State | 2 | Uniform 2 | Noise 1 | Two-state discontinuous transition (green) | 0.649674 |
| 12 | 3 State | 3 | Uniform 2 | Noise 1 | Two-state discontinuous transition (green) | 0.503427 |
| 12 | 3 State | 1 | Uniform 2 | Noise 2 | Higher order state transition (magenta) | 0.343287 |
| 12 | 3 State | 2 | Uniform 2 | Noise 2 | Two-state discontinuous transition (green) | 0.142396 |
| 12 | 3 State | 3 | Uniform 2 | Noise 2 | Higher order state transition (magenta) | 0.246489 |
| 12 | 3 State | 1 | Uniform 2 | Noise 3 | Two-state discontinuous transition (green) | 0.119629 |
| 12 | 3 State | 2 | Uniform 2 | Noise 3 | Two-state discontinuous transition (green) | 0.203664 |
| 12 | 3 State | 3 | Uniform 2 | Noise 3 | Two-state discontinuous transition (green) | 0.203691 |
| 1 | Sigmoid | 1 | High | Noise 1 | Assembled at all expression levels (black) | 0.999998 |
| 1 | Sigmoid | 2 | High | Noise 1 | Assembled at all expression levels (black) | 0.999997 |
| 1 | Sigmoid | 3 | High | Noise 1 | Assembled at all expression levels (black) | 0.999679 |
| 1 | Sigmoid | 1 | High | Noise 2 | Assembled at all expression levels (black) | 0.997261 |
| 1 | Sigmoid | 2 | High | Noise 2 | Assembled at all expression levels (black) | 0.994899 |
| 1 | Sigmoid | 3 | High | Noise 2 | Assembled at all expression levels (black) | 0.997667 |
| 1 | Sigmoid | 1 | High | Noise 3 | Assembled at all expression levels (black) | 0.97836 |
| 1 | Sigmoid | 2 | High | Noise 3 | Assembled at all expression levels (black) | 0.956675 |
| 1 | Sigmoid | 3 | High | Noise 3 | Assembled at all expression levels (black) | 0.92511 |
| 2 | Sigmoid | 1 | High | Noise 1 | Assembled at all expression levels (black) | 0.99999 |
| 2 | Sigmoid | 2 | High | Noise 1 | Assembled at all expression levels (black) | 1 |

| log10(c50) | Generating-Function | Replicate | Saturation | Noise-Profile | Classification | Confidence-Score |
|---|---|---|---|---|---|---|
| 2 | Sigmoid | 3 | High | Noise 1 | Assembled at all expression levels (black) | 0.999979 |
| 2 | Sigmoid | 1 | High | Noise 2 | Assembled at all expression levels (black) | 0.999996 |
| 2 | Sigmoid | 2 | High | Noise 2 | Assembled at all expression levels (black) | 0.993101 |
| 2 | Sigmoid | 3 | High | Noise 2 | Assembled at all expression levels (black) | 0.989398 |
| 2 | Sigmoid | 1 | High | Noise 3 | Assembled at all expression levels (black) | 0.96026 |
| 2 | Sigmoid | 2 | High | Noise 3 | Assembled at all expression levels (black) | 0.958562 |
| 2 | Sigmoid | 3 | High | Noise 3 | Assembled at all expression levels (black) | 0.917499 |
| 2 | Sigmoid | 1 | Low | Noise 1 | Assembled at all expression levels (black) | 0.999997 |
| 2 | Sigmoid | 2 | Low | Noise 1 | Assembled at all expression levels (black) | 0.999999 |
| 2 | Sigmoid | 3 | Low | Noise 1 | Assembled at all expression levels (black) | 1 |
| 2 | Sigmoid | 1 | Low | Noise 2 | Assembled at all expression levels (black) | 0.995477 |
| 2 | Sigmoid | 2 | Low | Noise 2 | Assembled at all expression levels (black) | 0.996187 |
| 2 | Sigmoid | 3 | Low | Noise 2 | Assembled at all expression levels (black) | 0.986597 |
| 2 | Sigmoid | 1 | Low | Noise 3 | Assembled at all expression levels (black) | 0.940585 |
| 2 | Sigmoid | 2 | Low | Noise 3 | Assembled at all expression levels (black) | 0.976755 |
| 2 | Sigmoid | 3 | Low | Noise 3 | Assembled at all expression levels (black) | 0.965287 |
| 4 | Sigmoid | 1 | High | Noise 1 | Two-state continuous transition (red) | 0.582868 |
| 4 | Sigmoid | 2 | High | Noise 1 | Two-state continuous transition (red) | 0.57489 |
| 4 | Sigmoid | 3 | High | Noise 1 | Two-state continuous transition (red) | 0.623844 |
| 4 | Sigmoid | 1 | High | Noise 2 | Two-state continuous transition (red) | 0.088625 |
| 4 | Sigmoid | 2 | High | Noise 2 | Two-state continuous transition (red) | 0.0636041 |
| 4 | Sigmoid | 3 | High | Noise 2 | Two-state continuous transition (red) | 0.0758606 |
| 4 | Sigmoid | 1 | High | Noise 3 | Two-state discontinuous transition (green) | 0.328957 |
| 4 | Sigmoid | 2 | High | Noise 3 | Two-state discontinuous transition (green) | 0.31025 |
| 4 | Sigmoid | 3 | High | Noise 3 | Two-state discontinuous transition (green) | 0.348495 |
| 4 | Sigmoid | 1 | Low | Noise 1 | Two-state continuous transition (red) | 0.587799 |
| 4 | Sigmoid | 2 | Low | Noise 1 | Two-state continuous transition (red) | 0.579076 |
| 4 | Sigmoid | 3 | Low | Noise 1 | Two-state continuous transition (red) | 0.578276 |
| 4 | Sigmoid | 1 | Low | Noise 2 | Two-state continuous transition (red) | 0.0663032 |
| 4 | Sigmoid | 2 | Low | Noise 2 | Two-state continuous transition (red) | 0.0814072 |
| 4 | Sigmoid | 3 | Low | Noise 2 | Two-state continuous transition (red) | 0.0222756 |
| 4 | Sigmoid | 1 | Low | Noise 3 | Two-state discontinuous transition (green) | 0.400209 |

| log10(c50) | Generating-Function | Replicate | Saturation | Noise-Profile | Classification | Confidence-Score |
|---|---|---|---|---|---|---|
| 4 | Sigmoid | 2 | Low | Noise 3 | Two-state discontinuous transition (green) | 0.320155 |
| 4 | Sigmoid | 3 | Low | Noise 3 | Two-state discontinuous transition (green) | 0.368697 |
| 4 | Sigmoid | 1 | Uniform 1 | Noise 1 | Two-state continuous transition (red) | 0.549374 |
| 4 | Sigmoid | 2 | Uniform 1 | Noise 1 | Two-state continuous transition (red) | 0.611786 |
| 4 | Sigmoid | 3 | Uniform 1 | Noise 1 | Two-state continuous transition (red) | 0.62269 |
| 4 | Sigmoid | 1 | Uniform 1 | Noise 2 | Two-state continuous transition (red) | 0.0953086 |
| 4 | Sigmoid | 2 | Uniform 1 | Noise 2 | Two-state continuous transition (red) | 0.0695092 |
| 4 | Sigmoid | 3 | Uniform 1 | Noise 2 | Two-state continuous transition (red) | 0.0623308 |
| 4 | Sigmoid | 1 | Uniform 1 | Noise 3 | Two-state discontinuous transition (green) | 0.373954 |
| 4 | Sigmoid | 2 | Uniform 1 | Noise 3 | Two-state discontinuous transition (green) | 0.31774 |
| 4 | Sigmoid | 3 | Uniform 1 | Noise 3 | Two-state discontinuous transition (green) | 0.349601 |
| 6 | Sigmoid | 1 | High | Noise 1 | Two-state continuous transition (red) | 0.665029 |
| 6 | Sigmoid | 2 | High | Noise 1 | Two-state continuous transition (red) | 0.677324 |
| 6 | Sigmoid | 3 | High | Noise 1 | Two-state continuous transition (red) | 0.700741 |
| 6 | Sigmoid | 1 | High | Noise 2 | Two-state continuous transition (red) | 0.36453 |
| 6 | Sigmoid | 2 | High | Noise 2 | Two-state continuous transition (red) | 0.35235 |
| 6 | Sigmoid | 3 | High | Noise 2 | Two-state continuous transition (red) | 0.415738 |
| 6 | Sigmoid | 1 | High | Noise 3 | Two-state continuous transition (red) | 0.0514515 |
| 6 | Sigmoid | 2 | High | Noise 3 | Two-state discontinuous transition (green) | 0.0169097 |
| 6 | Sigmoid | 3 | High | Noise 3 | Two-state continuous transition (red) | 0.0211382 |
| 6 | Sigmoid | 1 | Low | Noise 1 | Two-state continuous transition (red) | 0.676762 |
| 6 | Sigmoid | 2 | Low | Noise 1 | Two-state continuous transition (red) | 0.675855 |
| 6 | Sigmoid | 3 | Low | Noise 1 | Two-state continuous transition (red) | 0.669753 |
| 6 | Sigmoid | 1 | Low | Noise 2 | Two-state continuous transition (red) | 0.39621 |
| 6 | Sigmoid | 2 | Low | Noise 2 | Two-state continuous transition (red) | 0.405384 |
| 6 | Sigmoid | 3 | Low | Noise 2 | Two-state continuous transition (red) | 0.397039 |
| 6 | Sigmoid | 1 | Low | Noise 3 | Two-state continuous transition (red) | 0.0522461 |
| 6 | Sigmoid | 2 | Low | Noise 3 | Two-state continuous transition (red) | 0.0281929 |
| 6 | Sigmoid | 3 | Low | Noise 3 | Two-state continuous transition (red) | 0.0279934 |
| 6 | Sigmoid | 1 | Uniform 1 | Noise 1 | Two-state continuous transition (red) | 0.671927 |
| 6 | Sigmoid | 2 | Uniform 1 | Noise 1 | Two-state continuous transition (red) | 0.690882 |
| 6 | Sigmoid | 3 | Uniform 1 | Noise 1 | Two-state continuous transition (red) | 0.676993 |

| log10(c50) | Generating-Function | Replicate | Saturation | Noise-Profile | Classification | Confidence-Score |
|---|---|---|---|---|---|---|
| 6 | Sigmoid | 1 | Uniform 1 | Noise 2 | Two-state continuous transition (red) | 0.382539 |
| 6 | Sigmoid | 2 | Uniform 1 | Noise 2 | Two-state continuous transition (red) | 0.460024 |
| 6 | Sigmoid | 3 | Uniform 1 | Noise 2 | Two-state continuous transition (red) | 0.382118 |
| 6 | Sigmoid | 1 | Uniform 1 | Noise 3 | Two-state continuous transition (red) | 0.100853 |
| 6 | Sigmoid | 2 | Uniform 1 | Noise 3 | Two-state continuous transition (red) | 0.0236738 |
| 6 | Sigmoid | 3 | Uniform 1 | Noise 3 | Two-state continuous transition (red) | 0.0845262 |
| 8 | Sigmoid | 1 | High | Noise 1 | Two-state continuous transition (red) | 0.682486 |
| 8 | Sigmoid | 2 | High | Noise 1 | Two-state continuous transition (red) | 0.677143 |
| 8 | Sigmoid | 3 | High | Noise 1 | Two-state continuous transition (red) | 0.637041 |
| 8 | Sigmoid | 1 | High | Noise 2 | Two-state continuous transition (red) | 0.465439 |
| 8 | Sigmoid | 2 | High | Noise 2 | Two-state continuous transition (red) | 0.436313 |
| 8 | Sigmoid | 3 | High | Noise 2 | Two-state continuous transition (red) | 0.440201 |
| 8 | Sigmoid | 1 | High | Noise 3 | Two-state continuous transition (red) | 0.283482 |
| 8 | Sigmoid | 2 | High | Noise 3 | Two-state continuous transition (red) | 0.234956 |
| 8 | Sigmoid | 3 | High | Noise 3 | Two-state continuous transition (red) | 0.214676 |
| 8 | Sigmoid | 1 | Low | Noise 1 | Two-state continuous transition (red) | 0.668071 |
| 8 | Sigmoid | 2 | Low | Noise 1 | Two-state continuous transition (red) | 0.646335 |
| 8 | Sigmoid | 3 | Low | Noise 1 | Two-state continuous transition (red) | 0.643409 |
| 8 | Sigmoid | 1 | Low | Noise 2 | Two-state continuous transition (red) | 0.466111 |
| 8 | Sigmoid | 2 | Low | Noise 2 | Two-state continuous transition (red) | 0.411876 |
| 8 | Sigmoid | 3 | Low | Noise 2 | Two-state continuous transition (red) | 0.471386 |
| 8 | Sigmoid | 1 | Low | Noise 3 | Two-state continuous transition (red) | 0.223226 |
| 8 | Sigmoid | 2 | Low | Noise 3 | Two-state continuous transition (red) | 0.169346 |
| 8 | Sigmoid | 3 | Low | Noise 3 | Two-state continuous transition (red) | 0.188756 |
| 8 | Sigmoid | 1 | Uniform 1 | Noise 1 | Two-state continuous transition (red) | 0.640101 |
| 8 | Sigmoid | 2 | Uniform 1 | Noise 1 | Two-state continuous transition (red) | 0.64232 |
| 8 | Sigmoid | 3 | Uniform 1 | Noise 1 | Two-state continuous transition (red) | 0.646699 |
| 8 | Sigmoid | 1 | Uniform 1 | Noise 2 | Two-state continuous transition (red) | 0.446686 |
| 8 | Sigmoid | 2 | Uniform 1 | Noise 2 | Two-state continuous transition (red) | 0.468282 |
| 8 | Sigmoid | 3 | Uniform 1 | Noise 2 | Two-state continuous transition (red) | 0.446119 |
| 8 | Sigmoid | 1 | Uniform 1 | Noise 3 | Two-state continuous transition (red) | 0.300605 |
| 8 | Sigmoid | 2 | Uniform 1 | Noise 3 | Two-state continuous transition (red) | 0.197703 |

| log10(c50) | Generating-Function | Replicate | Saturation | Noise-Profile | Classification | Confidence-Score |
|---|---|---|---|---|---|---|
| 8 | Sigmoid | 3 | Uniform 1 | Noise 3 | Two-state continuous transition (red) | 0.156698 |
| 10 | Sigmoid | 1 | High | Noise 1 | Higher order state transition (magenta) | 0.065959 |
| 10 | Sigmoid | 2 | High | Noise 1 | Higher order state transition (magenta) | 0.248574 |
| 10 | Sigmoid | 3 | High | Noise 1 | Two-state continuous transition (red) | 0.21258 |
| 10 | Sigmoid | 1 | High | Noise 2 | Higher order state transition (magenta) | 0.028059 |
| 10 | Sigmoid | 2 | High | Noise 2 | Higher order state transition (magenta) | 0.0235942 |
| 10 | Sigmoid | 3 | High | Noise 2 | Higher order state transition (magenta) | 0.127148 |
| 10 | Sigmoid | 1 | High | Noise 3 | Higher order state transition (magenta) | 0.23705 |
| 10 | Sigmoid | 2 | High | Noise 3 | Two-state discontinuous transition (green) | 0.10443 |
| 10 | Sigmoid | 3 | High | Noise 3 | Higher order state transition (magenta) | 0.0217016 |
| 10 | Sigmoid | 1 | Low | Noise 1 | Two-state continuous transition (red) | 0.261796 |
| 10 | Sigmoid | 2 | Low | Noise 1 | Two-state continuous transition (red) | 0.243962 |
| 10 | Sigmoid | 3 | Low | Noise 1 | Two-state continuous transition (red) | 0.170127 |
| 10 | Sigmoid | 1 | Low | Noise 2 | Higher order state transition (magenta) | 0.104578 |
| 10 | Sigmoid | 2 | Low | Noise 2 | Higher order state transition (magenta) | 0.00848424 |
| 10 | Sigmoid | 3 | Low | Noise 2 | Higher order state transition (magenta) | 0.0558006 |
| 10 | Sigmoid | 1 | Low | Noise 3 | Higher order state transition (magenta) | 0.223217 |
| 10 | Sigmoid | 2 | Low | Noise 3 | Higher order state transition (magenta) | 0.0550116 |
| 10 | Sigmoid | 3 | Low | Noise 3 | Higher order state transition (magenta) | 0.0093684 |
| 10 | Sigmoid | 1 | Uniform 2 | Noise 1 | Two-state continuous transition (red) | 0.223953 |
| 10 | Sigmoid | 2 | Uniform 2 | Noise 1 | Two-state continuous transition (red) | 0.244822 |
| 10 | Sigmoid | 3 | Uniform 2 | Noise 1 | Two-state continuous transition (red) | 0.283781 |
| 10 | Sigmoid | 1 | Uniform 2 | Noise 2 | Higher order state transition (magenta) | 0.14233 |
| 10 | Sigmoid | 2 | Uniform 2 | Noise 2 | Higher order state transition (magenta) | 0.0513458 |
| 10 | Sigmoid | 3 | Uniform 2 | Noise 2 | Higher order state transition (magenta) | 0.0281355 |
| 10 | Sigmoid | 1 | Uniform 2 | Noise 3 | Higher order state transition (magenta) | 0.238799 |
| 10 | Sigmoid | 2 | Uniform 2 | Noise 3 | Higher order state transition (magenta) | 0.116542 |
| 10 | Sigmoid | 3 | Uniform 2 | Noise 3 | Higher order state transition (magenta) | 0.105537 |
| 12 | Sigmoid | 1 | Uniform 2 | Noise 1 | Two-state discontinuous transition (green) | 0.0304707 |
| 12 | Sigmoid | 2 | Uniform 2 | Noise 1 | No assembly at all expression levels (blue) | 0.999637 |
| 12 | Sigmoid | 3 | Uniform 2 | Noise 1 | No assembly at all expression levels (blue) | 0.03979 |
| 12 | Sigmoid | 1 | Uniform 2 | Noise 2 | No assembly at all expression levels (blue) | 0.999942 |

| log10(c50) | Generating-Function | Replicate | Saturation | Noise-Profile | Classification | Confidence-Score |
|---|---|---|---|---|---|---|
| 12 | Sigmoid | 2 | Uniform 2 | Noise 2 | No assembly at all expression levels (blue) | 1 |
| 12 | Sigmoid | 3 | Uniform 2 | Noise 2 | No assembly at all expression levels (blue) | 0.99986 |
| 12 | Sigmoid | 1 | Uniform 2 | Noise 3 | No assembly at all expression levels (blue) | 0.996983 |
| 12 | Sigmoid | 2 | Uniform 2 | Noise 3 | No assembly at all expression levels (blue) | 1 |
| 12 | Sigmoid | 3 | Uniform 2 | Noise 3 | No assembly at all expression levels (blue) | 0.999466 |
| 1 | Step | 1 | High | Noise 1 | Assembled at all expression levels (black) | 0.999686 |
| 1 | Step | 2 | High | Noise 1 | Assembled at all expression levels (black) | 0.999982 |
| 1 | Step | 3 | High | Noise 1 | Assembled at all expression levels (black) | 0.999683 |
| 1 | Step | 1 | High | Noise 2 | Assembled at all expression levels (black) | 0.997853 |
| 1 | Step | 2 | High | Noise 2 | Assembled at all expression levels (black) | 0.999997 |
| 1 | Step | 3 | High | Noise 2 | Assembled at all expression levels (black) | 0.999989 |
| 1 | Step | 1 | High | Noise 3 | Assembled at all expression levels (black) | 0.94387 |
| 1 | Step | 2 | High | Noise 3 | Assembled at all expression levels (black) | 0.983098 |
| 1 | Step | 3 | High | Noise 3 | Assembled at all expression levels (black) | 0.993609 |
| 2 | Step | 1 | High | Noise 1 | Assembled at all expression levels (black) | 0.999684 |
| 2 | Step | 2 | High | Noise 1 | Assembled at all expression levels (black) | 0.999685 |
| 2 | Step | 3 | High | Noise 1 | Assembled at all expression levels (black) | 0.999687 |
| 2 | Step | 1 | High | Noise 2 | Assembled at all expression levels (black) | 0.99999 |
| 2 | Step | 2 | High | Noise 2 | Assembled at all expression levels (black) | 0.996139 |
| 2 | Step | 3 | High | Noise 2 | Assembled at all expression levels (black) | 0.99958 |
| 2 | Step | 1 | High | Noise 3 | Assembled at all expression levels (black) | 0.980209 |
| 2 | Step | 2 | High | Noise 3 | Assembled at all expression levels (black) | 0.985168 |
| 2 | Step | 3 | High | Noise 3 | Assembled at all expression levels (black) | 0.977156 |
| 2 | Step | 1 | Low | Noise 1 | Assembled at all expression levels (black) | 0.999686 |
| 2 | Step | 2 | Low | Noise 1 | Assembled at all expression levels (black) | 0.999684 |
| 2 | Step | 3 | Low | Noise 1 | Assembled at all expression levels (black) | 0.999685 |
| 2 | Step | 1 | Low | Noise 2 | Assembled at all expression levels (black) | 0.994105 |
| 2 | Step | 2 | Low | Noise 2 | Assembled at all expression levels (black) | 0.995933 |
| 2 | Step | 3 | Low | Noise 2 | Assembled at all expression levels (black) | 0.999732 |
| 2 | Step | 1 | Low | Noise 3 | Assembled at all expression levels (black) | 0.98579 |
| 2 | Step | 2 | Low | Noise 3 | Assembled at all expression levels (black) | 0.975905 |
| 2 | Step | 3 | Low | Noise 3 | Assembled at all expression levels (black) | 0.988926 |

| log10(c50) | Generating-Function | Replicate | Saturation | Noise-Profile | Classification | Confidence-Score |
|---|---|---|---|---|---|---|
| 4 | Step | 1 | High | Noise 1 | Two-state discontinuous transition (green) | 0.527439 |
| 4 | Step | 2 | High | Noise 1 | Two-state discontinuous transition (green) | 0.549141 |
| 4 | Step | 3 | High | Noise 1 | Two-state discontinuous transition (green) | 0.535265 |
| 4 | Step | 1 | High | Noise 2 | Two-state discontinuous transition (green) | 0.819632 |
| 4 | Step | 2 | High | Noise 2 | Two-state discontinuous transition (green) | 0.738042 |
| 4 | Step | 3 | High | Noise 2 | Two-state discontinuous transition (green) | 0.842044 |
| 4 | Step | 1 | High | Noise 3 | Two-state discontinuous transition (green) | 0.639317 |
| 4 | Step | 2 | High | Noise 3 | Two-state discontinuous transition (green) | 0.6174 |
| 4 | Step | 3 | High | Noise 3 | Two-state discontinuous transition (green) | 0.911071 |
| 4 | Step | 1 | Low | Noise 1 | Two-state discontinuous transition (green) | 0.540014 |
| 4 | Step | 2 | Low | Noise 1 | Two-state discontinuous transition (green) | 0.526254 |
| 4 | Step | 3 | Low | Noise 1 | Two-state discontinuous transition (green) | 0.512421 |
| 4 | Step | 1 | Low | Noise 2 | Two-state discontinuous transition (green) | 0.871783 |
| 4 | Step | 2 | Low | Noise 2 | Two-state discontinuous transition (green) | 0.841139 |
| 4 | Step | 3 | Low | Noise 2 | Two-state discontinuous transition (green) | 0.807766 |
| 4 | Step | 1 | Low | Noise 3 | Two-state discontinuous transition (green) | 0.37312 |
| 4 | Step | 2 | Low | Noise 3 | Two-state discontinuous transition (green) | 0.624564 |
| 4 | Step | 3 | Low | Noise 3 | Two-state discontinuous transition (green) | 0.920083 |
| 4 | Step | 1 | Uniform 1 | Noise 1 | Two-state discontinuous transition (green) | 0.538637 |
| 4 | Step | 2 | Uniform 1 | Noise 1 | Two-state discontinuous transition (green) | 0.514547 |
| 4 | Step | 3 | Uniform 1 | Noise 1 | Two-state discontinuous transition (green) | 0.506274 |
| 4 | Step | 1 | Uniform 1 | Noise 2 | Two-state discontinuous transition (green) | 0.800145 |
| 4 | Step | 2 | Uniform 1 | Noise 2 | Two-state discontinuous transition (green) | 0.804407 |
| 4 | Step | 3 | Uniform 1 | Noise 2 | Two-state discontinuous transition (green) | 0.833043 |
| 4 | Step | 1 | Uniform 1 | Noise 3 | Two-state discontinuous transition (green) | 0.685381 |
| 4 | Step | 2 | Uniform 1 | Noise 3 | Two-state discontinuous transition (green) | 0.869528 |
| 4 | Step | 3 | Uniform 1 | Noise 3 | Two-state discontinuous transition (green) | 0.675083 |
| 6 | Step | 1 | High | Noise 1 | Two-state discontinuous transition (green) | 0.49362 |
| 6 | Step | 2 | High | Noise 1 | Two-state discontinuous transition (green) | 0.540372 |
| 6 | Step | 3 | High | Noise 1 | Two-state discontinuous transition (green) | 0.546093 |
| 6 | Step | 1 | High | Noise 2 | Two-state discontinuous transition (green) | 0.836041 |
| 6 | Step | 2 | High | Noise 2 | Two-state discontinuous transition (green) | 0.850862 |

| log10(c50) | Generating-Function | Replicate | Saturation | Noise-Profile | Classification | Confidence-Score |
|---|---|---|---|---|---|---|
| 6 | Step | 3 | High | Noise 2 | Two-state discontinuous transition (green) | 0.833631 |
| 6 | Step | 1 | High | Noise 3 | Two-state discontinuous transition (green) | 0.940245 |
| 6 | Step | 2 | High | Noise 3 | Two-state discontinuous transition (green) | 0.896513 |
| 6 | Step | 3 | High | Noise 3 | Two-state discontinuous transition (green) | 0.921404 |
| 6 | Step | 1 | Low | Noise 1 | Two-state discontinuous transition (green) | 0.520163 |
| 6 | Step | 2 | Low | Noise 1 | Two-state discontinuous transition (green) | 0.52087 |
| 6 | Step | 3 | Low | Noise 1 | Two-state discontinuous transition (green) | 0.530269 |
| 6 | Step | 1 | Low | Noise 2 | Two-state discontinuous transition (green) | 0.84703 |
| 6 | Step | 2 | Low | Noise 2 | Two-state discontinuous transition (green) | 0.845815 |
| 6 | Step | 3 | Low | Noise 2 | Two-state discontinuous transition (green) | 0.849417 |
| 6 | Step | 1 | Low | Noise 3 | Two-state discontinuous transition (green) | 0.932094 |
| 6 | Step | 2 | Low | Noise 3 | Two-state discontinuous transition (green) | 0.911229 |
| 6 | Step | 3 | Low | Noise 3 | Two-state discontinuous transition (green) | 0.926136 |
| 6 | Step | 1 | Uniform 1 | Noise 1 | Two-state discontinuous transition (green) | 0.503459 |
| 6 | Step | 2 | Uniform 1 | Noise 1 | Two-state discontinuous transition (green) | 0.548664 |
| 6 | Step | 3 | Uniform 1 | Noise 1 | Two-state discontinuous transition (green) | 0.560756 |
| 6 | Step | 1 | Uniform 1 | Noise 2 | Two-state discontinuous transition (green) | 0.845126 |
| 6 | Step | 2 | Uniform 1 | Noise 2 | Two-state discontinuous transition (green) | 0.870816 |
| 6 | Step | 3 | Uniform 1 | Noise 2 | Two-state discontinuous transition (green) | 0.848562 |
| 6 | Step | 1 | Uniform 1 | Noise 3 | Two-state discontinuous transition (green) | 0.869073 |
| 6 | Step | 2 | Uniform 1 | Noise 3 | Two-state discontinuous transition (green) | 0.923083 |
| 6 | Step | 3 | Uniform 1 | Noise 3 | Two-state discontinuous transition (green) | 0.905966 |
| 8 | Step | 1 | High | Noise 1 | Two-state discontinuous transition (green) | 0.552718 |
| 8 | Step | 2 | High | Noise 1 | Two-state discontinuous transition (green) | 0.52334 |
| 8 | Step | 3 | High | Noise 1 | Two-state discontinuous transition (green) | 0.524608 |
| 8 | Step | 1 | High | Noise 2 | Two-state discontinuous transition (green) | 0.825563 |
| 8 | Step | 2 | High | Noise 2 | Two-state discontinuous transition (green) | 0.854495 |
| 8 | Step | 3 | High | Noise 2 | Two-state discontinuous transition (green) | 0.844215 |
| 8 | Step | 1 | High | Noise 3 | Two-state discontinuous transition (green) | 0.916052 |
| 8 | Step | 2 | High | Noise 3 | Two-state discontinuous transition (green) | 0.908063 |
| 8 | Step | 3 | High | Noise 3 | Two-state discontinuous transition (green) | 0.909361 |
| 8 | Step | 1 | Low | Noise 1 | Two-state discontinuous transition (green) | 0.566237 |

| log10(c50) | Generating-Function | Replicate | Saturation | Noise-Profile | Classification | Confidence-Score |
|---|---|---|---|---|---|---|
| 8 | Step | 2 | Low | Noise 1 | Two-state discontinuous transition (green) | 0.54619 |
| 8 | Step | 3 | Low | Noise 1 | Two-state discontinuous transition (green) | 0.53079 |
| 8 | Step | 1 | Low | Noise 2 | Two-state discontinuous transition (green) | 0.83522 |
| 8 | Step | 2 | Low | Noise 2 | Two-state discontinuous transition (green) | 0.869549 |
| 8 | Step | 3 | Low | Noise 2 | Two-state discontinuous transition (green) | 0.871142 |
| 8 | Step | 1 | Low | Noise 3 | Two-state discontinuous transition (green) | 0.918619 |
| 8 | Step | 2 | Low | Noise 3 | Two-state discontinuous transition (green) | 0.929365 |
| 8 | Step | 3 | Low | Noise 3 | Two-state discontinuous transition (green) | 0.938753 |
| 8 | Step | 1 | Uniform 1 | Noise 1 | Two-state discontinuous transition (green) | 0.529859 |
| 8 | Step | 2 | Uniform 1 | Noise 1 | Two-state discontinuous transition (green) | 0.609937 |
| 8 | Step | 3 | Uniform 1 | Noise 1 | Two-state discontinuous transition (green) | 0.561395 |
| 8 | Step | 1 | Uniform 1 | Noise 2 | Two-state discontinuous transition (green) | 0.864244 |
| 8 | Step | 2 | Uniform 1 | Noise 2 | Two-state discontinuous transition (green) | 0.831551 |
| 8 | Step | 3 | Uniform 1 | Noise 2 | Two-state discontinuous transition (green) | 0.853081 |
| 8 | Step | 1 | Uniform 1 | Noise 3 | Two-state discontinuous transition (green) | 0.884264 |
| 8 | Step | 2 | Uniform 1 | Noise 3 | Two-state discontinuous transition (green) | 0.900825 |
| 8 | Step | 3 | Uniform 1 | Noise 3 | Two-state discontinuous transition (green) | 0.926658 |
| 10 | Step | 1 | High | Noise 1 | Two-state discontinuous transition (green) | 0.493287 |
| 10 | Step | 2 | High | Noise 1 | Two-state discontinuous transition (green) | 0.437141 |
| 10 | Step | 3 | High | Noise 1 | Two-state discontinuous transition (green) | 0.455513 |
| 10 | Step | 1 | High | Noise 2 | Two-state discontinuous transition (green) | 0.60051 |
| 10 | Step | 2 | High | Noise 2 | Two-state discontinuous transition (green) | 0.629702 |
| 10 | Step | 3 | High | Noise 2 | Two-state discontinuous transition (green) | 0.712188 |
| 10 | Step | 1 | High | Noise 3 | Two-state discontinuous transition (green) | 0.407975 |
| 10 | Step | 2 | High | Noise 3 | Two-state discontinuous transition (green) | 0.446893 |
| 10 | Step | 3 | High | Noise 3 | Two-state discontinuous transition (green) | 0.401275 |
| 10 | Step | 1 | Low | Noise 1 | Two-state discontinuous transition (green) | 0.510572 |
| 10 | Step | 2 | Low | Noise 1 | Two-state discontinuous transition (green) | 0.412006 |
| 10 | Step | 3 | Low | Noise 1 | Two-state discontinuous transition (green) | 0.408051 |
| 10 | Step | 1 | Low | Noise 2 | Two-state discontinuous transition (green) | 0.628384 |
| 10 | Step | 2 | Low | Noise 2 | Two-state discontinuous transition (green) | 0.508627 |
| 10 | Step | 3 | Low | Noise 2 | Two-state discontinuous transition (green) | 0.459173 |

| log10(c50) | Generating-Function | Replicate | Saturation | Noise-Profile | Classification | Confidence-Score |
|---|---|---|---|---|---|---|
| 10 | Step | 1 | Low | Noise 3 | Two-state discontinuous transition (green) | 0.399977 |
| 10 | Step | 2 | Low | Noise 3 | Two-state discontinuous transition (green) | 0.388248 |
| 10 | Step | 3 | Low | Noise 3 | Two-state discontinuous transition (green) | 0.523575 |
| 10 | Step | 1 | Uniform 2 | Noise 1 | Two-state discontinuous transition (green) | 0.514575 |
| 10 | Step | 2 | Uniform 2 | Noise 1 | Two-state discontinuous transition (green) | 0.444036 |
| 10 | Step | 3 | Uniform 2 | Noise 1 | Two-state discontinuous transition (green) | 0.331044 |
| 10 | Step | 1 | Uniform 2 | Noise 2 | Two-state discontinuous transition (green) | 0.565756 |
| 10 | Step | 2 | Uniform 2 | Noise 2 | Two-state discontinuous transition (green) | 0.445484 |
| 10 | Step | 3 | Uniform 2 | Noise 2 | Two-state discontinuous transition (green) | 0.77615 |
| 10 | Step | 1 | Uniform 2 | Noise 3 | Two-state discontinuous transition (green) | 0.463704 |
| 10 | Step | 2 | Uniform 2 | Noise 3 | Two-state discontinuous transition (green) | 0.462512 |
| 10 | Step | 3 | Uniform 2 | Noise 3 | Two-state discontinuous transition (green) | 0.519952 |
| 12 | Step | 1 | Uniform 2 | Noise 1 | No assembly at all expression levels (blue) | 0.99983 |
| 12 | Step | 2 | Uniform 2 | Noise 1 | No assembly at all expression levels (blue) | 1 |
| 12 | Step | 3 | Uniform 2 | Noise 1 | No assembly at all expression levels (blue) | 0.999914 |
| 12 | Step | 1 | Uniform 2 | Noise 2 | No assembly at all expression levels (blue) | 1 |
| 12 | Step | 2 | Uniform 2 | Noise 2 | No assembly at all expression levels (blue) | 0.99994 |
| 12 | Step | 3 | Uniform 2 | Noise 2 | No assembly at all expression levels (blue) | 1 |
| 12 | Step | 1 | Uniform 2 | Noise 3 | Higher order state transition (magenta) | 0.00449382 |
| 12 | Step | 2 | Uniform 2 | Noise 3 | No assembly at all expression levels (blue) | 1 |
| 12 | Step | 3 | Uniform 2 | Noise 3 | No assembly at all expression levels (blue) | 1 |

**Table S2: Classifications of the cPrDs.** Here, $\log_{10}(c_{50})$ is only calculated for those DAmFRET histograms that show a transition. The "Confidence-Score" denotes how confident we are in each of the classifications as described in the Materials and Methods section.

| Gene | Replicate | log10(c50) | Classification | Confidence-Score |
|---|---|---|---|---|
| EPL1 | 1 | N/A | No assembly at all concentrations (Blue Class) | 1 |
| EPL1 | 2 | N/A | No assembly at all concentrations (Blue Class) | 1 |
| EPL1 | 3 | N/A | No assembly at all concentrations (Blue Class) | 1 |
| EPL1 | 4 | N/A | No assembly at all concentrations (Blue Class) | 1 |
| NRD1 | 1 | N/A | No assembly at all concentrations (Blue Class) | 1 |
| NRD1 | 2 | N/A | No assembly at all concentrations (Blue Class) | 0.5437 |
| NRD1 | 3 | N/A | No assembly at all concentrations (Blue Class) | 0.80726 |
| NRD1 | 4 | N/A | No assembly at all concentrations (Blue Class) | 1 |
| SLT2 | 1 | N/A | No assembly at all concentrations (Blue Class) | 0.99847 |
| SLT2 | 2 | N/A | No assembly at all concentrations (Blue Class) | 1 |
| SLT2 | 3 | N/A | No assembly at all concentrations (Blue Class) | 0.99911 |
| SLT2 | 4 | N/A | No assembly at all concentrations (Blue Class) | 1 |
| SAP30 | 1 | 4.89634 | Incomplete Transition (Yellow Class) | 0.13802 |
| SAP30 | 2 | 4.45529 | Incomplete Transition (Yellow Class) | 0.31665 |
| SAP30 | 3 | 4.14964 | No assembly at all concentrations (Blue Class) | 1 |
| SAP30 | 4 | 4.55432 | No assembly at all concentrations (Blue Class) | 1 |
| SCD6 | 1 | N/A | No assembly at all concentrations (Blue Class) | 0.99557 |
| SCD6 | 2 | N/A | No assembly at all concentrations (Blue Class) | 1 |
| SCD6 | 3 | N/A | No assembly at all concentrations (Blue Class) | 1 |
| SCD6 | 4 | N/A | No assembly at all concentrations (Blue Class) | 1 |
| SKG6 | 1 | N/A | No assembly at all concentrations (Blue Class) | 1 |
| SKG6 | 2 | N/A | No assembly at all concentrations (Blue Class) | 1 |
| SKG6 | 3 | N/A | No assembly at all concentrations (Blue Class) | 1 |
| SKG6 | 4 | N/A | No assembly at all concentrations (Blue Class) | 1 |
| PDR1 | 1 | 3.9755 | Incomplete Transition (Yellow Class) | 0.85186 |
| PDR1 | 2 | 3.93765 | Incomplete Transition (Yellow Class) | 0.70634 |
| PDR1 | 3 | 3.90418 | Incomplete Transition (Yellow Class) | 0.67611 |
| PDR1 | 4 | 4.04877 | Incomplete Transition (Yellow Class) | 0.93098 |
| CLG1 | 1 | 3.21112 | No assembly at all concentrations (Blue Class) | 1 |
| CLG1 | 2 | 2.96033 | No assembly at all concentrations (Blue Class) | 1 |
| CLG1 | 3 | 3.0682 | Incomplete Transition (Yellow Class) | 0.99026 |
| CLG1 | 4 | 3.32408 | Incomplete Transition (Yellow Class) | 0.99539 |

| Gene | Replicate | log10(c50) | Classification | Confidence-Score |
|---|---|---|---|---|
| TAF12 | 1 | N/A | No assembly at all concentrations (Blue Class) | 1 |
| TAF12 | 2 | N/A | No assembly at all concentrations (Blue Class) | 1 |
| TAF12 | 3 | N/A | No assembly at all concentrations (Blue Class) | 1 |
| TAF12 | 4 | N/A | No assembly at all concentrations (Blue Class) | 1 |
| PIN3 | 1 | N/A | No assembly at all concentrations (Blue Class) | 1 |
| PIN3 | 2 | N/A | No assembly at all concentrations (Blue Class) | 1 |
| PIN3 | 3 | N/A | No assembly at all concentrations (Blue Class) | 0.98588 |
| PIN3 | 4 | N/A | No assembly at all concentrations (Blue Class) | 1 |
| MRN1 | 1 | 2.72915 | Discontinuous Transition (Green Class) | 0.08591 |
| MRN1 | 2 | 2.73075 | Continuous Transition (Red Class) | 0.05195 |
| MRN1 | 3 | 2.75452 | Continuous Transition (Red Class) | 0.33956 |
| MRN1 | 4 | 2.78835 | Continuous Transition (Red Class) | 0.09381 |
| RLM1 | 1 | 3.96469 | Incomplete Transition (Yellow Class) | 0.92604 |
| RLM1 | 2 | 3.8242 | Incomplete Transition (Yellow Class) | 0.84133 |
| RLM1 | 3 | 3.59836 | Incomplete Transition (Yellow Class) | 0.97448 |
| RLM1 | 4 | 3.62472 | Incomplete Transition (Yellow Class) | 0.97394 |
| RAT1 | 1 | N/A | No assembly at all concentrations (Blue Class) | 1 |
| RAT1 | 2 | N/A | No assembly at all concentrations (Blue Class) | 1 |
| RAT1 | 3 | 3.69647 | No assembly at all concentrations (Blue Class) | 1 |
| RAT1 | 4 | N/A | No assembly at all concentrations (Blue Class) | 0.96488 |
| PDC2 | 1 | N/A | No assembly at all concentrations (Blue Class) | 1 |
| PDC2 | 2 | N/A | No assembly at all concentrations (Blue Class) | 0.99987 |
| PDC2 | 3 | N/A | No assembly at all concentrations (Blue Class) | 0.99962 |
| PDC2 | 4 | N/A | No assembly at all concentrations (Blue Class) | 1 |
| GLN3 | 1 | 2.67471 | Continuous Transition (Red Class) | 0.04627 |
| GLN3 | 2 | 2.65183 | Continuous Transition (Red Class) | 0.05413 |
| GLN3 | 3 | 2.61877 | Continuous Transition (Red Class) | 0.01888 |
| GLN3 | 4 | 2.68507 | Continuous Transition (Red Class) | 0.01034 |
| NEW1 | 1 | 2.31905 | Discontinuous Transition (Green Class) | 0.08652 |
| NEW1 | 2 | 2.18216 | Discontinuous Transition (Green Class) | 0.02195 |
| NEW1 | 3 | 2.21344 | Discontinuous Transition (Green Class) | 0.22294 |
| NEW1 | 4 | 2.26084 | Discontinuous Transition (Green Class) | 0.04384 |

| Gene | Replicate | log10(c50) | Classification | Confidence-Score |
|---|---|---|---|---|
| ASG1 | 1 | N/A | Assembled at all concentrations (Black Class) | 0.99982 |
| ASG1 | 2 | N/A | No assembly at all concentrations (Blue Class) | 1 |
| ASG1 | 3 | N/A | No assembly at all concentrations (Blue Class) | 1 |
| ASG1 | 4 | N/A | No assembly at all concentrations (Blue Class) | 1 |
| PCF11 | 1 | 3.44441 | No assembly at all concentrations (Blue Class) | 1 |
| PCF11 | 2 | N/A | No assembly at all concentrations (Blue Class) | 1 |
| PCF11 | 3 | N/A | No assembly at all concentrations (Blue Class) | 1 |
| PCF11 | 4 | N/A | No assembly at all concentrations (Blue Class) | 1 |
| PUF4 | 1 | 4.49992 | No assembly at all concentrations (Blue Class) | 1 |
| PUF4 | 2 | N/A | No assembly at all concentrations (Blue Class) | 0.94218 |
| PUF4 | 3 | N/A | No assembly at all concentrations (Blue Class) | 0.99755 |
| PUF4 | 4 | N/A | No assembly at all concentrations (Blue Class) | 1 |
| KSP1 | 1 | 2.98527 | Incomplete Transition (Yellow Class) | 0.51979 |
| KSP1 | 2 | 3.22157 | Incomplete Transition (Yellow Class) | 0.59687 |
| KSP1 | 3 | 2.92111 | Discontinuous Transition (Green Class) | 0.0548 |
| KSP1 | 4 | 3.08364 | Incomplete Transition (Yellow Class) | 0.70802 |
| YAK1 | 1 | N/A | No assembly at all concentrations (Blue Class) | 1 |
| YAK1 | 2 | N/A | No assembly at all concentrations (Blue Class) | 1 |
| YAK1 | 3 | N/A | No assembly at all concentrations (Blue Class) | 1 |
| YAK1 | 4 | N/A | No assembly at all concentrations (Blue Class) | 1 |
| PUB1 | 1 | 2.91398 | Continuous Transition (Red Class) | 0.05802 |
| PUB1 | 2 | 2.75553 | Discontinuous Transition (Green Class) | 0.14562 |
| PUB1 | 3 | 2.8276 | Discontinuous Transition (Green Class) | 0.06111 |
| PUB1 | 4 | 2.89928 | Continuous Transition (Red Class) | 0.01423 |
| GTS1 | 1 | 2.10756 | Continuous Transition (Red Class) | 0.22661 |
| GTS1 | 2 | 2.21239 | Continuous Transition (Red Class) | 0.14278 |
| GTS1 | 3 | 2.17367 | Continuous Transition (Red Class) | 0.04612 |
| GTS1 | 4 | 2.09999 | Continuous Transition (Red Class) | 0.13173 |
| MED2 | 1 | N/A | No assembly at all concentrations (Blue Class) | 1 |
| MED2 | 2 | N/A | No assembly at all concentrations (Blue Class) | 1 |
| MED2 | 3 | N/A | No assembly at all concentrations (Blue Class) | 1 |
| MED2 | 4 | N/A | No assembly at all concentrations (Blue Class) | 1 |

| Gene | Replicate | log10(c50) | Classification | Confidence-Score |
|---|---|---|---|---|
| RIM4 | 1 | 3.12192 | Incomplete Transition (Yellow Class) | 0.64586 |
| RIM4 | 2 | 3.02762 | Incomplete Transition (Yellow Class) | 0.72864 |
| RIM4 | 3 | 2.9832 | Incomplete Transition (Yellow Class) | 0.58787 |
| RIM4 | 4 | 3.11439 | Incomplete Transition (Yellow Class) | 0.88375 |
| URE2 | 1 | 4.33175 | Incomplete Transition (Yellow Class) | 0.98621 |
| URE2 | 2 | 4.24693 | Incomplete Transition (Yellow Class) | 0.94034 |
| URE2 | 3 | 4.28272 | Incomplete Transition (Yellow Class) | 0.99414 |
| URE2 | 4 | 4.21487 | Incomplete Transition (Yellow Class) | 0.99087 |
| GAL11 | 1 | N/A | No assembly at all concentrations (Blue Class) | 0.99988 |
| GAL11 | 2 | N/A | No assembly at all concentrations (Blue Class) | 0.9626 |
| GAL11 | 3 | N/A | No assembly at all concentrations (Blue Class) | 1 |
| GAL11 | 4 | N/A | No assembly at all concentrations (Blue Class) | 1 |
| JSN1 | 1 | 3.14041 | Continuous Transition (Red Class) | 0.00444 |
| JSN1 | 2 | 3.06694 | Incomplete Transition (Yellow Class) | 0.17727 |
| JSN1 | 3 | 3.10638 | Incomplete Transition (Yellow Class) | 0.26706 |
| JSN1 | 4 | 3.11342 | Incomplete Transition (Yellow Class) | 0.34124 |
| SLA2 | 1 | N/A | Assembled at all concentrations (Black Class) | 1 |
| SLA2 | 2 | 2.08704 | Incomplete Transition (Yellow Class) | 0.02772 |
| SLA2 | 3 | 1.59391 | Continuous Transition (Red Class) | 0.06914 |
| SLA2 | 4 | 2.73825 | No assembly at all concentrations (Blue Class) | 1 |
| YLR177W | 1 | N/A | No assembly at all concentrations (Blue Class) | 1 |
| YLR177W | 2 | N/A | No assembly at all concentrations (Blue Class) | 0.80456 |
| YLR177W | 3 | N/A | No assembly at all concentrations (Blue Class) | 0.99989 |
| YLR177W | 4 | N/A | No assembly at all concentrations (Blue Class) | 1 |
| WWM1 | 1 | N/A | No assembly at all concentrations (Blue Class) | 1 |
| WWM1 | 2 | N/A | No assembly at all concentrations (Blue Class) | 0.924 |
| WWM1 | 3 | N/A | No assembly at all concentrations (Blue Class) | 1 |
| WWM1 | 4 | N/A | No assembly at all concentrations (Blue Class) | 1 |
| SIN3 | 1 | 3.96607 | No assembly at all concentrations (Blue Class) | 1 |
| SIN3 | 2 | N/A | No assembly at all concentrations (Blue Class) | 1 |
| SIN3 | 3 | N/A | No assembly at all concentrations (Blue Class) | 0.62964 |
| SIN3 | 4 | N/A | No assembly at all concentrations (Blue Class) | 0.98578 |

| Gene | Replicate | log10(c50) | Classification | Confidence-Score |
|---|---|---|---|---|
| CAF40 | 1 | N/A | No assembly at all concentrations (Blue Class) | 1 |
| CAF40 | 2 | N/A | No assembly at all concentrations (Blue Class) | 1 |
| CAF40 | 3 | N/A | No assembly at all concentrations (Blue Class) | 0.99927 |
| CAF40 | 4 | N/A | No assembly at all concentrations (Blue Class) | 1 |
| LSM4 | 1 | 2.88497 | Continuous Transition (Red Class) | 0.16361 |
| LSM4 | 2 | 2.80195 | Continuous Transition (Red Class) | 0.08323 |
| LSM4 | 3 | 2.87527 | Continuous Transition (Red Class) | 0.13479 |
| LSM4 | 4 | 2.92284 | Continuous Transition (Red Class) | 0.15809 |
| HRR25 | 1 | N/A | No assembly at all concentrations (Blue Class) | 1 |
| HRR25 | 2 | N/A | No assembly at all concentrations (Blue Class) | 1 |
| HRR25 | 3 | N/A | No assembly at all concentrations (Blue Class) | 1 |
| HRR25 | 4 | N/A | No assembly at all concentrations (Blue Class) | 1 |
| NRP1 | 1 | 2.7444 | Discontinuous Transition (Green Class) | 0.26692 |
| NRP1 | 2 | 2.65568 | Discontinuous Transition (Green Class) | 0.35043 |
| NRP1 | 3 | 2.60782 | Discontinuous Transition (Green Class) | 0.54497 |
| NRP1 | 4 | 2.69772 | Discontinuous Transition (Green Class) | 0.23052 |
| YBR016W | 1 | 4.31839 | Incomplete Transition (Yellow Class) | 0.97747 |
| YBR016W | 2 | 4.35962 | Incomplete Transition (Yellow Class) | 0.88839 |
| YBR016W | 3 | 4.52583 | Incomplete Transition (Yellow Class) | 0.97685 |
| YBR016W | 4 | 4.78696 | Incomplete Transition (Yellow Class) | 0.60306 |
| CCR4 | 1 | N/A | No assembly at all concentrations (Blue Class) | 0.81097 |
| CCR4 | 2 | N/A | No assembly at all concentrations (Blue Class) | 0.28548 |
| CCR4 | 3 | N/A | No assembly at all concentrations (Blue Class) | 0.95129 |
| CCR4 | 4 | N/A | No assembly at all concentrations (Blue Class) | 0.99248 |
| GPR1 | 1 | 4.15015 | Incomplete Transition (Yellow Class) | 0.76452 |
| GPR1 | 2 | 4.04825 | Incomplete Transition (Yellow Class) | 0.78389 |
| GPR1 | 3 | 3.99001 | Continuous Transition (Red Class) | 0.16833 |
| GPR1 | 4 | 4.08861 | Incomplete Transition (Yellow Class) | 0.76349 |
| ASM4 | 1 | 2.66528 | Discontinuous Transition (Green Class) | 0.43844 |
| ASM4 | 2 | 2.66779 | Discontinuous Transition (Green Class) | 0.32 |
| ASM4 | 3 | 2.56876 | Discontinuous Transition (Green Class) | 0.12556 |
| ASM4 | 4 | 2.69419 | Discontinuous Transition (Green Class) | 0.34481 |

| Gene | Replicate | log10(c50) | Classification | Confidence-Score |
|---|---|---|---|---|
| YCK1 | 1 | N/A | No assembly at all concentrations (Blue Class) | 0.99813 |
| YCK1 | 2 | 4.14573 | No assembly at all concentrations (Blue Class) | 1 |
| YCK1 | 3 | N/A | No assembly at all concentrations (Blue Class) | 1 |
| YCK1 | 4 | 5.56258 | No assembly at all concentrations (Blue Class) | 1 |
| CLA4 | 1 | N/A | No assembly at all concentrations (Blue Class) | 1 |
| CLA4 | 2 | N/A | No assembly at all concentrations (Blue Class) | 1 |
| CLA4 | 3 | N/A | No assembly at all concentrations (Blue Class) | 1 |
| CLA4 | 4 | N/A | No assembly at all concentrations (Blue Class) | 0.98662 |
| TIF4632 | 1 | 3.4569 | No assembly at all concentrations (Blue Class) | 1 |
| TIF4632 | 2 | 3.62353 | No assembly at all concentrations (Blue Class) | 1 |
| TIF4632 | 3 | 5.41175 | No assembly at all concentrations (Blue Class) | 1 |
| TIF4632 | 4 | 3.99629 | No assembly at all concentrations (Blue Class) | 1 |
| MCA1 | 1 | 3.49122 | Continuous Transition (Red Class) | 0.56947 |
| MCA1 | 2 | 3.24386 | Continuous Transition (Red Class) | 0.46906 |
| MCA1 | 3 | 3.6103 | Continuous Transition (Red Class) | 0.33321 |
| MCA1 | 4 | 3.43038 | Continuous Transition (Red Class) | 0.65242 |
| POP2 | 1 | N/A | No assembly at all concentrations (Blue Class) | 0.98089 |
| POP2 | 2 | N/A | No assembly at all concentrations (Blue Class) | 1 |
| POP2 | 3 | 3.69219 | No assembly at all concentrations (Blue Class) | 1 |
| POP2 | 4 | N/A | No assembly at all concentrations (Blue Class) | 1 |
| ENT2 | 1 | 4.03786 | Incomplete Transition (Yellow Class) | 0.98559 |
| ENT2 | 2 | 3.90737 | Incomplete Transition (Yellow Class) | 0.94646 |
| ENT2 | 3 | 3.92842 | Incomplete Transition (Yellow Class) | 0.98292 |
| ENT2 | 4 | 4.31815 | Incomplete Transition (Yellow Class) | 0.75218 |
| ENT1 | 1 | 3.72344 | Incomplete Transition (Yellow Class) | 0.49747 |
| ENT1 | 2 | 3.65129 | Incomplete Transition (Yellow Class) | 0.41638 |
| ENT1 | 3 | 3.81596 | Incomplete Transition (Yellow Class) | 0.71046 |
| ENT1 | 4 | 3.88488 | Incomplete Transition (Yellow Class) | 0.78601 |
| PUF2 | 1 | N/A | Assembled at all concentrations (Black Class) | 0.81628 |
| PUF2 | 2 | 1.02905 | Assembled at all concentrations (Black Class) | 1 |
| PUF2 | 3 | 1.73897 | Continuous Transition (Red Class) | 0.29297 |
| PUF2 | 4 | N/A | Assembled at all concentrations (Black Class) | 1 |

| Gene | Replicate | log10(c50) | Classification | Confidence-Score |
|---|---|---|---|---|
| NAB2 | 1 | N/A | No assembly at all concentrations (Blue Class) | 0.49827 |
| NAB2 | 2 | N/A | No assembly at all concentrations (Blue Class) | 1 |
| NAB2 | 3 | N/A | No assembly at all concentrations (Blue Class) | 0.9986 |
| NAB2 | 4 | N/A | No assembly at all concentrations (Blue Class) | 1 |
| SKG3 | 1 | N/A | No assembly at all concentrations (Blue Class) | 1 |
| SKG3 | 2 | N/A | No assembly at all concentrations (Blue Class) | 1 |
| SKG3 | 3 | N/A | No assembly at all concentrations (Blue Class) | 1 |
| SKG3 | 4 | N/A | No assembly at all concentrations (Blue Class) | 1 |
| UPC2 | 1 | 3.63474 | Incomplete Transition (Yellow Class) | 0.98373 |
| UPC2 | 2 | 3.52171 | Incomplete Transition (Yellow Class) | 0.98987 |
| UPC2 | 3 | 2.7294 | Continuous Transition (Red Class) | 0.01122 |
| UPC2 | 4 | 2.69679 | Discontinuous Transition (Green Class) | 0.05187 |
| EPO1 | 1 | N/A | No assembly at all concentrations (Blue Class) | 1 |
| EPO1 | 2 | N/A | No assembly at all concentrations (Blue Class) | 0.99271 |
| EPO1 | 3 | N/A | No assembly at all concentrations (Blue Class) | 1 |
| EPO1 | 4 | N/A | No assembly at all concentrations (Blue Class) | 1 |
| SSD1 | 1 | 3.78296 | Incomplete Transition (Yellow Class) | 0.99272 |
| SSD1 | 2 | 3.78232 | No assembly at all concentrations (Blue Class) | 1 |
| SSD1 | 3 | 4.13132 | No assembly at all concentrations (Blue Class) | 1 |
| SSD1 | 4 | 3.98893 | No assembly at all concentrations (Blue Class) | 1 |
| NGR1 | 1 | 2.75225 | Discontinuous Transition (Green Class) | 0.09884 |
| NGR1 | 2 | 2.63139 | Continuous Transition (Red Class) | 0.04355 |
| NGR1 | 3 | 2.58948 | Discontinuous Transition (Green Class) | 0.13361 |
| NGR1 | 4 | 2.74044 | Discontinuous Transition (Green Class) | 0.079 |
| PGD1 | 1 | 3.6516 | Incomplete Transition (Yellow Class) | 0.94968 |
| PGD1 | 2 | 3.6636 | Incomplete Transition (Yellow Class) | 0.94764 |
| PGD1 | 3 | 3.72017 | Incomplete Transition (Yellow Class) | 0.97784 |
| PGD1 | 4 | 3.70865 | Incomplete Transition (Yellow Class) | 0.96308 |
| SLM1 | 1 | N/A | No assembly at all concentrations (Blue Class) | 0.98933 |
| SLM1 | 2 | N/A | No assembly at all concentrations (Blue Class) | 0.92972 |
| SLM1 | 3 | N/A | No assembly at all concentrations (Blue Class) | 0.75374 |
| SLM1 | 4 | N/A | No assembly at all concentrations (Blue Class) | 0.99123 |

| Gene | Replicate | log10(c50) | Classification | Confidence-Score |
|---|---|---|---|---|
| SDD4 | 1 | 3.16676 | Continuous Transition (Red Class) | 0.11301 |
| SDD4 | 2 | 3.08165 | Continuous Transition (Red Class) | 0.0967 |
| SDD4 | 3 | 3.22497 | Continuous Transition (Red Class) | 0.23309 |
| SDD4 | 4 | 3.22042 | Discontinuous Transition (Green Class) | 0.01282 |
| AKL1 | 1 | N/A | No assembly at all concentrations (Blue Class) | 0.99983 |
| AKL1 | 2 | N/A | No assembly at all concentrations (Blue Class) | 1 |
| AKL1 | 3 | N/A | No assembly at all concentrations (Blue Class) | 1 |
| AKL1 | 4 | N/A | No assembly at all concentrations (Blue Class) | 0.85309 |
| RBS1 | 1 | 1.82975 | Continuous Transition (Red Class) | 0.17792 |
| RBS1 | 2 | 2.05539 | Continuous Transition (Red Class) | 0.72572 |
| RBS1 | 3 | 2.03963 | Continuous Transition (Red Class) | 0.96385 |
| RBS1 | 4 | 1.91689 | Continuous Transition (Red Class) | 0.90912 |
| MCM1 | 1 | 5.30619 | No assembly at all concentrations (Blue Class) | 1 |
| MCM1 | 2 | N/A | No assembly at all concentrations (Blue Class) | 1 |
| MCM1 | 3 | N/A | No assembly at all concentrations (Blue Class) | 0.65426 |
| MCM1 | 4 | N/A | No assembly at all concentrations (Blue Class) | 0.74376 |
| NSP1 | 1 | 2.9803 | Discontinuous Transition (Green Class) | 0.7901 |
| NSP1 | 2 | 2.94259 | Discontinuous Transition (Green Class) | 0.72427 |
| NSP1 | 3 | 2.97746 | Discontinuous Transition (Green Class) | 0.74131 |
| NSP1 | 4 | 2.9443 | Discontinuous Transition (Green Class) | 0.74389 |
| VTS1 | 1 | N/A | No assembly at all concentrations (Blue Class) | 0.75444 |
| VTS1 | 2 | N/A | No assembly at all concentrations (Blue Class) | 0.74521 |
| VTS1 | 3 | N/A | No assembly at all concentrations (Blue Class) | 0.39748 |
| VTS1 | 4 | 5.27819 | No assembly at all concentrations (Blue Class) | 1 |
| YBL081W | 1 | 2.35525 | Discontinuous Transition (Green Class) | 0.12068 |
| YBL081W | 2 | 2.2571 | Discontinuous Transition (Green Class) | 0.19335 |
| YBL081W | 3 | 2.29278 | Discontinuous Transition (Green Class) | 0.44009 |
| YBL081W | 4 | 2.24925 | Discontinuous Transition (Green Class) | 0.26212 |
| NUP42 | 1 | N/A | No assembly at all concentrations (Blue Class) | 0.97464 |
| NUP42 | 2 | 4.37295 | No assembly at all concentrations (Blue Class) | 1 |
| NUP42 | 3 | 4.30003 | No assembly at all concentrations (Blue Class) | 1 |
| NUP42 | 4 | N/A | No assembly at all concentrations (Blue Class) | 0.33485 |

| Gene | Replicate | log10(c50) | Classification | Confidence-Score |
|---|---|---|---|---|
| SNF2 | 1 | N/A | No assembly at all concentrations (Blue Class) | 0.99699 |
| SNF2 | 2 | N/A | No assembly at all concentrations (Blue Class) | 1 |
| SNF2 | 3 | N/A | No assembly at all concentrations (Blue Class) | 1 |
| SNF2 | 4 | N/A | No assembly at all concentrations (Blue Class) | 1 |
| HRP1 | 1 | 3.03281 | Continuous Transition (Red Class) | 0.23741 |
| HRP1 | 2 | 2.93458 | Continuous Transition (Red Class) | 0.04084 |
| HRP1 | 3 | 2.95435 | Continuous Transition (Red Class) | 0.22321 |
| HRP1 | 4 | 3.02647 | Continuous Transition (Red Class) | 0.12591 |
| SGF73 | 1 | N/A | No assembly at all concentrations (Blue Class) | 0.98477 |
| SGF73 | 2 | 4.64626 | No assembly at all concentrations (Blue Class) | 1 |
| SGF73 | 3 | 4.32142 | No assembly at all concentrations (Blue Class) | 1 |
| SGF73 | 4 | N/A | No assembly at all concentrations (Blue Class) | 0.9778 |
| NAB3 | 1 | 1.13884 | Assembled at all concentrations (Black Class) | 1 |
| NAB3 | 2 | 1.19238 | Assembled at all concentrations (Black Class) | 1 |
| NAB3 | 3 | N/A | Assembled at all concentrations (Black Class) | 0.70136 |
| NAB3 | 4 | 1.07915 | Assembled at all concentrations (Black Class) | 1 |
| PAN1 | 1 | 2.70573 | Continuous Transition (Red Class) | 0.00028 |
| PAN1 | 2 | 2.72818 | Discontinuous Transition (Green Class) | 0.05652 |
| PAN1 | 3 | 2.64859 | Discontinuous Transition (Green Class) | 0.11667 |
| PAN1 | 4 | 2.74148 | Discontinuous Transition (Green Class) | 0.00485 |
| CYC8 | 1 | 3.08171 | Discontinuous Transition (Green Class) | 0.28287 |
| CYC8 | 2 | 3.00557 | Discontinuous Transition (Green Class) | 0.28361 |
| CYC8 | 3 | 2.96179 | Discontinuous Transition (Green Class) | 0.42775 |
| CYC8 | 4 | 3.18428 | Discontinuous Transition (Green Class) | 0.34745 |
| NUP49 | 1 | 3.35552 | Incomplete Transition (Yellow Class) | 0.60793 |
| NUP49 | 2 | 3.32693 | Incomplete Transition (Yellow Class) | 0.42966 |
| NUP49 | 3 | 3.36907 | Incomplete Transition (Yellow Class) | 0.49218 |
| NUP49 | 4 | 3.33511 | Incomplete Transition (Yellow Class) | 0.48106 |
| CBK1 | 1 | 3.61579 | Incomplete Transition (Yellow Class) | 0.93827 |
| CBK1 | 2 | 3.94991 | No assembly at all concentrations (Blue Class) | 1 |
| CBK1 | 3 | 5.39718 | No assembly at all concentrations (Blue Class) | 1 |
| CBK1 | 4 | 4.24137 | No assembly at all concentrations (Blue Class) | 1 |

| Gene | Replicate | log10(c50) | Classification | Confidence-Score |
|---|---|---|---|---|
| PSP1 | 1 | 2.7653 | Discontinuous Transition (Green Class) | 0.72513 |
| PSP1 | 2 | 2.98055 | Continuous Transition (Red Class) | 0.00252 |
| PSP1 | 3 | 2.67973 | Discontinuous Transition (Green Class) | 0.69536 |
| PSP1 | 4 | 2.72202 | Discontinuous Transition (Green Class) | 0.67023 |
| AZF1 | 1 | 3.80441 | Incomplete Transition (Yellow Class) | 0.99503 |
| AZF1 | 2 | 3.74481 | Incomplete Transition (Yellow Class) | 0.99488 |
| AZF1 | 3 | 3.81324 | No assembly at all concentrations (Blue Class) | 1 |
| AZF1 | 4 | 3.80368 | No assembly at all concentrations (Blue Class) | 1 |
| YAP1801 | 1 | 2.2158 | Discontinuous Transition (Green Class) | 0.19507 |
| YAP1801 | 2 | 1.98637 | Discontinuous Transition (Green Class) | 0.09256 |
| YAP1801 | 3 | 3.63809 | No assembly at all concentrations (Blue Class) | 1 |
| YAP1801 | 4 | 1.80598 | Discontinuous Transition (Green Class) | 0.24605 |
| NUP57 | 1 | 3.94703 | No assembly at all concentrations (Blue Class) | 1 |
| NUP57 | 2 | 3.70453 | Incomplete Transition (Yellow Class) | 0.99302 |
| NUP57 | 3 | 4.58337 | No assembly at all concentrations (Blue Class) | 1 |
| NUP57 | 4 | 3.81348 | Incomplete Transition (Yellow Class) | 0.9975 |
| SNF5 | 1 | N/A | No assembly at all concentrations (Blue Class) | 1 |
| SNF5 | 2 | N/A | No assembly at all concentrations (Blue Class) | 1 |
| SNF5 | 3 | N/A | No assembly at all concentrations (Blue Class) | 0.95201 |
| SNF5 | 4 | N/A | No assembly at all concentrations (Blue Class) | 1 |
| SOK2 | 1 | 2.46392 | Continuous Transition (Red Class) | 0.19211 |
| SOK2 | 2 | 2.5083 | Continuous Transition (Red Class) | 0.1593 |
| SOK2 | 3 | 2.48824 | Continuous Transition (Red Class) | 0.08865 |
| SOK2 | 4 | 2.4494 | Continuous Transition (Red Class) | 0.15341 |
| AIM3 | 1 | 2.90383 | Discontinuous Transition (Green Class) | 0.18089 |
| AIM3 | 2 | 2.60868 | Discontinuous Transition (Green Class) | 0.28213 |
| AIM3 | 3 | 2.82519 | Discontinuous Transition (Green Class) | 0.28513 |
| AIM3 | 4 | 2.83215 | Discontinuous Transition (Green Class) | 0.16509 |
| SCD5 | 1 | 2.65369 | Continuous Transition (Red Class) | 0.02031 |
| SCD5 | 2 | 2.72875 | Continuous Transition (Red Class) | 0.09531 |
| SCD5 | 3 | 2.65141 | Continuous Transition (Red Class) | 0.05025 |
| SCD5 | 4 | 2.68091 | Continuous Transition (Red Class) | 0.15817 |

| Gene | Replicate | log10(c50) | Classification | Confidence-Score |
|---|---|---|---|---|
| DEF1 | 1 | 2.80259 | Continuous Transition (Red Class) | 0.26867 |
| DEF1 | 2 | 2.36541 | Continuous Transition (Red Class) | 0.46392 |
| DEF1 | 3 | 2.28945 | Continuous Transition (Red Class) | 0.3407 |
| DEF1 | 4 | 2.31908 | Continuous Transition (Red Class) | 0.4924 |
| SWI1 | 1 | 3.26805 | Discontinuous Transition (Green Class) | 0.12983 |
| SWI1 | 2 | 3.15395 | Discontinuous Transition (Green Class) | 0.28509 |
| SWI1 | 3 | 3.16916 | Discontinuous Transition (Green Class) | 0.577 |
| SWI1 | 4 | 3.27442 | Discontinuous Transition (Green Class) | 0.28426 |
| NUP116 | 1 | 1 | Assembled at all concentrations (Black Class) | 1 |
| NUP116 | 2 | 1.89005 | Discontinuous Transition (Green Class) | 0.40577 |
| NUP116 | 3 | 1.90558 | Discontinuous Transition (Green Class) | 0.54484 |
| NUP116 | 4 | 1.58415 | Higher Order State (Magenta Class) | 0.41807 |
| NUP100 | 1 | N/A | Assembled at all concentrations (Black Class) | 0.43223 |
| NUP100 | 2 | 1.42113 | Assembled at all concentrations (Black Class) | 1 |
| NUP100 | 3 | 1.46024 | Assembled at all concentrations (Black Class) | 1 |
| NUP100 | 4 | N/A | Assembled at all concentrations (Black Class) | 0.33569 |

**Table S3: Description of the titles in Figure S2 that are used to explain how each synthetic dataset was generated.** Further descriptions of how the datasets were generated are described in the main text and the Materials and Methods section. Noise Profile 1 implies low noise, Noise Profile 2 implies medium noise, and Noise Profile 3 implies high noise.

| Column Number | Column Value | Description |
|---|---|---|
| 1 | X | Data generated using a sigmoidal function. |
| 1 | Y | Data generated using a step function. |
| 1 | Z | 3-state data generated with overlapping regions. |
| 2 - 3 | NN | Location of discontinuity in log10(c50) concentration. |
| 4 | L | Low concentration type (points added below the saturation concentration) |
| 4 | H | High concentration type (points added above the saturation concentration) |
| 4 | U | Uniform concentration type (points added uniformly) |
| 4 | V | Uniform Sparse concentration type (fewer points added uniformly) |
| 5 | A | Noise Profile 1 |
| 5 | B | Noise Profile 2 |
| 5 | C | Noise Profile 3 |
| 6 | 1 | Replicate 1 |
| 6 | 2 | Replicate 2 |
| 6 | 3 | Replicate 3 |

**Table S4: Plasmid numbers, cell counts, and amino acid sequences of each of the cPrDs studied.**
